# Perturbative formulation of general continuous-time Markov model of sequence evolution via insertions/deletions, Part I: Theoretical basis

**DOI:** 10.1101/023598

**Authors:** Kiyoshi Ezawa, Dan Graur, Giddy Landan

## Abstract

**Background:** Insertions and deletions (indels) account for more nucleotide differences between two related DNA sequences than substitutions do, and thus it is imperative to develop a stochastic evolutionary model that enables us to reliably calculate the probability of the sequence evolution through indel processes. Recently, such probabilistic models are mostly based on either hidden Markov models (HMMs) or transducer theories, both of which give the indel component of the probability of a given sequence alignment as a product of either probabilities of column-to-column transitions or block-wise contributions along the alignment. However, it is not *a priori* clear how these models are related with any *genuine* stochastic evolutionary model, which describes the stochastic evolution of an *entire* sequence along the time-axis. Moreover, none of these models can fully accommodate biologically realistic features, such as overlapping indels, power-law indel-length distributions, and indel rate variation across regions.

**Results:** Here, we theoretically tackle the *ab initio* calculation of the probability of a given sequence alignment under a *genuine* evolutionary model, more specifically, a general continuous-time Markov model of the evolution of an *entire* sequence via insertions and deletions. Our model allows general indel rate parameters including length distributions but does not impose any unrealistic restrictions on indels. Using techniques of the perturbation theory in physics, we expand the probability into a series over different numbers of indels. Our derivation of this perturbation expansion elegantly bridges the gap between Gillespie’s (1977) intuitive derivation of his own stochastic simulation method, which is now widely used in evolutionary simulators, and Feller’s (1940) mathematically rigorous theorems that underpin Gillespie′s method. We find a sufficient and nearly necessary set of conditions under which the probability can be expressed as the product of an overall factor and the contributions from regions separated by gapless columns of the alignment. The indel models satisfying these conditions include those with some kind of rate variation across regions, as well as space-homogeneous models. We also prove that, though with a caveat, pairwise probabilities calculated by the method of Miklós et al. (2004) are equivalent to those calculated by our *ab initio* formulation, at least under a space-homogenous model.

**Conclusions:** Our ab initio perturbative formulation provides a firm theoretical ground that other indel models can rest on.

[This paper and three other papers (Ezawa, Graur and Landan 2015a,b,c) describe a series of our efforts to develop, apply, and extend the *ab initio* perturbative formulation of a general continuous-time Markov model of indels.]

## Introduction

### Background

The evolution of biomolecules, namely DNA, RNA, and protein sequences, is driven by mutations such as base substitutions, insertions and deletions (indels), recombination, and other genomic rearrangements (*e.g.*, Graur and Li 2000; Gascuel 2005; Lynch 2007). Among them, substitutions and indels have been considered particularly important because they are modeled, either implicitly or explicitly, in the algorithms for the sequence alignments, which have played central roles in the sequence analysis in bioinformatics (*e.g.*, Gusfield 1997; Notredame 2007). Probably due to the relative ease in handling them, analyses on substitutions have predominated in the field of molecular evolutionary study thus far, in particular using the probabilistic (or likelihood) theory of substitutions that is now widely accepted (e.g., Felsenstein 1981, 2004; Yang 2006). It should not be forgotten, however, that some recent comparative genomic analyses have revealed that indels account for more base differences between the genomes of closely related species than substitutions (*e.g.*, Britten 2002; Britten *et al*. 2003; Kent *et al*. 2003; The International Chimpanzee Chromosome 22 Consortium 2004; The Chimpanzee Sequencing and Analysis Consortium 2005). It is therefore imperative to develop a stochastic model that enables us to reliably calculate the probability of sequence evolution via mutations including insertions and deletions.

As far as we know, the development of probabilistic theories of indels dates back to the groundbreaking work of Bishop and Thompson (1986), where they obtained the most likely (ML) pairwise alignment (PWA) under a simple stochastic model of single-base indels and substitutions. Then, in their pioneering work, Thorne, Kishino and Felsenstein (1991) presented a simple yet more refined stochastic model of sequence evolution, often called the TKF91 model, which evolves a DNA sequence via substitutions, insertions and deletions, all of single bases. Using this TKF91 model, they worked out the ML alignment, as well as the summation of probabilities over all possible alignments, between two homologous sequences. And they used the latter to reliably estimate the model parameters. An obvious drawback of this model is that they incorporate only single-base indels, whereas indels of multiple contiguous bases have been known to occur frequently by experiments. This drawback is somewhat mitigated by their subsequent model, the TKF92 model (Thorne et al. 1992), which allowed for a geometric indel length distribution, but which imposed an unrealistic restriction that indels can occur only in the unit of unbreakable fragments. Such efforts to “inch toward reality” were taken over by some researchers, resulting in a few biologically more realistic models and algorithms (*e.g.*, Miklós and Toroczkai 2001; Knudsen and Miyamoto 2003; Miklós et al. 2004; Kim and Sinha 2007). (See below for more details on the biological realism.) The use of probabilistic models of indels seems to have expanded as the 21st century began, since the TKF91 model was recast into a hidden Markov model (HMM) (Hein 2001; Holmes and Bruno 2001) and a transducer theory (Holmes 2003), because these models facilitates the constructions of the dynamic programming (DP) to search for the ML alignment and of the DP to sum probabilities over possible alignments. For example, the statistical alignment algorithms were immediately extended from a sequence pair to multiple sequences (Hein 2001; Holmes and Bruno 2001; Holmes 2003), and their time complexity was substantially reduced (*e.g.*, Lunter et al. 2003). Regarding the Markov chain Monte Carlo (MCMC) methods to simultaneously sample multiple sequence alignments, phylogenetic trees, and model parameters (Holmes and Bruno 2001), considerabl efforts were made to speed up the algorithm and to accelerate the convergence of the MCMC trajectories (Lunter et al. 2005; Redelings and Suchard 2005, 2007; Suchard and Redelings 2006; Novak et al. 2008). Then, the HMMs and transducer theories to describe indels were extended to accommodate a general geometric distribution of indel lengths, either based on the TKF92 model (Thorne et al. 1992; Metzler 2003), by taking account of some evolutionary effects on gap patterns (Knudsen and Miyamoto 2003; Miklós et al. 2004; Rivas 2005), or by simply applying standard HMMs/transducers or their modifications (*e.g.*, Löytynoja and Goldman 2005; Redelings and Suchard 2007; Lunter et al. 2008; Paten et al. 2008). Such models with a geometric indel length distribution were then applied to the algorithms to reconstruct the multiple sequence alignment (MSA), which either search for a single optimum MSA (Do et al. 2005; Löytynoja and Goldman 2005, 2008; Löytynoja et al. 2012) or sample a number of fairly likely MSAs (Paten et al. 2008; Westesson et al. 2012). These indel probabilistic models were also used in some algorithms to reconstruct ancestral sequences from an input MSA, either by using the input MSA as it is (Diallo et al. 2007, 2010), while locally improving the alignment via a ML criterion (Kim and Sinha 2007), or while taking account of alignment uncertainties (Paten et al. 2008; Westesson et al. 2012). The models were also used for the secondary structure prediction of protein sequences (Miklós et al. 2008). And some very recent studies further strengthened the mathematical bases of HMMs and/or transducers (*e.g.*, Bouchard-Côté 2013) and achieved further algorithmic efficiencies in representing MSA uncertainties (*e.g.*, Herman et al. 2015). Meanwhile, in order to speed up the alignment estimation and/or the phylogenetic analysis, further simplifications of the TKF91 model were also made, either via an extension of base substitution models to include a gap as a “fifth character” (McGuire et al. 2001; Rivas 2005; Rivas and Eddy 2008), or via an approximation by a model of Poisson indel processes (Bouchard-Côté and Jordan 2013). See excellent reviews (e.g., Rivas 2005; Bradley and Holmes 2007; Miklós et al. 2009) for more details on the recent developments and applications of these indel probabilistic models.

Thus, concerning the algorithmic efficiency and the scope of applications, as well as their *mathematical* bases, the probabilistic models of indels have advanced in many great steps. However, these desirable properties *alone* are not sufficient for a theoretical model to be used *reliably* for the purpose. In addition to the *mathematical* soundness, a reliable model must also approximate well, or at least decently, the *real* phenomena it is intended to describe. In the case of an indel probabilistic model, there are two key properties for this requisite: one is the *evolutionary consistency*, and the other is the flexibility to accommodate various biologically realistic features, *i.e.*, the *biological realism*, of indels. Let us first explain the evolutionary consistency. From the theoretical viewpoint, there should be no argument about the idea that a *genuine* stochastic model of sequence evolution via indels must be the one that describes the evolution of the *entire* sequence in question along the time axis (or down a lineage or a branch). The probability calculation under such a genuine evolutionary model must naturally proceed via the multiplicative accumulation of the probabilities of *vertical* transitions, each from the state *of the entire sequence* at a time to its state at the next time (separated either infinitesimally or by a finite but small interval). In contrast, standard HMMs and transducer theories calculate the indel component of the probability of an alignment as the product of the probabilities of *horizontal* transitions, each from the state of a column to that of the next column. Although more general forms of HMMs and transducers also exist, they still calculate the probability *horizontally* as the product of block-wise contributions (*e.g.*, Miklós et al. 2004; Kim and Sinha 2007). Therefore, it is *a priori* not clear whether or not the HMMs or transducer theories are related to any *genuine* evolutionary models, and, if they are, how. It should be worth a mention that some HMMs and transducer theories were actually derived from the exact solutions of “genuine” evolutionary models, such as the TKF91 and TKF92 models and the model proposed by Miklós and Toroczkai (2001). It must be noted, however, that these models were devised so that their exact solutions will give a probability that can be trivially factorized. In consequence, these models inevitably impose some biologically unrealistic restrictions on the indel events, such as single-base indels (TKF91), indels occurring in the unit of unbreakable fragments (TKF92), and single-base deletions while permitting breakable multiple-base insertions (Miklós and Toroczkai 2001). To the best of our knowledge, no studies thus far *explicitly* showed that the indel probability calculated under a *genuine* and *biologically realistic* evolutionary model can be expressed as the product of either column-wise or block-wise contributions. Nevertheless, it should also be worth a mention that a few attempts were made to relate genuine evolutionary models with HMMs/transducers. Knudsen and Miyamoto (2003) started with the assumption that the probability is given by the product of column-to-column transition probabilities. Then they determined the explicit forms of the transition probabilities by taking account of an evolutionary indel model. Unfortunately, the resulting model was similar to a standard HMM, in the sense that it could not incorporate the full effects of general overlapping indels and that it could only implement geometric indel length distributions. In what could be called a milestone study, Miklós et al. (2004) proposed a “long indel” model, which can take account of overlapping indels up to the level desired by users (at least in principle). In this study, they conjectured that the probability of a given pairwise alignment can be calculated as a product of contributions from “chop zone’s each of which is delimited by neighboring gapless columns. Then the contribution from each chop zone was calculated up to a user-specified number of overlapping indel events according to a continuous-time Markov model. Unfortunately, although they *conceptually* started with a genuine evolutionary model, *i.e.*, a continuous-time Markov model of an *entire* sequence evolution (called the “SID model,” see below), they did not *explicitly* show through equations that their conjectured probability is indeed related in any way to the *ab initio* probabilities derived from the genuine model. Although their verbal justification was plausible and will turn out to be correct in this study (Appendix A6), it was unclear in their study to what extent conditions on the indel rate parameters can be relaxed while keeping the probability factorable. For a further, sound advance of the study of molecular evolution via indels, it is essential to resolve these outstanding issues. At the same time, it would also be important to examine the parameter regions where the HMMs/transducers can well approximate the probability that was calculated by a genuine and biologically realistic evolutionary model. Such analyses would reveal up to how far we can trust the models that we use or develop. One of the most frequently cited problems of most HMMs/transducers, except those by Miklós and Toroczkai (2001) and by Miklós et al. (2004), is that these models cannot accommodate overlapping indels, which makes the models to violate the multiplicativity condition, aka the Chapman-Kolmogorov equation (*e.g.*, Westesson et al. 2012). Some simulation analyses seemed to show that this problem does not usually impact the results of the analyses significantly (*e.g.*, Thorne et al. 1992; Knudsen and Miyamoto 2003; Metzler 2003). However, their analyses depended on models with geometric indel length distributions, and there is no guarantee that their conclusions remain valid even with more realistic distributions. It is thus crucial to exactly delimit the parameter regions where the effects of overlapping indels are indeed negligible. This would require either analytical expressions of the probabilities under a genuine evolutionary model or a more systematic numerical or simulation study.

Regarding the biological realism, it should be mentioned first that real, biological indel lengths were frequently shown to follow the power-law distributions (*e.g.*, Gonnet et al. 1992; Benner et al. 1993; Gu and Li 1995; Kent et al. 2003; Zhang and Gerstein 2003; Chang and Benner 2004; Yamane et al. 2006; Fan et al. 2007), at least up to several kilobases (The international Chimpanzee Chromosome 22 Consortium 2004). Moreover, it is widely believed that the indel rates should vary among regions, due to selection and the mutational predispositions of the regions themselves (caused, *e.g.*, by their sequence or epigenomic contexts) (*e.g.*, Gu et al. 2008). On the contrary, normal HMMs and transducer theories can at best handle geometric distributions of indel lengths, which behave very differently from the power-law. For example, under a normal geometric distribution, long indels get much rarer than empirically observed. Although the problem could be somewhat mitigated by extending HMMs or transducers to allow for mixed geometric distributions (*e.g.*, Miklós et al. 2004; Lunter et al. 2008), it is still difficult to reproduce the observed frequency of indels that are, *e.g.*, as long as hundreds of bases. From the viewpoint of the biological realism, works by Miklós et al. (2004) and Kim and Sinha (2007) are notable. The “long indel” model of Miklós et al. (2004) can in principle handle any indel length distributions that are uniform across the sequence (except for indels reaching either end), as long as the insertion length distribution and the deletion length distribution depend on each other via the time reversibility condition (aka the detailed balance condition). As Miklós et al. themselves noted, they imposed the time reversibility just for the technical convenience of simplifying the calculation of the probabilities of pairwise alignments. As they rightly argued, however, there is no biological reason to expect that realistic indel models must satisfy the time reversibility (see also Rivas and Eddy 2008). Although the condition is not necessarily essential for the model, imposing it can restrict the indel length distributions awkwardly. Another problem of the “long indel” model would be that it is space-homogeneous, and that it was unclear how this HMM can be extended to accommodate space-heterogeneity. The model of Kim and Sinha (2007) is even more flexible. Their model is a kind of HMM that calculates the probability of a given multiple sequence alignment (MSA) as a product of contributions from gapless and gapped blocks. Thus, it can accommodate any functional forms of insertion and deletion length distributions in principle. And, because their model does not impose the time reversibility, the two length distributions can be independent of each other. Their model, however, has a major problem, as most other HMMs and transducer theories do. That is, their model is not exactly evolutionarily consistent. In consequence, their model cannot accommodate overlapping indels along a single branch, though it can handle overlapping indels that occurred along different branches.

Meanwhile, some researchers developed *genuine* molecular evolution simulators, such as Dawg (Cartwright 2005), INDELible (Fletcher and Yang 2009), and indel-Seq-Gen version 2.0 (Strope et al. 2009). They can simulate the evolution of an *entire* sequence along the time axis or down a phylogenetic tree, under a fairly biologically realistic model of indels that allow for both overlapping indels and a flexible setting of rate parameters and length distributions, including the power-law distributions. Thus, if we want, we could examine problems concerning, *e.g.*, the principles of evolutionary models by performing systematic, computer-intensive analyses via one of these genuine molecular evolution simulators. Nevertheless, it should be definitely more desirable if we have a theoretical formulation that can somehow, analytically or numerically, calculate the *ab initio* probabilities of indel processes under a *genuine* and biologically realistic evolutionary model.

Thus far, theoretical studies on molecular evolution seem to have been obsessed with exact solutions, whether analytical or numerical. This is partly because exact solutions were successfully obtained for the continuous-time Markov models of substitutions of nucleotides and amino acids (see, *e.g.*, Yang 2006). However, exact solutions are not necessarily a must-have for a scientific field to develop successfully. As a case in point, let us briefly review the elementary particle physics, one of the most successful disciplines of natural science in the 20th century. Its standard model consists of two major components: the electroweak theory describing the electromagnetic and weak interactions (Glashow 1961; Weinberg 1967; Salam 1968), and the quantum chromodynamics (QCD) describing the strong interactions (*e.g.*, Gross and Wilczek 1973; Politzer 1974). To the best of our knowledge, these models have never been solved exactly. Instead, the success of the particle physics rested heavily on approximations, and among the most important approximation methods is the perturbation theory (*e.g.*, Dirac 1958; Messiah 1961b), in which elementary particles behave as free particles in most of the time and occasionally undergo perturbations due to interactions. The key to its success was the fact that the interaction coefficients were small enough for the interactions to be treated as occasional perturbations. (Although the coefficients of the strong interactions are normally large, they approach zero in the high-energy limit (Gross and Wilczek 1973; Politzer 1974), which enabled the perturbation theory to work.) Getting back to the molecular evolution, recent genome-wide analyses showed that the rate of indels is at most on the order of 1/10 of the substitution rate (Lunter 2007; Cartwright 2009). Thus, as long as we are dealing with sequences that are detectably homologous to each other, the expected number of indels per site along a branch will be well below 1. This gives us a hope that we can calculate the probabilities of indel processes by applying techniques of the perturbation theory in physics (*e.g.*, Dirac 1958; Messiah 1961b).

### About this paper

This paper reports our somewhat successful and absolutely orthodox theoretical attempt to calculate, *from the first principle*, the probability of a given sequence alignment under a *genuine* evolutionary model, more specifically, a continuous-time Markov model on an infinite set of states that describes the evolution of an *entire* sequence along the time axis via insertions and deletions. We calculate the alignment probability under a *fixed* tree topology and branch lengths, and we handle both pairwise alignments (PWAs) and multiple sequence alignments (MSAs). Our continuous-time Markov model allows for general indel rate parameters including indel length distributions, and it does not impose any unrealistic restrictions on the permitted indels. This generalization includes (but is not limited to) allowing the model to be non-time-reversible, as the models of Eddy and Rivas (2008) and Kim and Sinha (2008) did. For clarity, we will focus only on the indel processes in the bulk of the paper, by not explicitly considering substitutions (while implicitly taking account of residue states of the sites of the sequence). However, as will be argued in Part IV (Ezawa, Graur and Landan 2015c), incorporating substitutions would be rather straightforward, as long as the substitution model involved is of a commonly used type. [NOTE: As far as we know, no studies thus far considered a *genuine* stochastic evolutionary model of indels that is this general. But there is only one such model that we know is only slightly less general, which is (the indel components of) the class of “substitution/insertion/deletion (SID) models” equipped with “rate grammars” (Miklós et al. 2004). The most general among them is only devoid of *inherent* heterogeneity across sites, though sites could be heterogeneous due to different residue contexts surrounding them. Unfortunately, this general SID model was only proposed but not theoretically developed further, probably because the substitution and indel events in its general form seemed intricately entangled. Instead, Miklós et al. (2004) theoretically developed its simpler version, *i.e.*, the aforementioned “long indel” model. Well, the general SID model could be regarded as the model presented here, when the latter is defined on the least general state space (*S^I^*, defined in subsection 2.1). Thus, the methods proposed in the current paper, such as the operator representation of indels and the equivalence relations between indel histories, can also be applied to the SID models, and facilitate their theoretical developments. Thus, in a sense, the current study could also be considered as reviving the general SID model.] We start in Section 1 of Results by introducing some convenient concepts from theoretical physics (Dirac 1958; Messiah 1961a). In Section 2 of Results, we formulate the *genuine* indel evolutionary model in terms of the concepts introduced in Section 1. A key innovation is the representation of each indel event as an operator that acts on the state of an *entire* sequence (, which is represented with a bra vector). This enables us to define a new concept, that is, the **“local-history-set” (LHS) equivalence class** of indel histories, which will play an essential role when proving the factorization of an alignment probability. In Section 3 of Results, using techniques of the perturbation theory in physics, we formally expand the probability of an alignment into a series of terms with different numbers of indels, where the fewest-indel terms are contributed by parsimonious indel histories and other terms come from non-parsimonious histories. This perturbation expansion formally proves that the widely used stochastic method of Gillespie (1977) indeed provides the basis of *genuine* evolutionary simulators, and our proof serves as a bridge between its mathematically rigorous proof by Feller (1940) and its intuitive derivation by Gillespie himself. In Section 4 of Results, we find a sufficient and nearly necessary set of conditions on the indel rate parameters and the ancestral sequence state probability under which the alignment probability can be expressed as the product of an overall factor and the contributions from regions separated by gapless columns of the alignment. Here the qualifier, “nearly necessary,” means that there may be some *isolated* cases where the probability can be factorized even if some of the conditions are violated. Nevertheless, even if there are, such cases are likely to require intricate and miraculous cancellations among terms, and thus are unlikely to be important in practical analyses. In Section 5 of Results, we give some example indel models as particular solutions of the conditions derived in Section 4. They include: models with space-homogeneous indel rates including the “long indel” model (Miklós et al. 2004), models with indel rates confined in separate regions, and models with the linear combinations of the above indel rates. In that section, we also show that, when its application is extended to each LHS equivalence class of indel histories during a time interval, the method of Miklós et al. (2004) gives the same probability as our *ab initio* formulation does, at least under a space- and time-homogeneous indel model. In Discussion, we will briefly discuss some possible applications of our theory. The topics also include the risks associated with the naïve application of our algorithm to *reconstructed* alignments. Appendix is devoted to detailed explanations on the proofs and derivations of some key results.

This paper is part I of a series of our papers that documents our efforts to develop, apply, and extend the *ab initio* perturbative formulation of the general continuous-time Markov model of sequence evolution via indels. Part I (this paper) gives the theoretical basis of this entire study. Part II (Ezawa, Graur and Landan 2015a) describes concrete perturbation calculations and examines the applicable ranges of other probabilistic models of indels. Part III (Ezawa, Graur and Landan 2015b) describes our algorithm to calculate the first approximation of the probability of a given MSA and simulation analyses to validate the algorithm. Finally, part IV (Ezawa, Graur and Landan 2015c) discusses how our formulation can incorporate substitutions and other mutations, such as duplications and inversions.

Before going on to the bulk of the manuscript, we explain important terminology and notation. In this paper, the term “an indel process” means a series of successive indel events with both the order and the specific timings specified, and the term “an indel history” means a series of successive indel events with only the order specified. This usage should conform to the common practice in this field. And, throughout this paper, the union symbol, such as in *A* ⋃ *B* and 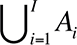, should be regarded as the union of *mutually disjoint* sets (*i.e.*, those satisfying 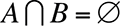 and 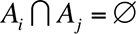 for 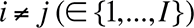, respectively, where Ø is an empty set), unless otherwise stated.

Another important note is on the representation of the continuous-time Markov model. In this series of studies, we employed the bra-ket notation for the sequence states, and represented mutations, such as indels, as operators, borrowing the concepts from the quantum mechanics (Dirac 1958; Messiah 1961a). However, those who are unfamiliar with quantum mechanics need not worry about it. As shown in Section 1, the formulation represented by the bra-ket-operator notation is equivalent to the standard formulation of the continuous-time Markov model represented by the vector-matrix notation, as long as the state space is finite or countably infinite. Therefore, if desired, the symbols of a bra 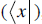, a ket 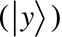, and an operator (*Ô*) could be regarded simply as convenient reminders of a row vector, a column vector, and a matrix, respectively. (It should be kept in mind, however, that the operator notation allows for more flexible representations of state changes, and thus is suitable for the description of indel processes.) Actually, this study has three essential points: (1) shifting the focus from the trajectory of sequence states to the series of indel events when describing an evolutionary process, (2) representing insertions/deletions as operators, and (3) the equivalence relationships between the series of two or more indels. These points enabled the general SID model (Miklós et al. 2004) and its further extension (*i.e.*, our *genuine* stochastic evolutionary model) to move forward and to be theoretically developed further.

## Results

### 1. Preparation: Introduction of bra-ket notation and operators

In this study we examine a continuous-time Markov model defined on a discrete infinite space of states. Although we could still in principle formally construct the theoretical framework in a traditional manner of using the row vectors, matrices, and column vectors, this could get somewhat cumbersome. Thus, instead, we will formulate the theory by using the concepts commonly used in quantum mechanics of physics (*e.g.*, Dirac 1958; Messiah 1961a), namely, the bra-ket notation of the state vectors and the operators. In this section, we introduce these concepts first in a general form and then using an example that most readers may be familiar with, *i.e.*, the continuous-time Markov model of base substitutions. Actually, their usage here is somewhat different from that in quantum mechanics, as will be discussed at the bottom of this section.

#### 1.1. General case

Let us first recall the conventional formulation of a general continuous-time Markov model on a finite space consisting of *N* states, *i* = 1, 2,…,*N*. One way of formulating the model is to specify a rate matrix, *Q* = (*q_ij_*). Let *q_ij_* denote the (*i, j*)- element of *Q, i.e.*, its element at the intersection of the *i* th row and the *j* th column. Then, the non-diagonal element *q_ij_* (*i* ≠ *j*) of a rate matrix *Q* is the rate (per a certain unit time) at which the system moves to the *j* th state, given it was in the *i* th state immediately before the time in question. The diagonal element, *q_ii_*, is usually given by the equation:

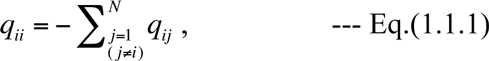

to guarantee that the summation of the probabilities over the states remain 1 all the time. Now, let the probability vector, 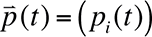, be a row vector whose *i* th element, *p_i_*(*t*), is the probability that the system is in the *i* th state at time *t*. Then, under the above Markov model, 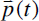 satisfies the 1st order time differential equation:

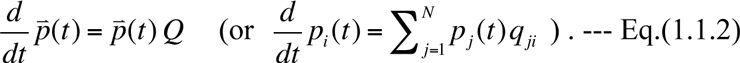

The general solution of this equation at a finite time *t*(> 0) is given by:

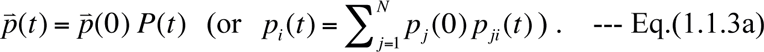

Here the “finite-time stochastic evolution matrix,” 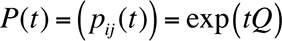, is an *N* × *N* matrix whose (*i, j*)-element *p_ij_* (*t*) is the probability that the system is in the *j* th state at time *t*, conditioned on that it was in the *i* th state initially (*i.e.*, at time *t* = 0):

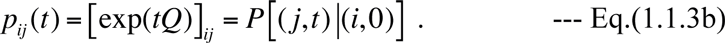

If Eq.(1.1.1) holds, the matrix elements satisfy 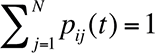 for all *i* = 1, 2,…, *N*. Meanwhile, 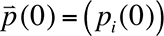 is the initial probability vector, whose *i* th component, *p_i_*(0), is the probability that the system was in the *i* th state at time *t* = 0. They satisfy 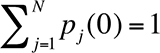. This could be made more explicit by using the basic row vectors, 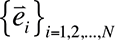. Here 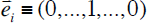 is the row vector with all zeros except the *i* th component, which is 1, and it represents the situation where the system is in the *i* th state. Using these basic vectors, the initial probability vector is expressed as:

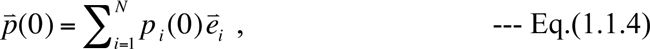

which is interpreted as the initial condition that the system is in the *i* th state with probability *p_i_* (0) (*i* = 1,2,…, *N*). Similarly, the probability vector at any time could be expressed as:

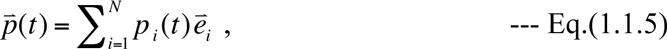

and interpreted as the situation where the system is in the *i* th state with probability *p_i_*(*t*) (*i* = 1, 2,…, *N*) given by Eq.(1.1.3a). Using the basic vectors, the conditional probabilities can be formally extracted from the stochastic evolution matrix, *P*(*t*) = exp(*tQ*), by a matrix multiplication:

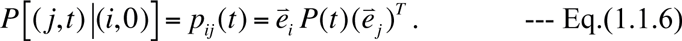

Here 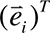 is the column vector obtained from the row vector, 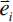, by a matrix transposition operation (*i.e.*, by interchanging the rows with the columns).

Now we can introduce the bra-ket notation and operators. First, we replace each basic row vector, 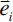, with the corresponding basic bra-vector, 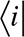, and replace each basic column vector, 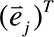, with the corresponding basic ket-vector, 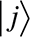. Then, the bra-vector corresponding to the probability vector 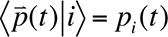 in Eq.(1.1.5) is given by the following linear combination of the basic bra-vectors:

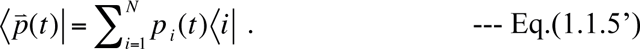

In the present formulation, the exclusive role of a ket-vector is that it serves as an “acceptor” of bra-vectors. More specifically, we will make the ket-vector, 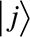, accept only the corresponding bra-vector, 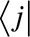, by defining the scalar product:

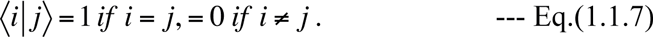

Using these scalar products, we get, *e.g.*, the equation, 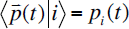, from Eq.(1.1.5’). Next, we introduce (linear) operators that transform each bra-vector into a specified linear combination of bra-vectors. The operators are analogs of matrices in the traditional formulation. For example, we could define an operator, 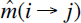, that transforms (or “mutates”) the *i* th state to the *j* th state, but does nothing else:

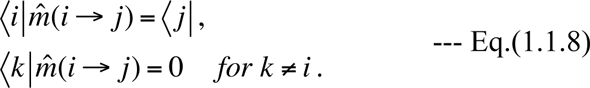

This operator corresponds to the matrix whose elements are all zero except the (*i, j*)–element, which is 1. Now, we define the (instantaneous) rate operator, 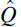, as follows:

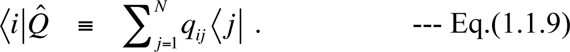

Then, we get the following equation:

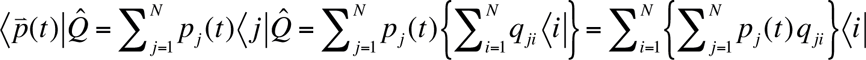

Then, substituting Eq.(1.1.2) for the expression in braces on the leftmost hand side, we have:

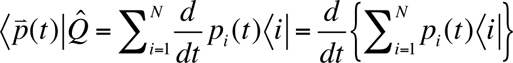

This means that we can recast the defining equation, Eq.(1.1.2), of the Markov model into the equation satisfied by the probability bra-vector 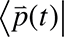:

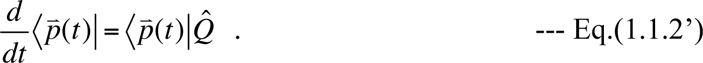

This equation can be integrated as:

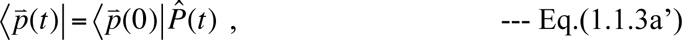

with the finite-time stochastic evolution operator, 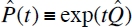. And the counterpart of Eq.(1.1.3b) is:

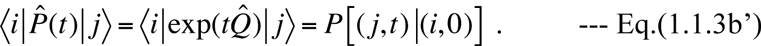

Solving Eq.(1.1.2’) for every possible initial probability bra-vector, 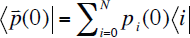, is equivalent to solving the following equation for the operator 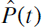

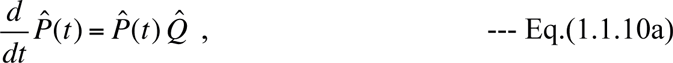

with the initial condition,

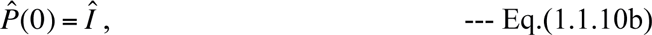

where *Î* is the identify operator: 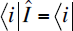 for every state *i*. Thus, if desired, Eqs.(1.1.10a,b) could be considered as the defining equation of the Markov model.

Thus far, we tacitly assumed that the Markov model is time-homogeneous, where the rate matrix *Q*, or the rate operator 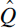, is independent of time *t*. In reality, the transition rate, *q_ij_*, could depend on time due to, *e.g.*, the temporal change of the environment the system is in. Here, we extend the formulation developed above to the system with a *time-dependent* rate matrix, 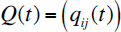, whose operator counterpart is denoted as 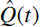. Because the model is no longer homogeneous in time, when we consider a finite-time evolution of probabilities, we need to specify the initial time *t_I_*, in addition to the final time 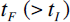. Let 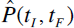 be the operator describing the finite-time stochastic evolution during the closed time interval, [*t_I_, t_F_*], that is: 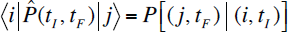 *for* 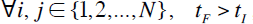, under a continuous-time *time-inhomogeneous* Markov model with the rate operator 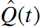. Then, the defining equations, Eqs.(1.1.10a,b), are extended to fit this model as:

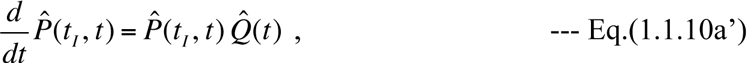

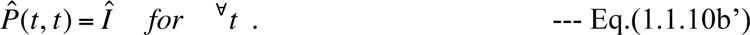

The general solution of the above equations is symbolically given by:

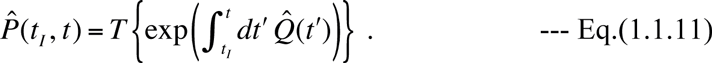

Here *T* {…} denotes (the summation of) the time-ordered product(s), which arrange(s) multiplied operators in the temporal order so that the earliest operator will come leftmost. For example,

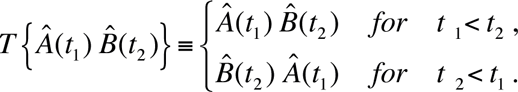

We could regard the time-ordered exponential in Eq.(1.1.11) as defined by a limit:

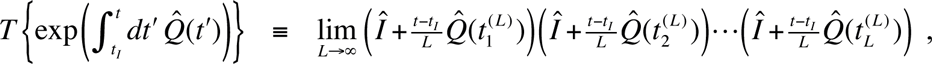

where 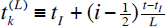, or as defined by a series:

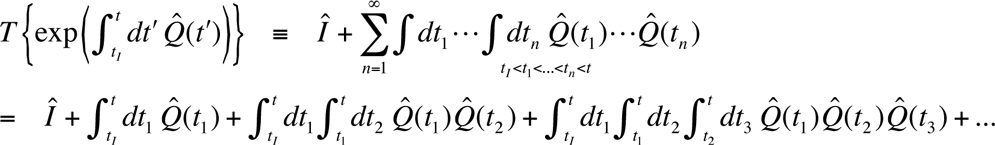

Moreover, the stochastic evolution operator given by Eq.(1.1.11) also satisfies the “backward equation”:

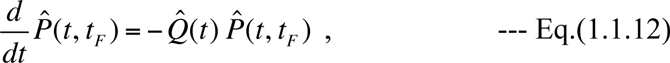

as well as the Chapman-Kolmogorov equation (aka the multiplicativity condition):

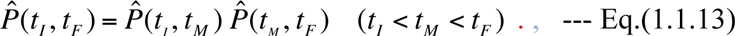

The latter could be rewritten in terms of conditional probabilities:

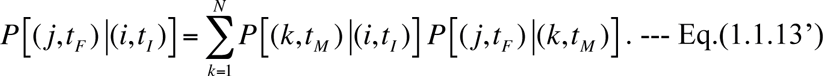

The last equation can be obtained by sandwiching the both sides of Eq.(1.1.13) with 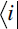 and 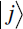, and by inserting the decomposition of the identity operator, 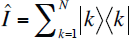, between the two stochastic evolution operators on its right-hand side.

As described above, we have reformulated a continuous-time Markov model on a finite set of states in terms of bra-vectors, ket-vectors and operators. Once we formulated it this way, we could extend the formulation to continuous-time Markov models on any discrete set of states, irrespective of whether it is finite, countably infinite, or uncountable, as long as the state space and the elementary transitions within it are well-defined. In the following sections, we will apply this formulation to describe the evolution of an entire sequence via insertions/deletions.

#### 1.2. Example: application to a model of base substitutions

Traditionally, the studies of molecular evolution via base substitutions have unfolded by using the continuous-time Markov models on the state space consisting of the four bases, *S* = {*T*, *C*, *A*, *G*}, regarding the substitutions at each site as independent of sites (*e.g.*, Yang 2006). The model could be constructed by following the footsteps described in the subsection 1.1, and other by letting the state index *i* take the values *T*, *C*, *A*, *G*. As an illustration, we here consider a simple but nontrivial example, *i.e.*, the model proposed by Felsenstein in 1981. This model is defined with the rate matrix, 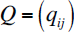, with the elements:

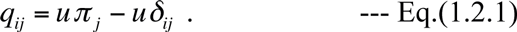

Here *u* gives the scale of the substitution rate, and 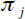 is the equilibrium frequency of state *j*, satisfying 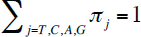. The symbol 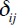 denotes Kronecker’s delta, which equals 1 for *i* = *j*, and 0 for *i* ≠ *j*. In this model, the rate operator 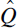 is defined with the equations:

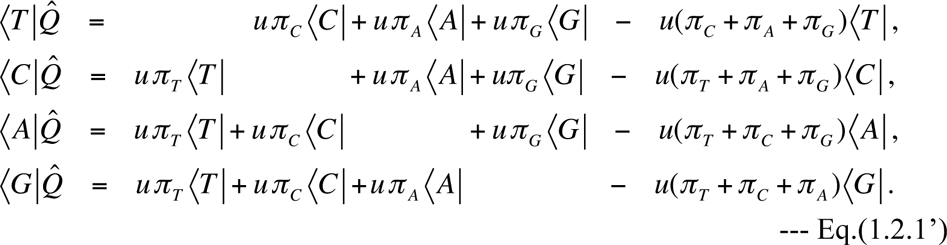

On the right-hand side of each equation, the first three terms represents the substitutions into different bases, and the last term gives the probability decrement resulting from the substitutions of the base on the left-hand side. Substituting Eq.(1.2.1’) into the identity, 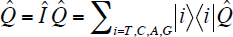, we find that the rate operator can be re-expressed as: 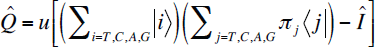. Using this, the stochastic evolution operator, 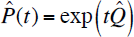, can be calculated as:

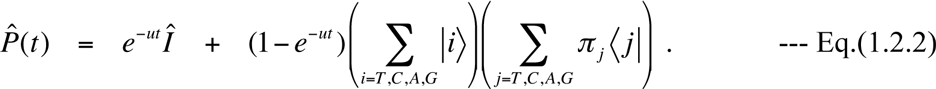

In terms of conditional probabilities, this is rewritten as:

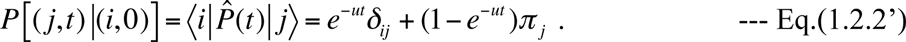

It would be worth a mention that, although we only considered the simplest non-symmetric model here, the bra-ket notation is also applicable to more general substitution models, as we will see in part IV (Ezawa, Graur and Landan 2015c).

#### 1.3. Differences from the quantum mechanics

Although we borrowed the bra-ket notation and the concept of operators from the quantum mechanics (*e.g.*, Dirac 1958; Messiah 1961a), there are some differences between quantum mechanics and the Markov model. For example, in the Markov model, we made the bra-probability vector 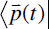 evolve, as in Eq.(1.1.2’), in order to clarify its correspondence with the traditional matrix equation for the conditional probabilities, Eq.(1.1.2). In contrast, in quantum mechanics, it is the ket-vector, 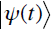, that is usually made evolve. This is simply by convention and, if desired, we could reformulate the quantum mechanics so that the bra-vector will evolve. Another difference, which is conceptually more important, is that, in quantum mechanics, it is the *squared absolute values* of the scalar products, 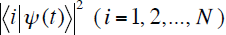, that are interpreted as the probabilities (and thus satisfy 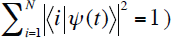). In the Markov model, in contrast, it is the scalar products themselves, 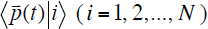, that give the probabilities (and thus satisfy 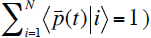. This should be related to another big difference that the time evolution in the quantum mechanics is in the pure-imaginary direction 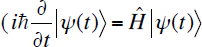, where 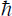 is the Planck constant and 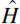 is the instantaneous time-evolution operator called the Hamiltonian), whereas the time evolution in the Markov model is in the real direction (see Eq.(1.1.2’)).

### 2. Definition and formulation of the model of insertions/deletions

As briefly mentioned in Introduction, this study uses a continuous-time Markov model defined on a discrete, infinite state space, in order to describe the stochastic evolution of an *entire* sequence along the time axis via insertions and delentions (indels), without any unnatural restrictions on the possible indel events, and allowing for general indel rate parameters. In this section, we will concretely define and formulate our model step by step.

#### 2.1. State space

Because a Markov process is a timed trajectory in a state space, we first need to set the state space *S* on which our Markov model is defined. We want to describe the indel events on a sequence, thus each element in *S* should represent some state of the sequence. In the bulk of this study, we forget about substitutions in order to focus on indels. Thus we will *not* consider a state as a string of residues that belongs to the space, 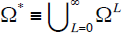, where Ω is the set of residues, that is, the set of four bases (for DNA sequences) or 20 amino-acids (for proteins), as usually done in the past (*e.g.*, Miklós et al. 2004). Instead, we will consider a state as an *array* of a number of *sites*, each of which always contains a residue, and we will represent an insertion/a deletion as an addition/a removal of contiguous sites into/from a position of the array (Figure 1). Depending on how detailed the states have to be represented, different state spaces may be used. We will propose three candidate spaces, *S^I^*, *S^II^* and *S^III^*, as follows. Whichever of these spaces we choose, we will assign a positive integer, *e.g.*, *x*, to each site, in order to represent its coordinate, *i.e.*, its position along the sequence. The leftmost sequence has *x* = 1, and *x* increases by 1 when moving to the right-adjacent site, and the rightmost site has *x* = *L*(*s*), which is the length of the sequence 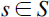.

**Figure 1.**
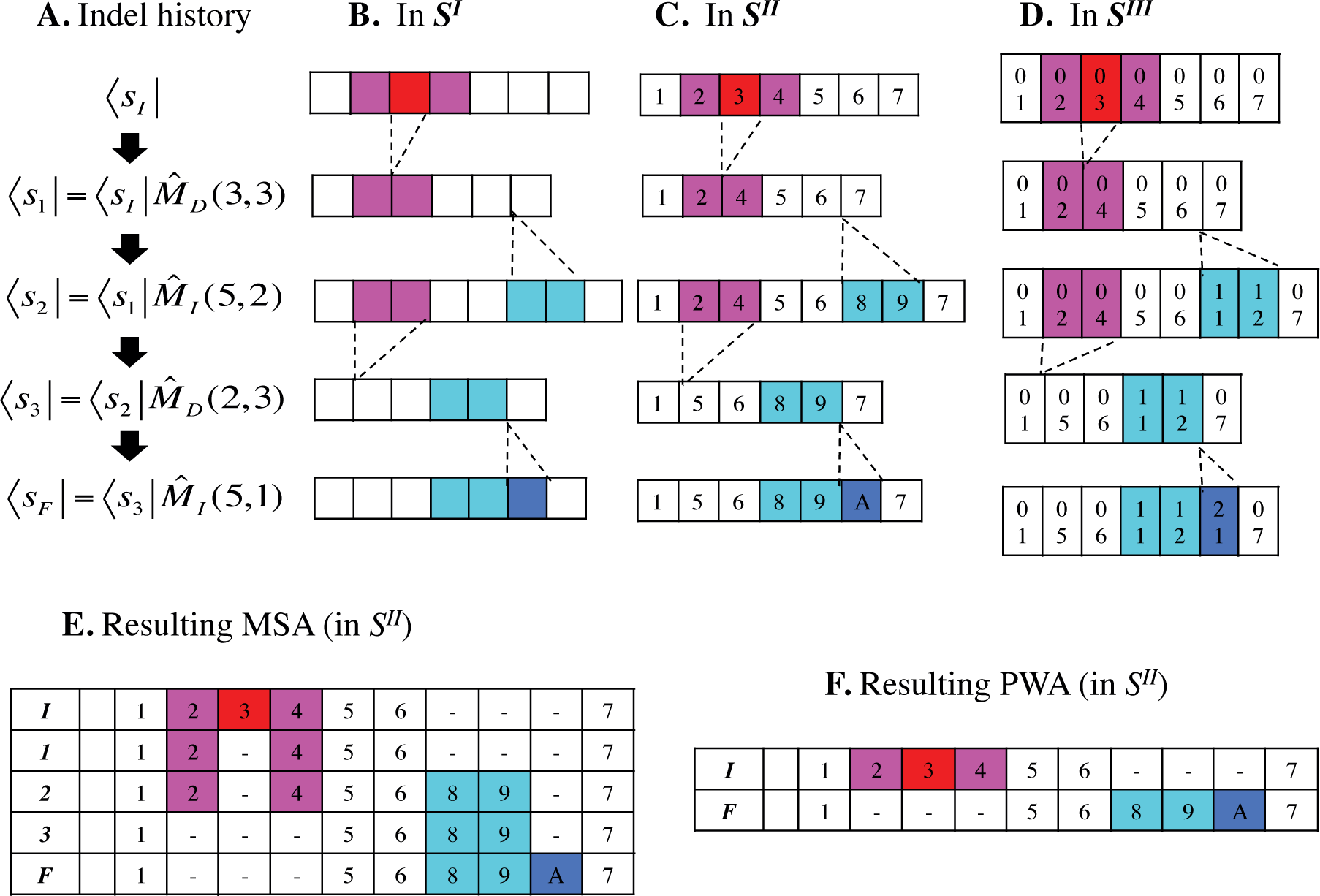
Example indel history and resulting alignments. Panel **A** shows an example indel history. Panels **B, C** and **D** illustrate its representation in the state spaces *S^I^*, *S^II^* and *S^III^*, respectively. Each sequence state in panel A is horizontally aligned with its representation in the three state spaces. **E.** The resulting MSA among the sequence states (in space *S^II^*) that the indel history went through. **F**. The resulting PWA between the initial and final sequences, represented in terms of the states in *S^II^*. In both E and F, the bold italicized characters in the leftmost column are the suffixes indicating the sequence states in panel A. In panels C, E, and F, the number in each site represents its ancestry, but not necessarily its site number (*i.e.*, its spatial coordinate, or order in the sequence). The ‘A’ in the final sequence represents ten (as in the hexadecimal numbering system), to overcome the space shortage. In panel D, each site has two numbers. The upper number is the sequence source identifier, and the lower number represents the relative position of the site in the original source sequence. For clarity, the deleted sites are colored magenta or red, and the inserted sites are colored cyan or blue. It should be noted, however, that these colorings (especially of the deleted ones) are not directly included in the sequence state representations. In this example, the initial state, *S_I_*, is of length 7 (*i.e.*, *L*(*S_I_*) = 7).

(i) *S^I^* (the state space of level 1) is the simplest conceivable space that satisfies the above requirement. In this space, a sequence of length *L* is represented by an array of *L blank* sites (Figure 1B). Because there is no way of distinguishing two sequences of the same length in this space, *S^I^* has a one-to-one correspondence with the set of non-negative integers, 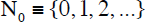, where 0 represents an empty sequence. This space is also equivalent to the aforementioned 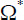 with Ω collapsed into a single-element set. Thus, a state 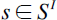 can be uniquely specified with its length, *L*(*s*). A merit of *S^I^* is its simplicity. A drawback is that the record of a trajectory in this space alone cannot completely reproduce an indel process or the alignment of the initial state and the final state. This is because an insertion of the same size changes a state in the same way no matter where in the sequence it occurs, and the same applies to a deletion. (This should be mostly solved if the residue identities are taken into account and if the trajectory of such sequence states is recorded in detail, except in the cases where indels involved repeated subsequences.) We will cover this drawback by keeping the insertion/deletion operators accumulated on the initial state, as a kind of memento. Another drawback of *S^I^* is that it is difficult to introduce positional variations (*e.g.*, in indel rates) aside from the dependence on the (implicit) residue identities of the relevant and neighboring sites, because this space treats all sites equally.

(ii) *S^II^* (the state space of level 2) equips each site of the array with an *ancestry*, which distinguishes the site with those with different ancestries (Figure 1C). The sites with the same ancestry are considered homologous, *i.e.*, descended from the same ancestral site. The set of ancestries, Ƴ, could be anything, as long as it is rich enough to distinguish all possible sets of homologous residues from each other. Although we tentatively let an integer denote an ancestry, what matters is whether the integers are the same or different, but not the relative order among them or their magnitudes. A state 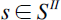 of length *L* can be specified with an *L*–tuple, 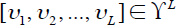. Thus, conceptually, 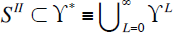 holds, where the first relation is an inclusion and not an equation, because we here consider that different sites in the ancestor, as well as newly inserted sites, have distinct ancestries. (However, an equation could hold if we also take account of duplications and consider that duplicated sites have an identical ancestry. See also part IV (Ezawa, Graur and Landan 2015c) for a related topic.) In the space *S^II^*, we can correctly align two or more sequences by comparing the ancestries of their sites (Figure 1E). Moreover, a trajectory in *S^II^* can uniquely reproduce the history of indels, aside from some ambiguities on deletions involving either end of the sequence (explained in the next subsection). Another merit of this space is that we could introduce positional variations due to factors different from the residue identities of the relevant and neighboring sites. For example, the factors could be the relative positions of the sites in the context of the 3D structure of the protein or RNA products of the gene, or they could be epigenetic contexts, such as predispositions to methylation, chromatin structures, etc. (*e.g.*, Chen et al. 2010; Pink and Hurst 2010). These factors could influence the mutation rate itself and/or the selection pressure on the mutations. In addition, even the same sequence motif could undergo different selection pressures depending on the gene it belongs to or its relative position within the gene. The ancestries, or the ancestral positions, of the sites may model these contextual factors much better than their spatial coordinates along the extant sequences, because the latter could be confounded by indels that hit the sequences during their evolution. This reasoning for the assignment of ancestries to sites seems somewhat similar to the philosophy behind profile HMMs, which are designed to model functional domains or motifs from the MSAs of sequence families (*e.g.*, Durbin et al. 1998; Rivas and Eddy 2013). Indeed, our idea of the “ancestries” of sites was partially inspired by the idea of a “position-specific evolutionary model” (Rivas and Eddy 2013).

(iii) *S^III^* (the state space of level 3) gives richer information than *S^II^*, by elaborating on the ancestry of each site using two attributes, (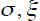), namely, the source of the site (σ) and its relative position within the source (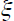) (Figure 1D). The “source” of the site means firstly that whether the site already existed in the initial sequence or not. If so, we assign *σ* = 0. If not, the “source” further means which of the inserted sequences the site belongs to; for example, we could assign *σ* = *k* to the sites inserted by the *k* th insertion in the time order. (However, as the ancestries in the space *S^II^*, we could also consider that the magnitudes of the source identifiers or their relative orders don’t matter.) The “relative positions” of the sites (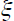’s) are integers representing how far two sites in the same source were from each other, either in the initial sequence state or immediately before they were inserted; the numbers must be consecutive if the sites were adjacent to each other. The “relative position” usually begins with 1, which represents the leftmost site among those inserted, but it could begin at a larger integer if the inserted sequence was a subsequence of a known sequence. Hence, a state 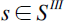 of length *L* is uniquely specified by an *L*-tuple of integer pairs, 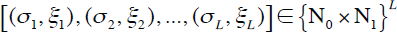, where 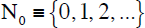 is the set of non-negative integers and 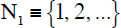 is the set of positive integers. Thus, conceptually, 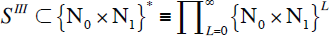 holds. In *S^III^*, the final state gives more than necessary for its alignment with the initial state. The state also gives more detailed (but still possibly incomplete) information on the indels that gave rise to this final state. It may also help annotate the final sequence in more details. And, with some modifications, this state space facilitates the incorporation of other rearrangements, such as duplications and inversions, into our model (see Ezawa, Graur and Landan 2015c).

As we have seen, a higher-level state space contains more information than a lower-level space. Thus, by suppressing some information, a higher-level space can be reduced to a lower-level space, but the former can never be recovered from the latter. For example, although a timed trajectory in the state space of either level 2 or 3 can fully recover the indel process, a timed trajectory in the level 1 state space cannot. Another important note is that, even in the state space of level 3, the alignment of the initial state with the final state cannot fully recover the indel history in general. To recover the full indel history, it is necessary to record the full trajectory of the sequence evolution in either *S^II^* or *S^III^*. We will do this concisely and in a focused manner by bookkeeping the successive actions of insertion and deletion operators, which will be introduced in the next subsection, on the sequence. Once we introduce this bookkeeping method, we could actually recover the full indel history even if we work with *S^I^*. Hereafter, the symbol *S* denotes the state space when we do not need to specify its level of details.

#### 2.2. Insertion and deletion operators

Here we will introduce the key components of our model formulation, namely, insertion operators and deletion operators. As in the long indel model of Miklós et al. (2004) and the indel model of Dawg (Cartwright 2005), we consider that the sequence in question, 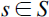, whose length will be denoted as *L*(*s*), is embedded in a sequence of a practically infinite length.

Let 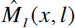 be the “insertion operator” that inserts a contiguous array of *l* sites between the *x* th and the (*x* + 1) th sites of the sequence *s*, when 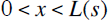. For example, the action of 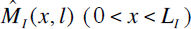 on an initial sequence, 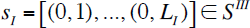, could be expressed as:

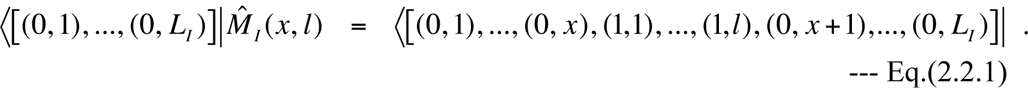

We also allow the 1st argument *x* to be 0 or *L*(*s*); we define 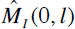 as an operator to prepend an array of *l* sites to the left-end of *s*, and define 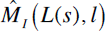 as an operator to append the array to the right-end of *s*. However, we will not consider the action of 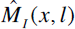 with *x* < 0 or *x* > *L*(*s*) on *s*.

Let 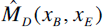 (with 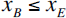 be the “deletion operator” that removes the sub-array between (and including) the *x_B_* th site and the *x_E_* th site from the sequence *s*, if 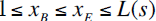. For example, the action of 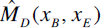 (with 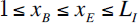) on *S_I_* as defined above could be expressed as:

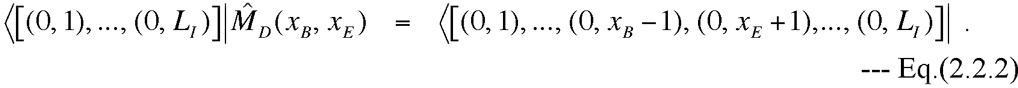

Because we consider the sequence *s* as embedded in a practically infinitely long sequence, we also allow deletions to stick out of an end or both ends of the sequence. We define the action of 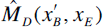 with 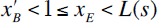 to be identical to that of 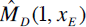, *i.e.*, the removal of the sub-array of *s* between (and including) the left-end and the *x_E_* th site. Likewise, we define the action of 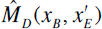 with 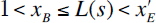 to be identical to that of 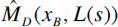, *i.e.*, the removal of the action of 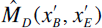 with 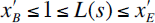 is defined as identical to that of 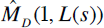, *i.e.*, the deletion of the whole sequence *s*, which results in an empty array, []. These identifications of the end-involving deletions were already known (Miklós et al. 2004; Cartwright 2005), but we are the first to formulate the identifications in terms of the equivalence relations between operators (see Eqs.(2.3.1a,b,c) in the next subsection).

With these definitions, a particular insertion or deletion operator acting on a particular state in the space *S* unambiguously results in another particular state in *S*. Thus, successive actions of some insertion and deletion operators on an initial state uniquely determine an indel history, or an *untimed* trajectory of the states in *S*. Figure 1A shows an example of such successive actions of operators, and panels B, C, and D in Figure 1 are its representations in the state spaces *S^I^*, *S^II^*, and *S^III^*, respectively. The indel history shown in Figure 1 can be recapitulated as the following “bookkeeping” representation of the accumulated actions of the indel operators:

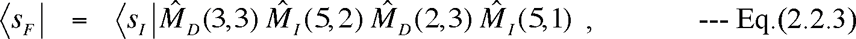

from which the (untimed) trajectory in the state space and the MSA are also recoverable. Alternatively, we could also represent the indel history as the initial state (*s_I_*) and an *ordered* set of the indel operators, 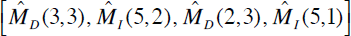 As shown in Figure 1E, the MSA of the initial, intermediate, and final sequences can be easily constructed by unfolding the bookkept actions of the indel operators, that is, by inserting gaps aligned with inserted sites into sequence states before the insertion, and by inserting gaps aligned with deleted sites into sequence states after the deletion. Then, by removing the intermediate sequences from the MSA, and possibly by removing the resulting “null” columns that contain only gaps, the PWA between the initial and final sequences can also be obtained (Figure 1F).

#### 2.3. Equivalence classes of indel histories during time interval (I)

In many applications, we are mainly interested in the pairwise alignment (PWA) between the initial and final sequences in the evolution during a time interval, [*t_I_*,*t_f_* which often corresponds to a branch in a phylogenetic tree. In general, a PWA could result from many different indel histories. Therefore, it is useful to identify some typical groups of indel histories that yield the same PWA.

First of all, using the definitions of the sticking-out deletion operators given in Subsection 2.2, we can set the following unary equivalence relations:

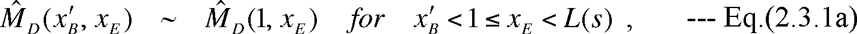

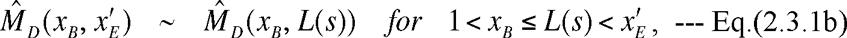

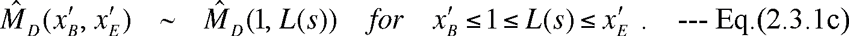

Here *L*(*s*) is the length of the sequence *s* that the operators act on. Using these unary equivalence relations, we first rewrite the sticking-out deletion operators with the equivalent operators that do not stick out of the sequence ends. Then, we consider more complex equivalence relations below.

Let us second consider the simplest “complex” histories, each of which consists of two indel events separated by at least a site that was preserved throughout the time interval (called a “preserved ancestral site” (PAS) hereafter). Figure 2, panels A-C gives an example, where two indel histories (panels A and B) result in the same PWA (panel C). The indel history in Figure 2A can be recapitulated as 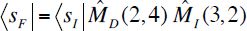, whereas the indel history in Figure 2B can be recapitulated as 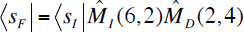. Even in the state space *S^III^*, however, both result in the identical state (bottom of panels A and B of Figure 2): 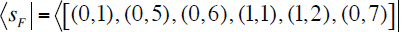. Here we assumed that 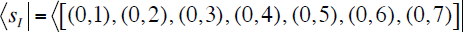. Thus, as far as states in *S^III^* are concerned, we get the following binary equivalence relation:

**Figure 2.**
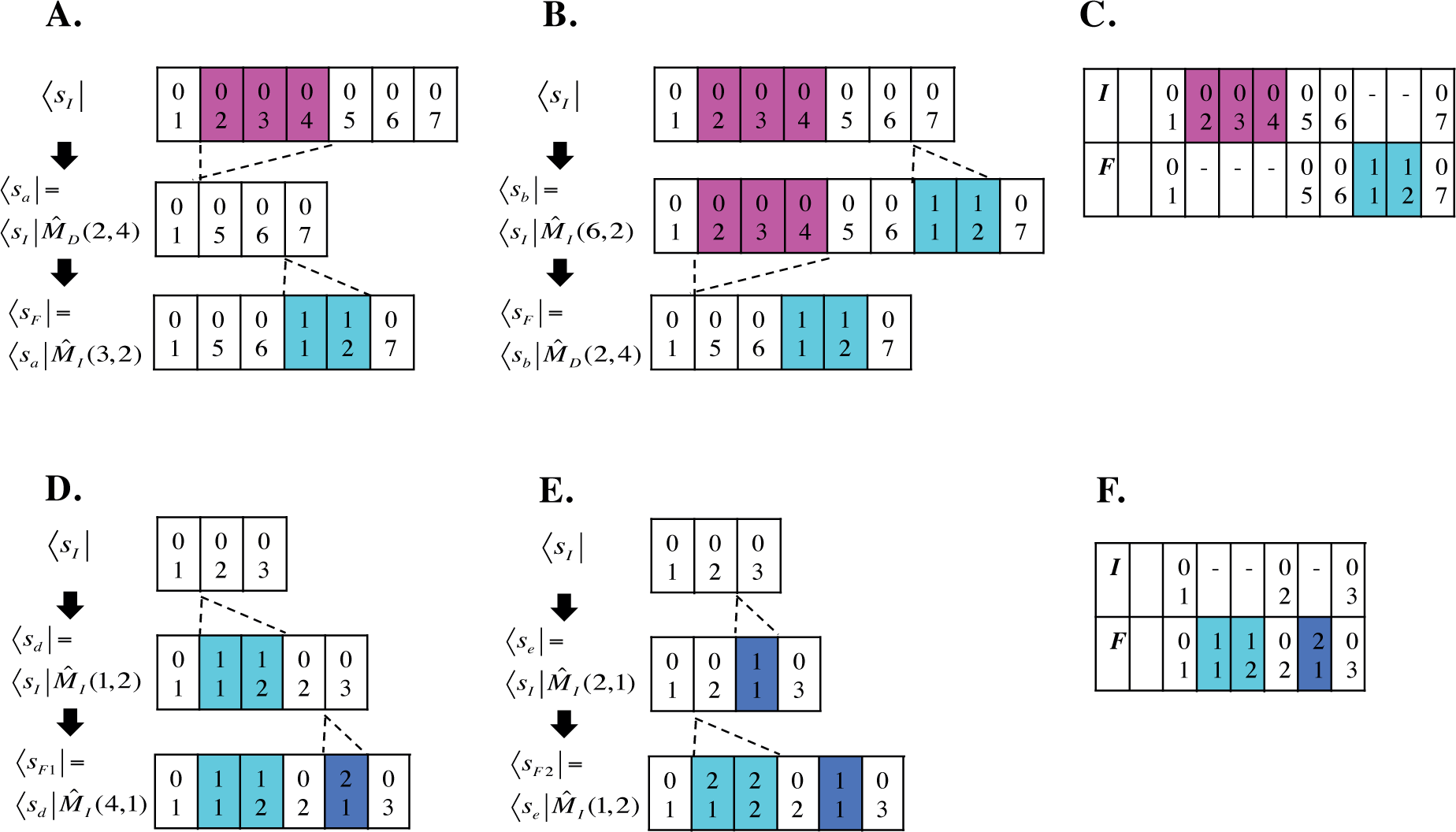
Equivalent indel histories involving two non-overlapping indel events. Panels **A** and **B** show equivalent histories, each involving a deletion and an insertion, which result in the same alignment(panel **C**). Panels **D** and **E** show equivalent histories, each involving two insetions. Both of them giv rise to the alignment in panel **F**. All indel histories are represented in the state space *S^III^*. As in Figure 1 deleted sites are colored magenta, and inserted sites are colored cyan or blue.

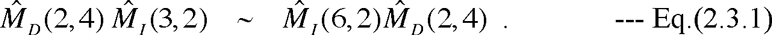

Of course, the two histories give the same PWA (Figure 2C). Another example is given in Figure 2, panels D-F, where two insertions are involved. The history in Figure 2D can be recapitulated as 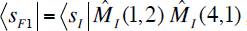, whereas the history in Figure 2E can be recapitulated as 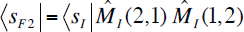. In this case, the resulting states in *S^III^* are slightly different, as indicated by the different state names: 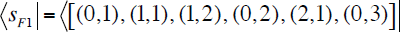, and 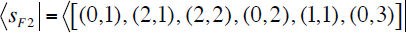.

Here we assumed that 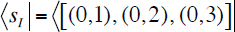. Therefore, the two operator products do not appear equivalent in *S^III^* in its strict sense. However, if we remember that what matters regarding the origin identifier (the first number in each pair of parentheses) is only whether the identifiers are the same or different, 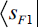 and 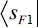 are indistinguishable. Moreover, the two histories give the same PWA (Figure 2F). Thus, in *S^III^* in this broad sense (and of course in *S^II^*), we get the following binary equivalence relation:

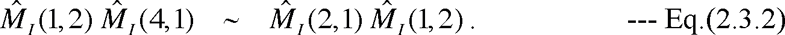

These equivalence relations, Eq.(2.3.1) and Eq.(2.3.2), can be generalized to provide the following four sets of binary equivalence relations in terms of PWA:

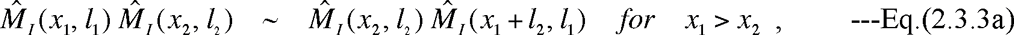

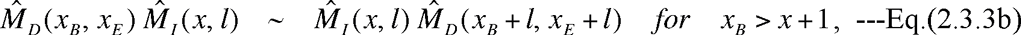

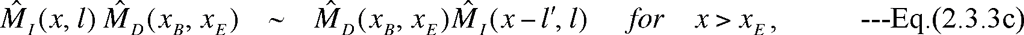

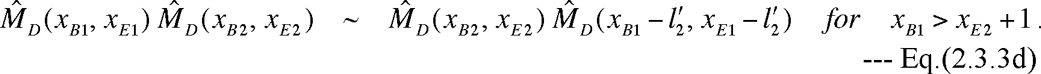

Here, 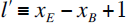 in Eq.(2.3.3c), and 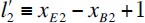 in Eq.(2.3.3d). These equivalence relations could be re-expressed in the following words. “The operator representing the event on the left along the sequence will not change whether it comes first or second. The operator representing the event on the right will shift its operational position to the left/right by the number of sites deleted/inserted before its operation, when it comes second.”

Now, we can extend the binary equivalence relations, Eqs.(2.3.3a-d), to the equivalence relations among more general complex indel histories, each consisting of more than two indel events. Let us consider a history of *N* indel events, which begins with an initial state 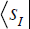 and is recapitulated as:

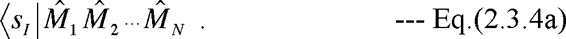

Here 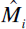 is the operator representing the *i* th event 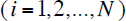 in the temporal order, which is 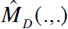 (for a deletion) or 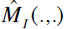 (for an insertion) with appropriate arguments. This indel history is also represented as 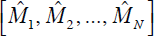 on *s_I_* Given an indel history, we can identify ancestral sites that have been kept undeleted during the history. Suppose that such preserved ancestral sites (PASs) separate the indel events 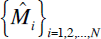 in the *global* history 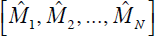 into *K local* subsets of indels, each of which is confined either between a pair of PASs or between a PAS and an end of the resulting PWA. Number the *K* local subsets as *k* = 1, 2, …, *K* from left to right, and let *N_k_* be the number of indel events in the *k* th local subset. Here the numbers satisfy 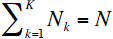. And let 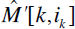 be the element of 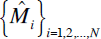 representing the *i_k_* th event (in the temporal order) in the *k* th local subset (*i_k_* = 1, 2,…, *N_k_*; *k* = 1, 2, …, *K*). Then, repeatedly applying the binary equivalence relations, Eqs.(2.3.3a-d), between the operators representing events belonging to *different* local subsets, we can move the operators around in the product in Eq.(2.3.4a) and find the following expression that is equivalent to Eq.(2.3.4a):

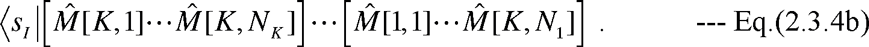

Here 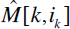is an operator that was obtained from 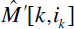 through the series of equivalence relations Eqs.(2.3.3a-d) that brought Eq.(2.3.4a) into Eq.(2.3.4b). As the operators in Eq.(2.3.4a), the operators in each pair of large square parentheses are arranged in temporal order, so that the earliest event in each local subset will come leftmost. But it should be noted that the order among the pairs of large square parentheses is the opposite of the actual spatial order among the local subsets, so that the rightmost local subset along the sequence (the *K* th one) will come leftmost in the product of operators. In this way, the operators in each local subset, *e.g.*, 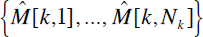, are exactly the same as those when the events in the subset alone struck the initial state 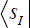. Thus the series of operators, 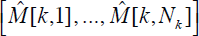, for the *k* th local subset defines the *k* th *local indel history* that was isolated from the *global* indel history 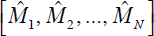 on 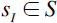.

For example, the history of *N* = 4 indels in Figure 1, recapitulated as Eq.(2.2.3), is equivalent to the following product of local indel histories: 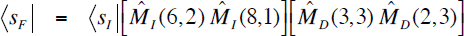

In this case, *K* = 2 and *N*_1_ = *N*_2_ = 2. The operators representing local indel histories are: 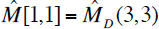, 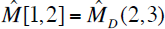, 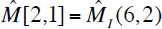, and 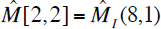.

Now, let us consider a history of *N* indel events other than that represented as Eq.(2.3.4a). If the history is shown to be equivalent to Eq.(2.3.4b) through a series of equivalence relations, Eqs.(2.3.3a-d), then it should also be connected to Eq.(2.3.4a) though another series of Eqs.(2.3.3a-d). Therefore, it should be equivalent to Eq.(2.3.4a) in this sense. Hence, we can define a particular equivalence class to be the set of all global indel histories that can be “decomposed” into the identical set of local indel histories, such as Eq.(2.3.4b), only through a series of equivalence relations, Eqs.(2.3.3a-d), between operators representing indel events separated by at least one PAS. This equivalence class will become essential to the proof of the factorability of a PWA probability. Thus, we will call it the **“local-history-set (LHS) equivalence class.”** In the equivalence class defined by a local history set (LHS), 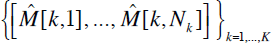 (with 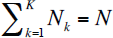, on an initial sequence state 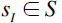, there are 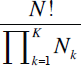 LHS-equivalent global indel histories beginning with *s_I_*. Each of the global histories corresponds to a way of reordering *N* indel events while retaining the relative temporal order among *N_k_* events within the *k* th local indel history (for every *k* = 1, …, *K*).

We can also identify equivalence relations involving the product of two operators representing overlapping indels or indels *not* separated by a PAS. Some of such relations are given in Appendix A1 (and illustrated in Figure 3). They are useful when discussing further equivalence relations between *local* indel histories giving rise to the identical local PWA. Most, if not all, of the equivalence relations between indel histories should be identified by the repeated applications of these relations, in addition to Eqs.(2.3.3a-d), and possibly Eqs.(2.3.1a-c).

**Figure 3.**
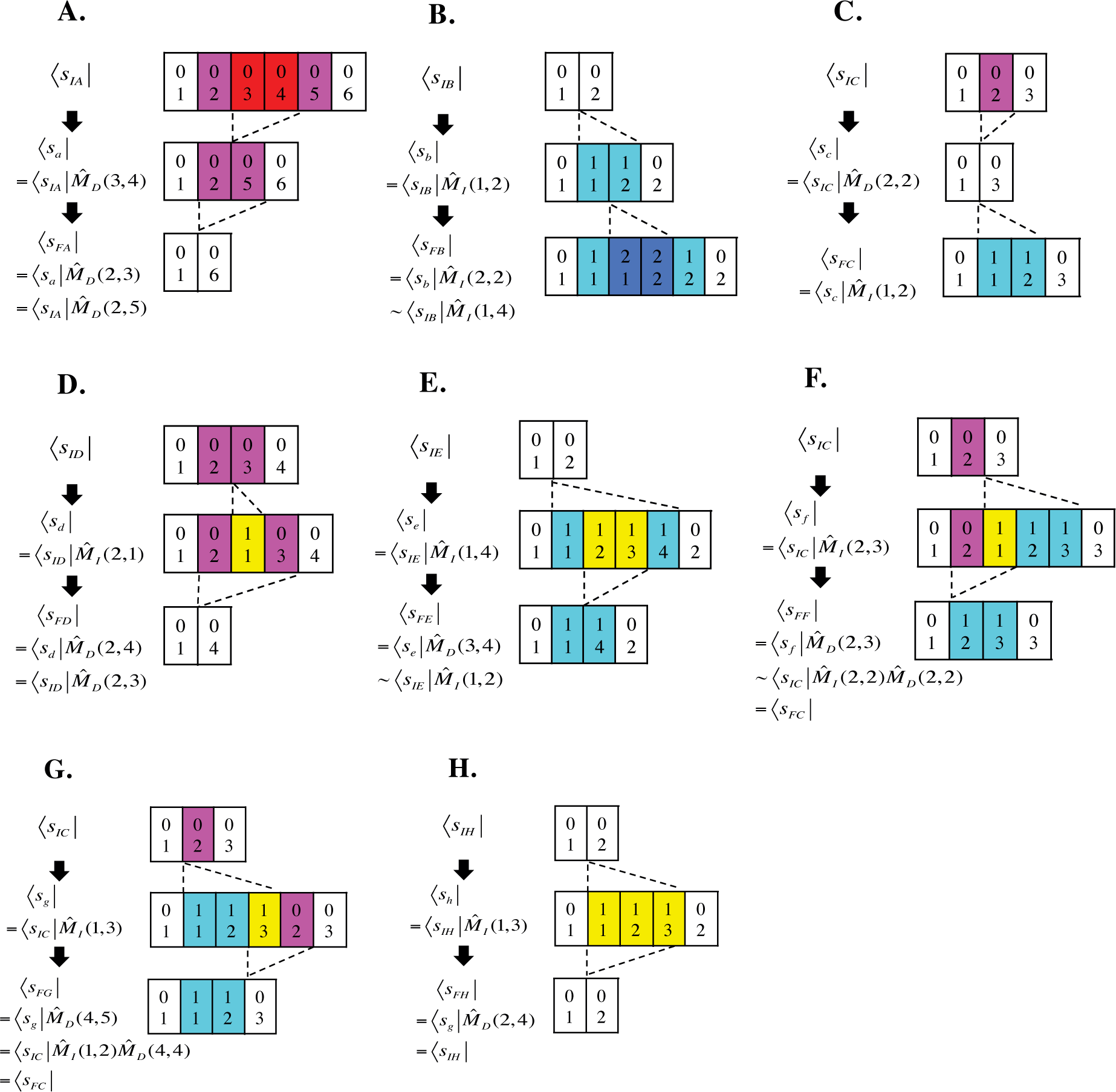
Equivalence relationships involving indels that overlap or touch each other. **A.** The successive action of two nested (or mutually touching) deletions 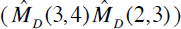 is equivalent to a single deletion (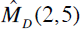) **B.** The successive action of two nested (or mutually touching) insertions 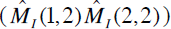 is equivalent to a single insertion (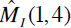) **C.** A deletion (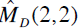) followed by an insertion between the deletion-flanking sites (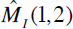) **D.** An insertion (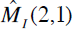) followed by the deletion of a region encompassing the inserted subsequence (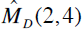) is equivalent to a single deletion (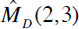) **E.** An insertion (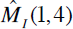) followed by the deletion of a region nested within the inserted sequence (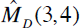) is equivalent to a single insertion (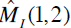). **F.** If the state space is *S^I^* or *S^II^*, an insertion (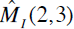) followed by a left-overlapping deletion (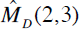) is equivalent to a non-overlapping but mutually touching pair of an insertion and a subsequent deletion 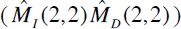 which is also equivalent to the result of panel C. **G**. If the state space is *S^I^* or *S^II^*, an insertion (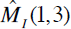) followed by a right-overlapping deletion (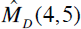) is equivalent to a non-overlapping but mutually touching pair of an insertion and a subsequent deletion 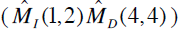, which is also equivalent to the result of panel C. **H.** An insertion (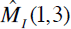) and a subsequent exact deletion of the inserted subsequence (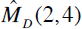) result in a sequence state identical to the initial one. The sequence states are represented in the space *S^III^*. The magenta and red boxes represent sites to be deleted. The cyan and blue boxes represent inserted sites. The yellow boxes represent inserted sites that are to be deleted. In each panel, the “=” represents the exact equality of the sequence states in the state space *S^III^*. The states connected by “∼” are not exactly equal in *S^III^*, but they give the same pairwise alignment with the initial state. (In other words, they become equal in *S^II^*).

#### 2.4. Evolutionary rate operator

Here we finalize the definition of our continuous-time Markov model by giving the evolutionary rate operator in terms of the insertion and deletion operators. First consider its action on the bra-vector, 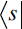, of a sequence state 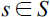 of length *L*(*s*) = *L*.

In this case, the insertion operators that can act on 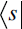 are 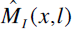 with *x* = 0, 1,…, *L* and *l* ≥ 1, and the deletion operators that can act on 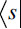 are 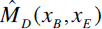 with 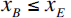, 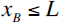, and 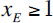. We begin with a very general situation where the rate parameters, 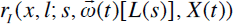 for the insertion 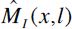 and 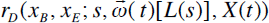 for the deletion 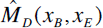, could depend on the sequence indel state (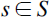) including its genomic and epigenomic contexts (as far as the state space *S* can accommodate), the residue identities filling the sites of the sequence 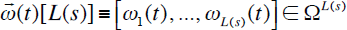, and other external factors (*X*(*t*)) including the cellular and subcellular locations of the gene product, population dynamics, ecological environment, climates, etc. The latter two arguments are considered as time-dependent parameters. It should be noted that, because we do not explicitly consider the sequence evolution via substitutions in the bulk of the paper, we regard 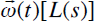 as an average behavior of the sequence residue states. Thus, sharp changes in the residue states, such as the creation or annihilation of the sequence motifs that drastically enhance or suppress the indel rates, will not be considered here. Such cases will be briefly considered in part IV (Ezawa, Graur and Landan 2015c). In the following, to simplify the notation, we will not explicitly express the dependence of the rate parameters on 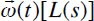, and *X*(*t*). Instead, we will collectively represent it by the dependence on time *t*, like 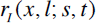 and 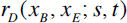. Given a set of indel rate parameters as above, the rate operator restricted to a subspace of states, 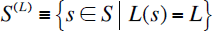, is defined by the following action on the state bra-vector 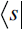 for 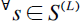:

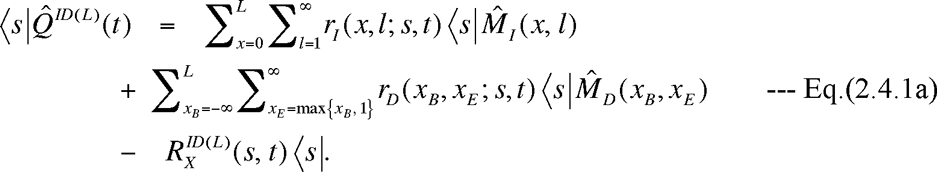

Here the first and the second double-summations give the state changes via an insertion and a deletion, respectively. The third term with the exit rate:

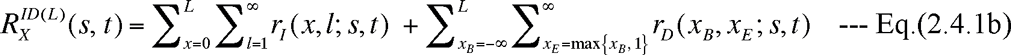

is necessary for keeping the total probability to be 1. From Eqs.(2.4.1a,b), we can define the indel rate operator, 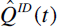, in the whole state space *S*, by using the decomposition of the identity operator, 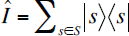, as:

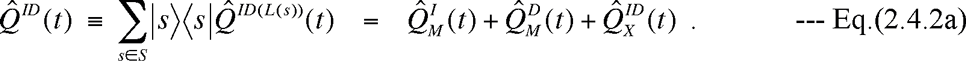

Here

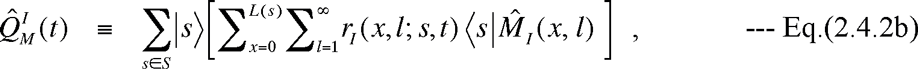

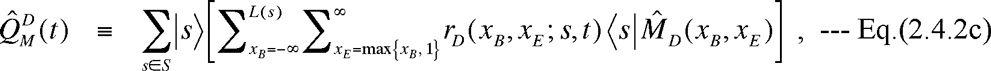

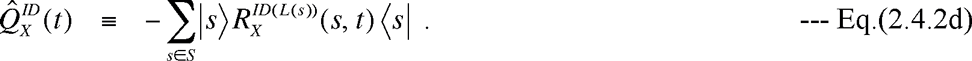

In practice, the probabilities of insertions/deletions of extremely long (sub-)sequences are practically zero, due to physical restrictions (*e.g.*, the chromosome length) or for biological reasons (*e.g.*, purifying selection). Thus, we could safely limit the lengths of insertions and deletions to less than or equal to some “cut-off” values. Let them be denoted here as 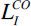 and 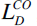, respectively. Then, Eqs.(2.4.2b,c) could be rewritten as:

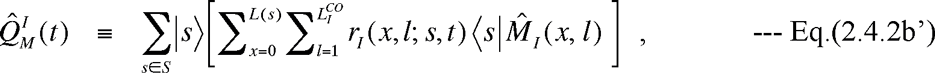

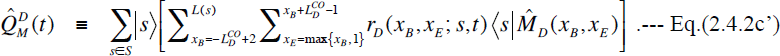

And, Eq.(2.4.1b) could also be rewritten as:

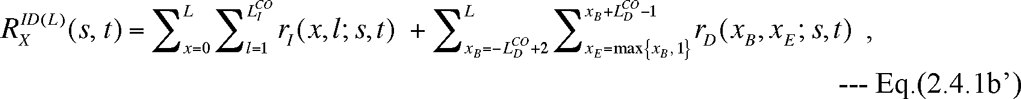

which in turn gets Eq.(2.4.2d) re-expressed as well.

Using the unary equivalence relations, Eqs.(2.3.1a,b,c), we can further decompose 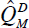, defined in Eq.(2.4.2c’), into contributions from the deletions in the middle of the sequence, on the left-end, on the right-end, and from the whole-sequence deletions, as given in Eqs.(A2.1a-e) in Appendix A2. This re-expression of Eqs.(2.4.2c’) could sometimes simplify theoretical thinking. It could also save computational costs by doing away with deletions that stick out of the boundaries of the sequence under consideration. It will be used only rarely in this paper.

If the state space *S^1^* is used, the continuous time Markov model defined by the Eqs.(2.4.2a-d) or their equivalents, Eqs.(2.4.2b’,c’) or Eqs.(2.4.3a-d), with time-independent parameters, is practically equivalent to the indel component of the substitution/insertion/deletion model equipped with a general rate grammar, as proposed by Miklós et al. (2004). A major difference between their and our formulations is that, whereas state trajectories played a central role in their model, we focused on indel histories, which enabled us to prove the factorability of the alignment probability calculated from the first principle. Although Miklós et al.’s general rate grammar (which they merely proposed) can accommodate the dependence of indel rate parameters on the sequence context through the residue identities of the sites in the sequence, it cannot accommodate their dependence on the ancestries of the sites, which could be a proxy of, *e.g.*, the 3D structural, genomic and epigenomic contexts of the sequence. Our general Markov model, in contrast, could accommodate the site ancestry dependence of indel rates, if we use the state space *S^II^* or *S^III^*.

Here, we give a couple of special cases of our general model, with the state space *S^I^*. First, the indel model for Dawg (Cartwright 2005) is equivalent to the model with the rate operator given by Eqs.(2.4.2a,b’,c’,d) and Eq.(2.4.1’) with the homogeneous, time-independent indel rate parameters:

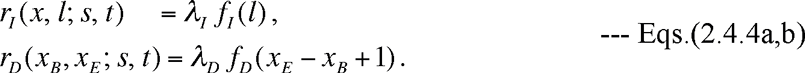

Here *λ_I_* and *λ_D_* are the per-location rates of insertion and deletion, respectively, and *f_I_*(*l*) and *f_D_*(*l*) are the distributions of insertion lengths and deletion lengths, respectively. Because Dawg’s model does not impose the time-reversibility condition, we can take *λ_I_*, *λ_D_*, *f_I_*(*l*), and *f_D_*(*l*) freely, as long as they are all non-negative and satisfy 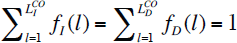for some cut-off values 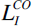 and 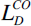. The exit rate, Eq.(II-4.1b’), can be calculated as:

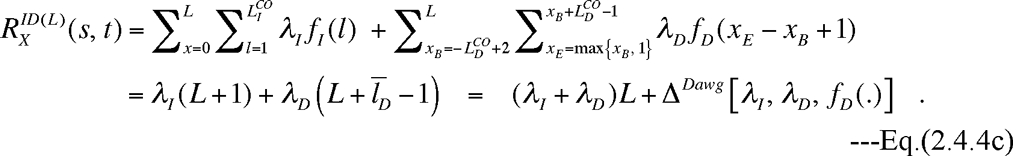

Here 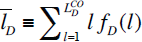 is the average deletion length, and 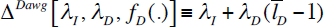 is a constant that depends on the indel rate parameters.

The long-indel model of Miklós et al. (2004) is very similar to the model of Dawg, but it also shows some differences. It could be defined by Eqs.(2.4.2a,b’,c’,d) and Eq.(2.4.1’) with the homogeneous, time-independent indel rate parameters:

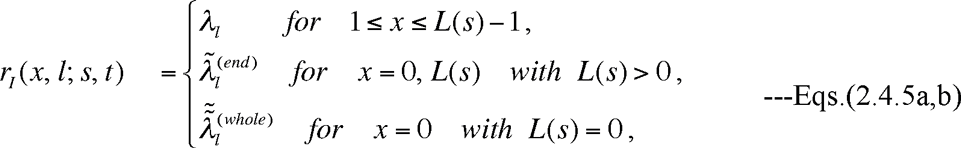

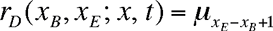

It should be noted that each insertion rate parameter in the original paper of Miklós et al. (2004) includes a multiplication factor, representing the probability of residue states that filled the inserted sites. The factor is omitted in the bulk of this paper, because we consider it to be treated in conjunction with the substitution model (as briefly discussed in Discussion). This long-indel model was required to satisfy the detailed-balance conditions and thus to be time-reversible. The appropriate state space for the indel component of this model is *S^I^*, thus a state 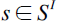 is uniquely determined by specifying its length, 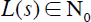. Letting 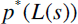 be the equilibrium distribution of the sequence length, the detailed-balance conditions are: 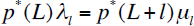 for the bulk, 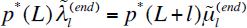 for the sequence ends with *L* > 0, and 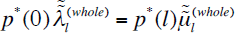 for *L* = 0, all for 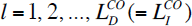. Here, 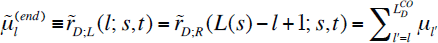 with 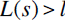 is the “effective rate” of the deletion of length *l* from either end of the sequence. And 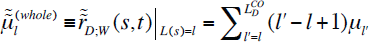is the “effective rate” of the whole-sequence deletion of length *l*. These equations for 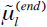 and 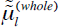 were obtained by substituting Eq.(2.4.5b) into the definitions of 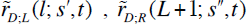, and 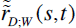 given by Eqs.(A2.1c,d,e), respectively. Solving the detailed balance conditions yields, as described in Miklós et al. (2004):

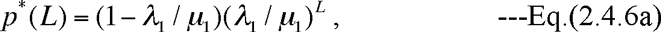

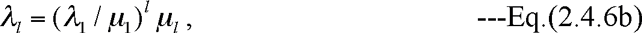

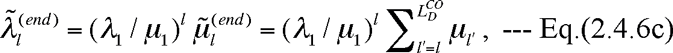

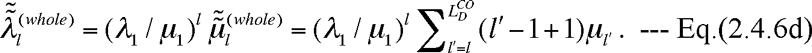

Thus, the sequence length must follow a geometric distribution with a fixed elongation probability 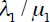, and the insertion length distribution and the deletion length distribution must depend on each other through Eqs.(2.4.6b,c,d). Aside from these differences due to the time-reversibility, the long indel model is very similar to Dawg’s indel model. We can easily see the correspondence by setting:

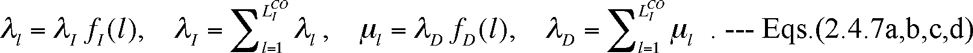

Using this correspondence, the exit rate in the long-indel model is given in a very similar form as in Dawg’s model:

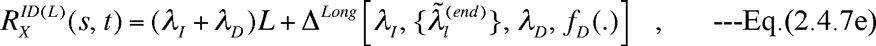

where 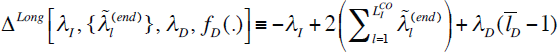 is a constant that depends on the indel rate parameters. One of the major differences between the two models is in the length distribution of insertions on a sequence end. The long indel model forces it to balance the length distribution of deletions on a sequence end, whereas Dawg’s model merely sets it equal to the (homogeneous) insertion length distribution at an inter-site position. We suppose that this particular difference does not matter so much in their applications to practical sequence analyses, where sequence ends are likely to be determined by artificial factors, such as sequence annotation, homology detection, etc.

Back to the general model, once the rate operator 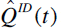 is given by Eqs.(2.4.2a,b’,c’,d) with Eq.(2.4.1’), we could at least formally solve the extension of Eqs.(1.1.10a’,b’) to the space state 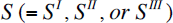:

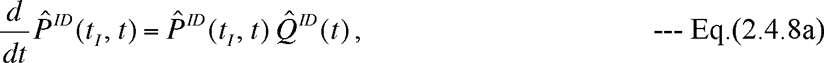

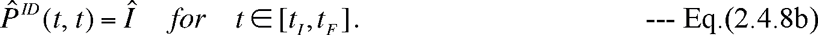

This yields the formal general solution for the stochastic indel evolution operator for the time interval [*t_I_*, *t*]:

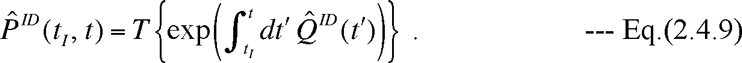

By definition, the evolution operator naturally satisfies the Chapman-Kolmogorov equation:

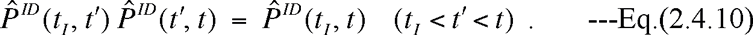

In practice, however, because *S* is an infinite state space, a naïve numerical computation of Eq.(2.4.9) is impossible. Analytic solutions to Eqs.(2.4.8a,b) cannot be obtained, either, except in special simple cases where the indel process of each site and each inter-site position can be handled separately, such as in the TKF91 model (Thorne et al. 1991). Good news is that 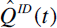 is quite sparse, that is, it connects each state 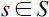 with only a finite number of states. Therefore, if we are only interested in the finite-time evolution of a sequence starting with a given state 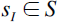, only a small subset of *S* will need to be explored. This is because we are essentially dealing with diffusion processes, like random walks, from a point (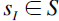). Taking account of this idea, we could approximately perform a numerical computation of Eq.(2.4.9) by, *e.g.*, using the definition of the time-ordered exponential:

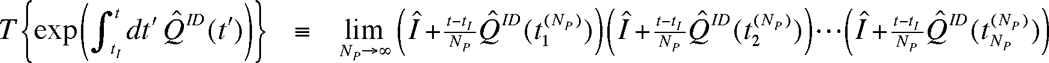

With 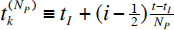. In the next section, however, we will rewrite Eq.(2.4.9) into a more convenient and insightful form, by using techniques of the perturbation theory in physics (*e.g.*, Dirac 1958; Messiah 1961a).

### 3. Perturbation expansion of alignment probability

#### 3.1. Perturbation expansion of probability of PWA between descendant and ancestral sequences

In the perturbation theory of quantum mechanics (*e.g.*, Dirac 1958; Messiah 1961b), the instantaneous time evolution operator 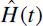 is considered as a sum of two operators, 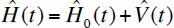, and the time evolution of the system is described as if the system mostly evolves according to the well-solvable time-evolution operator 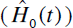 and is occasionally perturbed by the “interaction” operator 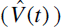.

Here we apply the technique of such perturbation theory to our general continuous-time Markov model. We first re-express our rate operator as:

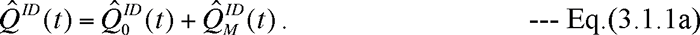

Here 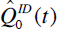 is the operator describing the mutation-free evolution, and 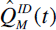 is the operator describing the state transition due to a mutation (indel):

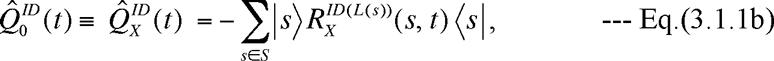

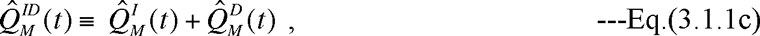

with 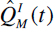 and 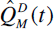 defined in Eqs.(2.4.2b,c). Using Eq.(3.1.1a), the time-differential equation of the stochastic evolution operator, Eq.(2.4.8a), can be rewritten as: 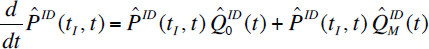, which is further rewritten as:

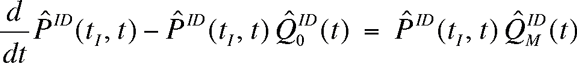

Multiplying each side of the above equation by 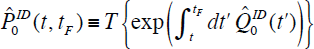 from the right, and using the “backward equation” Eq.(1.1.12) with 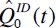 substituted for 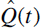, we have:

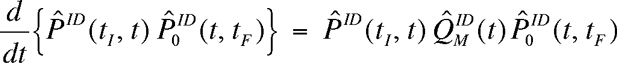

Performing the time integration from *t_I_* to *t_F_* of both sides, and using 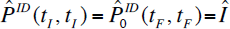, we get an important integral equation:

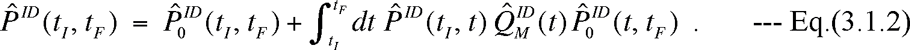

Now, we formally expand 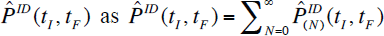, where 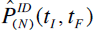 is the collection of terms containing *N* indel operators each. Substituting this expansion into Eq.(3.1.2) and comparing the terms with the same number of indel operators on both sides, we find:

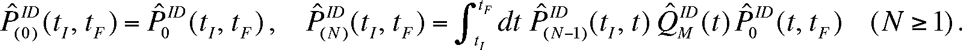

The second equation can be recursively solved to give:

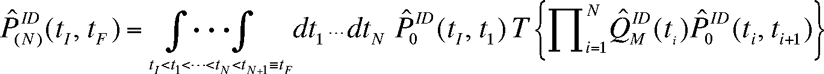

for *N* ≥ 1. Substituting this expression into the above expansion, we finally obtain the formal perturbation expansion of the stochastic evolution operator:

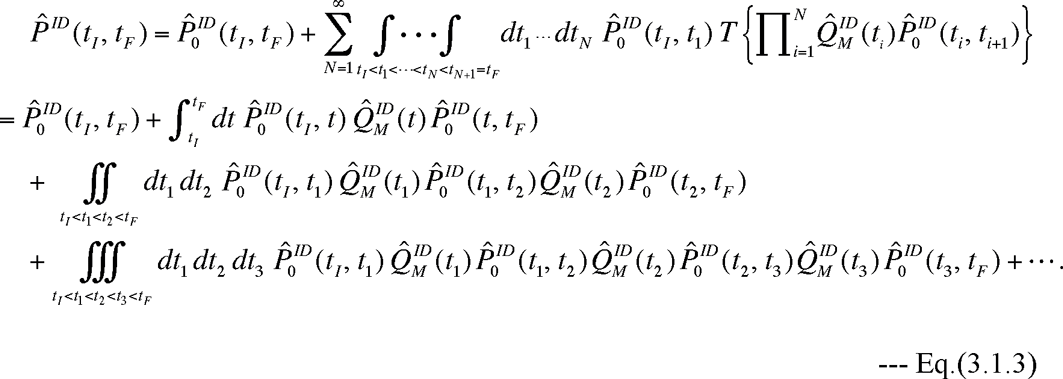

From this expansion, we can see that 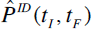 also satisfies another important integral equation:

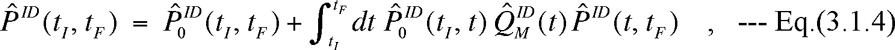

which could also be derived from the backward equation, Eq.(1.1.12), with Eq.(3.1.1a) substituted for 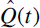. Actually, Eqs.(3.1.2,3,4) hold for general continuous-time Markov models, not limited to the indel evolutionary model, if we replace 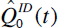 with any “perturbation-free” rate operator and replace 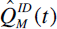 with the remainder, which will be treated as a “perturbation” operator. (See, *e.g.*, part IV (Ezawa, Graur and Landan 2015c).) [NOTE: Historically, it was Feller’s Theorems 1 and 2 (1940) that first proved that Eq.(3.1.3) gives the general solution to Eqs.(2.4.8a,b) in a general context (not limited to indel models). And Gillespie (1977) gave a more intuitive derivation of the solution (actually Eqs.(3.1.8a,b) below). The proof we gave here serves as a bridge between Feller’s mathematically rigorous proof and Gillespie’s intuitive derivation. Our proof here first derived the integral equation Eq.(3.1.2) satisfied by the *entire* stochastic evolution operator (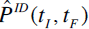) and then derived the recursion relations satisfied by 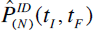’s. Eq.(3.1.2) will play an important role in part II (Ezawa, Graur and Landan 2015a). Moreover, our operator representation provides a more concise and intuitively clearer derivation than Feller’s theorems, which were represented in terms of probability components. Furthermore, as we can see in Part IV mentioned above, our derivation via the perturbation formulation could be more flexible than Feller’s and Gillespie’s, which always separate all exit-rate terms from all state-transition terms.]

In the present case, we could obtain a more concrete expression. First of all, from Eq.(3.1.1b), we have:

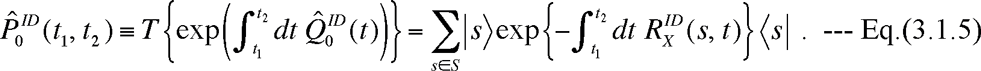

Here we omitted the explicit reminder of the sequence length dependence (*i.e.*, the superscript “(*L*(*s*))” in 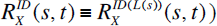, as it is obvious from the first argument, *s*. Because 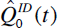 is diagonal here, the time order doesn’t matter, and the right-hand side of Eq.(3.1.5) is given by the exponentials of ordinary time-integrations. Substituting Eq.(3.1.5) into the expansion, Eq.(3.1.3), we have:

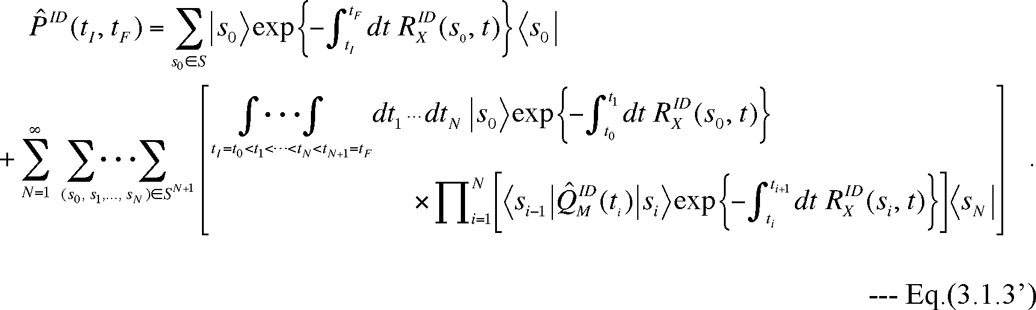

To further simplify Eq.(3.1.3’), we symbolically rewrite the definition of 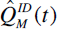, Eq.(3.1.1c) with Eqs.(2.4.2b,c), as:

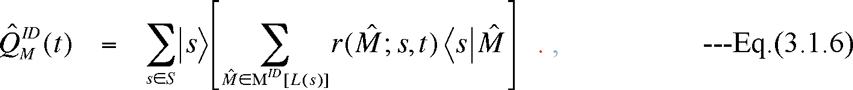

Here 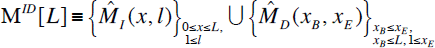 denotes the set of insertion and deletion operators that can act on the sequence of length *L*, and 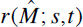 denotes the (generally time-and state-dependent) rate parameter of the indel operator 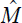 Because the action of an indel operator on a state uniquely results in another single state, we have the following identify for any function *F*(*s*) of state 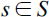:

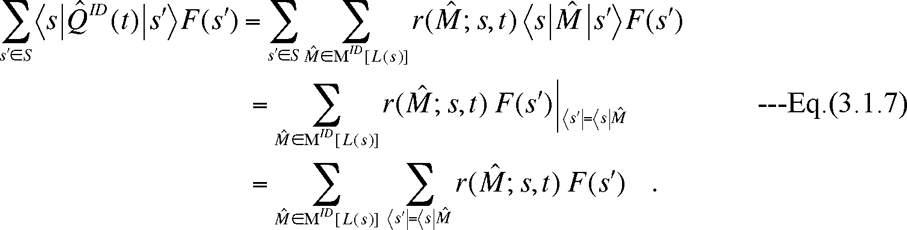

On the rightmost hand side, the single-element summation, 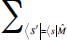, fixes the state 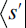 to be 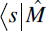. Substituting the identity, Eq.(3.1.7), into Eq.(3.1.3’), we get:

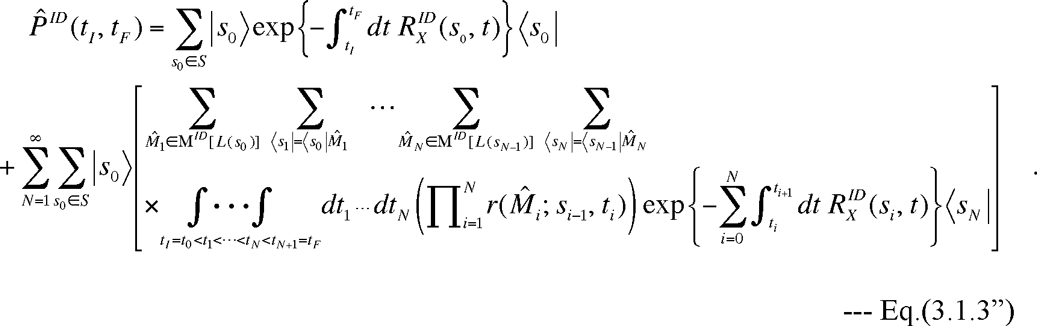

On the right hand side, the 2*N* –fold summation in the big square brackets represents the following set of recursive procedures: first sum over all possible indel operator, 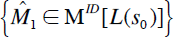, that can act on 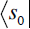; then move on to the next state 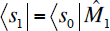 for each indel operator 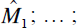 then sum over all possible indel operators, 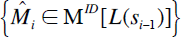, that can act on 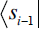; then move on to the next state 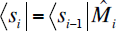 for each operator 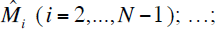 and finally reach 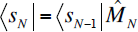 for each 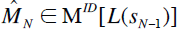. This is indeed equivalent to summing over all possible histories, 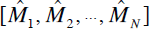, of *N* indels that begins with the state 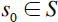, and letting the intermediate and final states uniquely determined by each history. Let 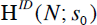 denote the space of all possible histories of *N* indels beginning with *s*_0_. Then, Eq.(3.1.3”) can be rewritten into the final expression of the perturbation expansion of the stochastic evolution operator:

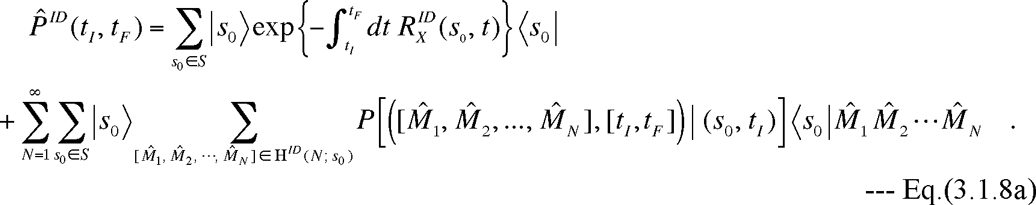

Here,

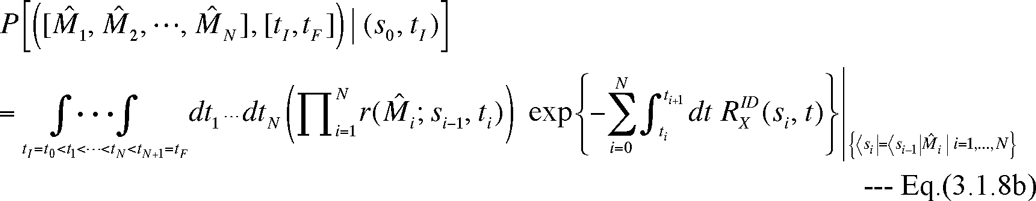

is the probability that an indel history 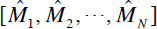 occurred during the time interval [*t_I_*, *t_F_*], given an initial sequence state *s*_0_ at time *t_I_*.

In fact, we can say that this perturbation expansion, Eqs.(3.1.8a,b), underlies the *genuine* molecular evolution simulators (Cartwright 2005; Fletcher and Yang 2009; Strope et al. 2009), which are based on the stochastic simulation algorithm proposed by Gillespie (1977). The first summation on the right-hand side of Eq.(III-1.8a) gives probabilities of the indel histories where the sequence underwent no indel events and the initial state 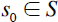 remained unchanged during the time interval [*t_I_*, *t_F_*]. Each probability, 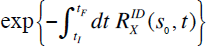, decays exactly at the exit rate, 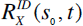, at which the state *s*_0_ undergoes an indel at time *t*. The second summation gives the probabilities of the histories where the sequence underwent at least one indel event. Let us consider, *e.g.*, an *N* –event history, 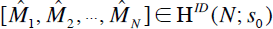. The probability of this history is given by the multiple-time integration of the probability distribution of the indel processes where the *N* indels occurred at various timings, 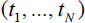 satisfying 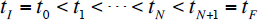. And the probability distribution of an indel process belonging to the above history is the product of the following factors (listed in temporal order): the probability, 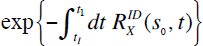, that the state 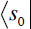 lasted from 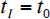 till *t_1_*; the rate, 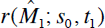, at which the event 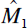 changes the state 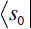 into 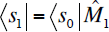 at time *t*_1_; the probability, 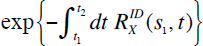, that the state 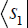 lasted from *t*_1_ till 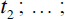 the rate, 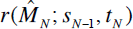, at which the event 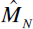 changes the state 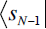 into 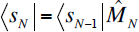 at time *t_N_*; and the probability, 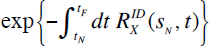, that the state 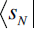 lasted from *t_N_* till 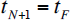. To the best of our knowledge, this study is the first to derive the explicit expression of the stochastic evolutionary operator, Eqs.(3.1.8a,b), underlying the *genuine* molecular evolution simulators, purely from the first principle (*i.e.*, the defining equation, Eqs.(2.4.8a,b), of the continuous-time Markov model of indel processes).

By sandwiching Eq.(3.1.8) with an ancestral state bra-vector 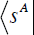 and a descendant state ket-vector 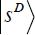 gives the conditional probability of the state 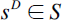 at time *t_F_* given the state 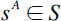 at time 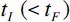: 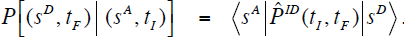.

In this case, only the contributions from indel histories consistent with the initial state *s^A^* and the final state *s^D^* will survive. Thus, letting 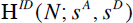 denote the set of such histories consisting of *N* indel events, we have:

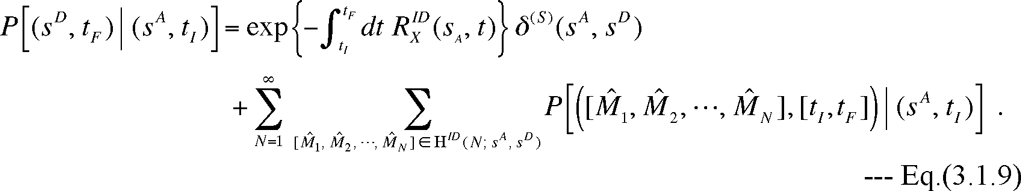

Here 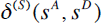 is Kronecker’s delta defined on the state space 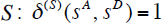 if 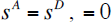 otherwise. When 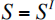, 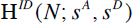 is the set of all *N*-event indel histories that change the sequence length from 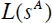 to 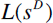. Thus, Eq.(3.1.9) is the summation of all possible alignments between the sequences of lengths 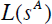 and 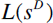. When 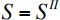, 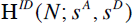 could be considered as the set of all *N*-event indel histories consistent with a given PWA between the sequences of lengths 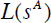 and 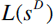, with some caveats discussed in the next subsection. When 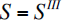, in contrast, 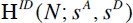 is only a subset of all *N*-event indel histories consistent with the given PWA, because 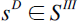 has a richer structure than necessary for merely giving the alignment with 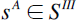. Thus, it would be convenient to introduce a separate symbol, 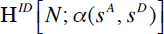, which denotes the set of *all N*-event indel histories consistent with a given PWA, 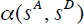, between an ancestral state 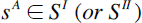 and a descendant state 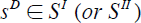. And let 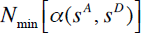 be the minimum number of indel events required to give a PWA, 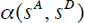. Then, we can provide the following expression for the conditional probability that 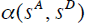 resulted during the time interval [*t_I_*, *t_F_*], given 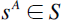 at time 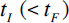:

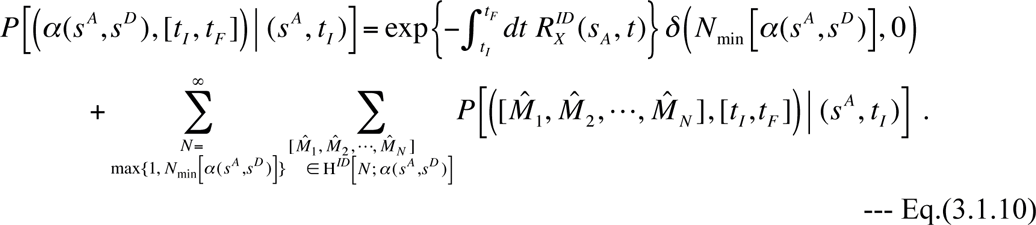

Kronecker’s delta is present in the first term because this term contributes only when 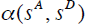 is consistent with the zero-event indel history. The conditional probability, Eq.(3.1.10), will be the building block of the probability of a given MSA, as we will see in Subsection 3.2. Finally, let

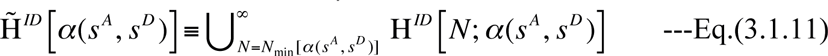

be the set of *all* global indel histories consistent with 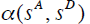, and also let

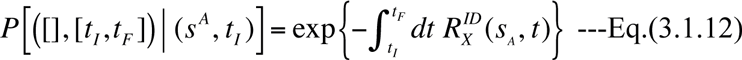

be the probability of a zero-event indel history given the ancestral state. Then, Eq.(3.1.10) can be simplified as:

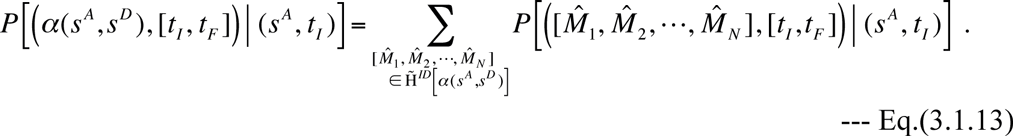

This form may be more convenient when discussing the factorization.

##### 3.1.1. Multiplicativity of perturbation expansion

An important aspect of our general continuous-time Markov model of indel processes is that, unlike any other indel probabilistic models proposed thus far (except those imposing overly simplistic restrictions on indels), it is multiplicative, that is, it satisfies the Chapman-Kolmogorov equation, Eq.(2.4.10):

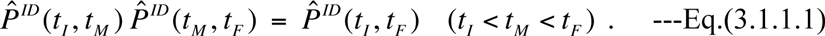

We can prove by induction that this equation is satisfied by the perturbation expansion, Eq.(3.1.3), order by order, as described in Appendix A3. This fact guarantees that our stochastic evolution operator, Eq.(3.1.3), and its more specific representation, Eq.(3.1.8), do indeed satisfy the Chapman-Kolmogorov equation, up to any desired degree of accuracy.

#### 3.2. Perturbation expansion of probability of given MSA

In Subsection 3.1, we obtained Eq.(3.1.13), which gives the probability, 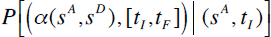, that the PWA, 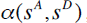, between an ancestral sequence *s^A^* and a descendant state *s^D^* resulted during the time interval [*t_I_*, *t_F_*], given *s^A^* at time *t_I_*. The right hand side of the equation is a summation of probabilities over the set, 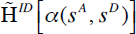, of all indel histories consistent with 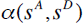. Once the probabilities of given PWAs were obtained this way, we could calculate the probability of a given MSA along the same line of thoughts as described in the introductions of Holmes and Bruno (2001) and Holmes (2003) (see also Redelings and Suchard (2005) for a superficially different but equivalent method), and we will basically follow their procedures here. We emphasize here, however, that our calculation is based purely on the continuous-time Markov model, which is a *genuine* evolutionary model of indels, as opposed to HMMs or transducer theories that past studies on indels were based on.

In this study, we formally calculate the probability of a MSA given a *rooted* phylogenetic tree, 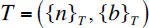, where 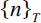 is the set of all nodes of the tree, and 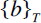 is the set of all branches of the tree. We decompose the set of all nodes as: 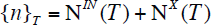, where 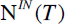 is the set of all internal nodes and 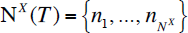 is the set of all external nodes. Here we let 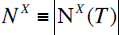 be the number of external nodes. The root node plays an important role and will be denoted as 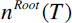, or simply 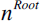. Because the tree is rooted, each branch *b* is directed. Thus, let *n^A^*(*b*) denote the “ancestral node” on the upstream end of *b*, and let *n^D^*(*b*) denote the “descendant node” on the downstream end of *b*. Let 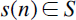 be a sequence state at the node 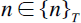, and, especially, let 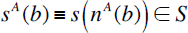 denote a sequence state at *n^A^* (*b*) and let 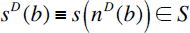 denote a sequence state at *n^D^* (*b*). Last but not least, we suppose that the branch lengths, 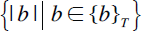, and the indel model parameters, 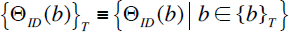, are all given. It should be noted here that the model parameters 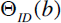 could vary depending on the branch, at least theoretically.

As supposed, *e.g.*, by Holmes and Bruno (2001), Holmes (2003), and Redelings and Suchard (2005), an indel history along a tree consists of indel histories along all branches of the tree that are interdependent, in the sense that the indel process of a branch *b* determines a sequence state *s^D^* (*b*) at its descendant node *n^D^* (*b*), on which the indel processes along its downstream branches depend. Thus, an indel history on a given root sequence state 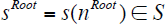 automatically determines the sequence states at all nodes, 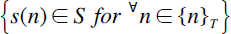. Let 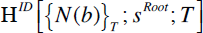 be the set of indel histories along the tree *T*. Each of its elements starts with a sequence state 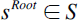 at the root and is composed of an *N*(*b*)-event indel history along each branch 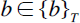. Then, a history 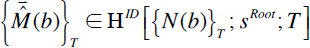 can be specified as follows:

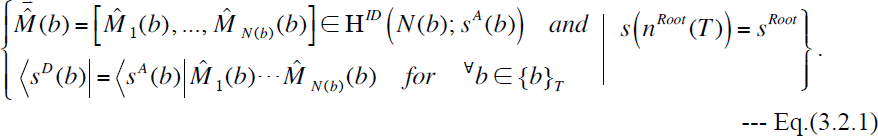

Here, as defined above Eq.(III-1.8a), 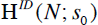 denotes the set of all *N*-event indel histories starting with the sequence state 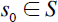. We also introduced the symbol, 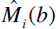, to represent the *i* th event in the indel history along the branch 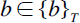. The probability of the indel history, Eq.(3.2.1), can be easily calculated. First, we already gave the probability of an indel history during the time interval [*t_I_*, *t_F_*], by Eq.(3.1.8b). Because we can correspond each branch 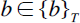 to a time interval 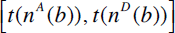 (with 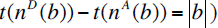, the probability of an indel history, 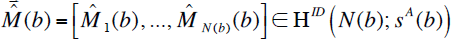), along a branch 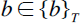 is given by:

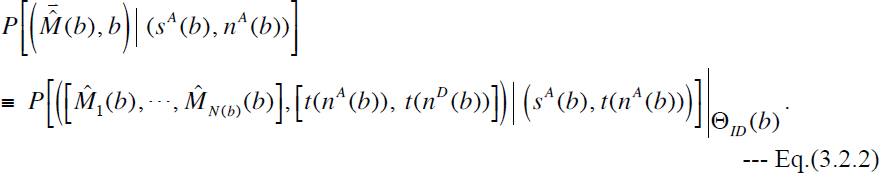

Here we explicitly showed the branch-dependence of the model parameters. Using Eq.(3.2.2) as a building block, the probability of an indel history along the tree *T*, 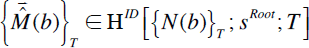, specified by Eq.(3.2.1), is given as:

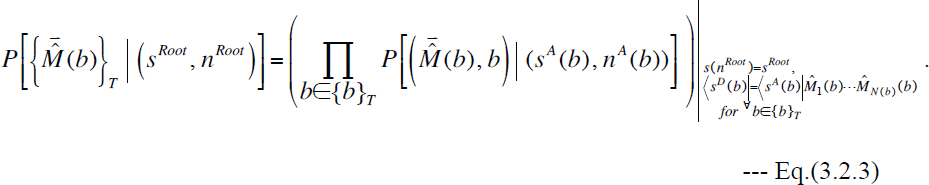

In this way, we can calculate the probability of any indel history 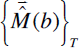 along the tree *T* starting with a given root state 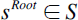. The set of all such indel histories could be expressed as:

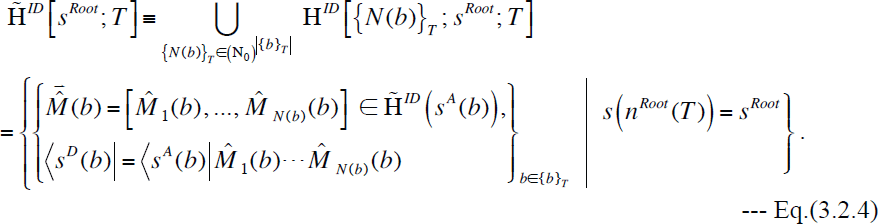

Here 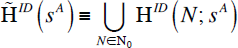 is the set of all indel histories along a branch starting with the sequence state 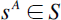.

Now, an important fact is that an indel history, along a tree starting with a root sequence state, uniquely (up to some discretional representational degrees of freedom discussed in Subsection 3.4) gives rise to a MSA, 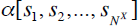, among the sequences at the external nodes, 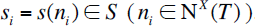. However, the converse is not true. That is, a given MSA, 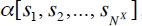, could result from a large number of indel histories along a tree, even when starting with a given sequence state at the root. (This statement will be elaborated on in Subsection 3.4.) Thus, let 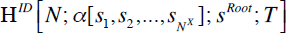 be the set of all *N*-event indel histories along the tree *T* that are consistent with the MSA 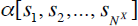 and that start with the root sequence state 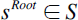. And let 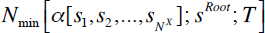 be the minimum number of events necessary for such histories. Then, under a given set of model parameters, the probability of the MSA given the phylogenetic tree and the root sequence state is formally expressed as:

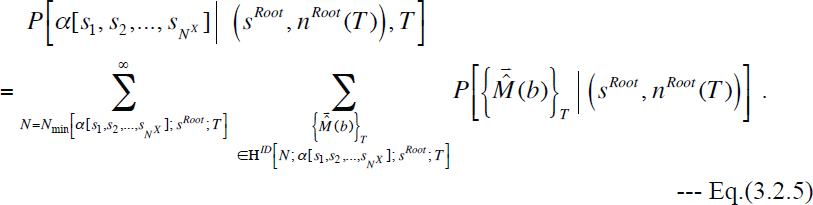

This provides a formal “perturbation expansion” of the probability of a given MSA, *conditioned on* a given root sequence state. To give the *unconditioned* probability of the MSA, 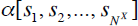, we multiply Eq.(3.2.5) with the probability of *s^Root^*, and sum over the set, 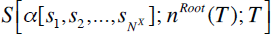, of all possible root sequence states consistent with the MSA:

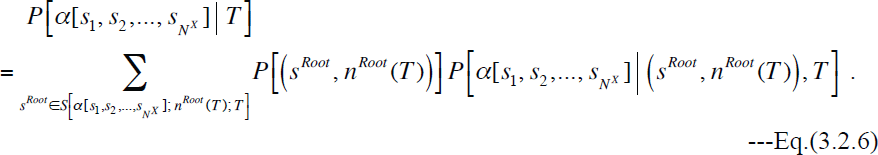

Here 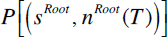 denotes the probability of the sequence state *s^Root^* at the node 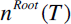. Because we allow for non-equilibrium evolution in general, we regard the probability of a sequence state as a function of the point on the tree (under the given phylogenetic tree and model parameters). It would probably be more convenient to rewrite the combination of Eq.(3.2.6) and Eq.(3.2.5) so that the summation over the number of events will be outermost. For this purpose, we introduce the space of pairs, each of a root sequence state and an *N*-event indel history starting with the root state, that are consistent with the MSA:

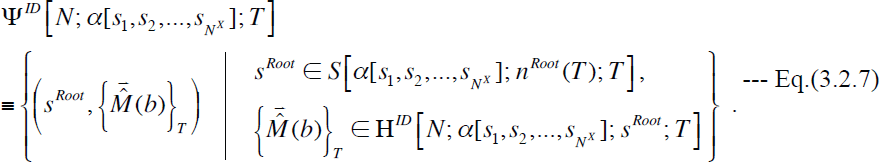

And we also introduce the *unconditioned* minimum number of indel events necessary to produce the MSA:

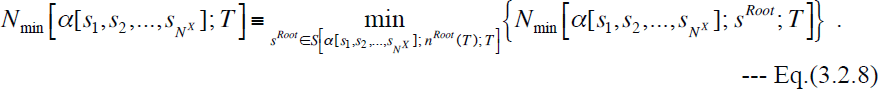

Then, the combination of Eq.(3.2.6) and Eq.(3.2.5) can be rewritten as:

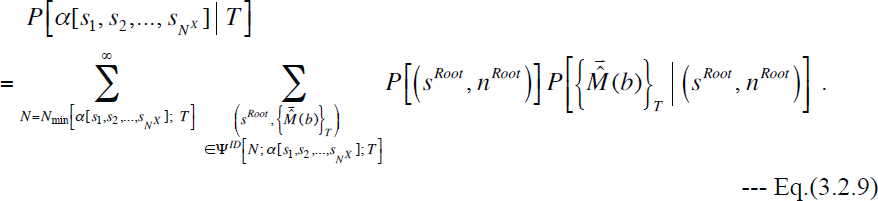

This is the formal “perturbation expansion” of the *unconditioned* probability of a given MSA. We consider this more convenient because we usually search for the indel histories and the root sequence states simultaneously. Or rather, the latter are usually given as a consequence of the search for the former. It would be very rare, if at all, in a practical analysis to give the root sequence state first and then give the indel histories on it. By introducing the set of all pairs consistent with the MSA:

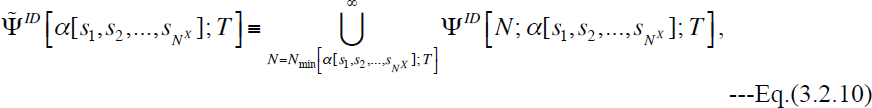

Eq.(3.2.9) could be rewritten in a more compact form:

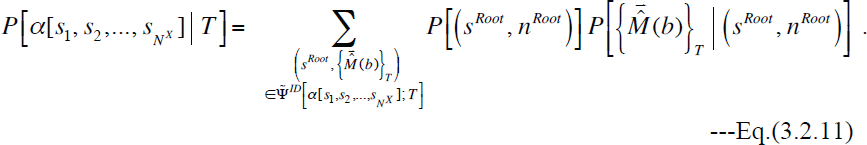

This facilitates the “decomposition” of the unconditioned probability, 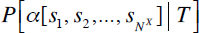, in different ways than in Eq.(3.2.9). For example, let 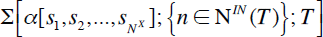 be the set of all sets, each of which consists of sequence states at all internal nodes, *i.e*, 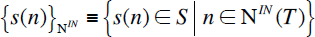 that collectively are consistent with 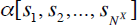. And let 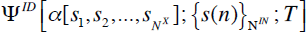 be the set of all indel histories that are consistent with both 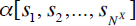 and 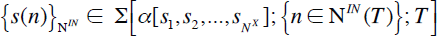. Then, 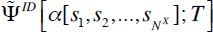 can be decomposed differently from Eq.(3.2.10) as:

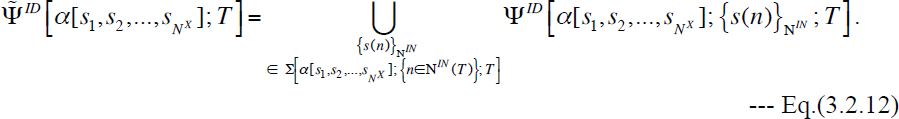

Thus, we get:

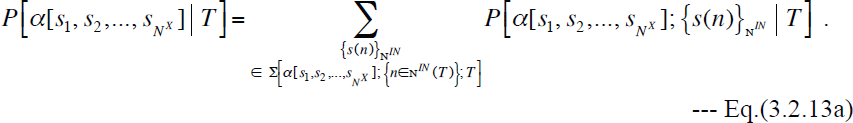

Here

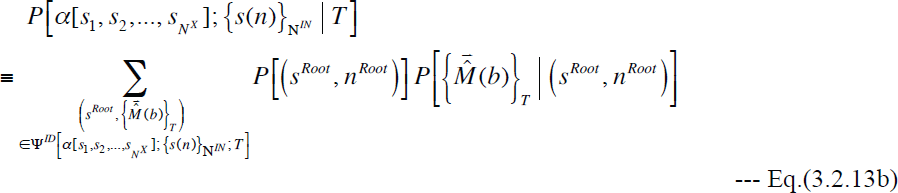

is the probability of simultaneously getting a MSA, 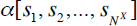, and a consistent set of states at internal nodes, 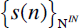. Eq.(3.2.13b) is a summation of the contributions from indel histories consistent with a specified 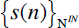. If we work in the state space *S^II^*, a particular set, 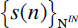, uniquely determines a pairwise alignment between sequence states at both ends of each branch (again up to the discretional representational degrees of freedom discussed in Subsection 3.3). Thus, taking account of Eqs.(3.2.1,3), Eq.(3.2.13b) could be further re-expressed as a product of 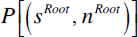 and the probabilities of such pairwise alignments:

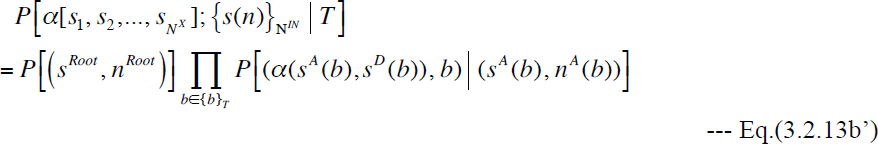

Each

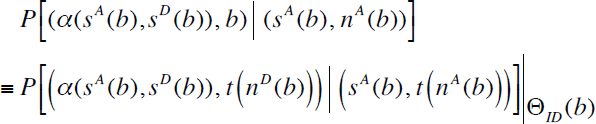

in the right-hand side of Eq.(3.2.13b’) can be calculated by using, *e.g.*, Eq.(3.1.13). This expression, Eq.(3.2.13a) accompanied by Eq.(3.2.13b’), is most in line with those proposed in the introductions of Holmes and Bruno (2001) and Holmes (2003). Another way to “decompose” 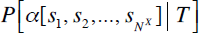 in Eq.(3.2.11) will be given in Subsection 4.2.

Let us now consider the set, 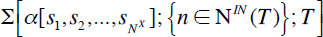 of sets of sequence states at all internal nodes consistent with a MSA, 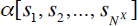. The union of these sets over states at internal nodes other than the root gives 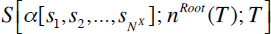, the set of root sequence states consistent with the MSA. So, we will only consider what 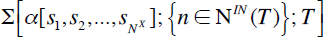 is like. An important clue comes from the fact that each column of a MSA accommodates only those sites descended from the same ancestral site, or gaps for sequences lacking such sites. It leads to the “phylogenetic correctness” condition that the MSA-consistent internal sequence states must satisfy (Chindelevitch et al. 2006; Diallo et al. 2007). The condition could be rephrased to fit the current context as: “if a site corresponding to a MSA column is present in the sequence states at two nodes of the tree, then, the site must also be present in all sequence states along the path connecting the two nodes.” (See panels A and B of Figure 4.) This condition not only restricts the sequence state at an internal node given the states at all external nodes (i.e., a MSA column), but also restricts possible interrelationships between the states at different internal nodes. More precisely, the nodes, both external and internal, with sequence states containing a particular site (more precisely, a site of a particular ancestry) must always form a single, connected “web” in the tree, which contains no external nodes without the site (Figure 4, panels A, B). This substantially limits the possible state configurations at each MSA column (panels C, D, E, F), and it helps explore possible indel histories in a reasonable amount of time in most practical cases (as argued, *e.g.*, by Kim and Sinha (2007)).

**Figure 4.**
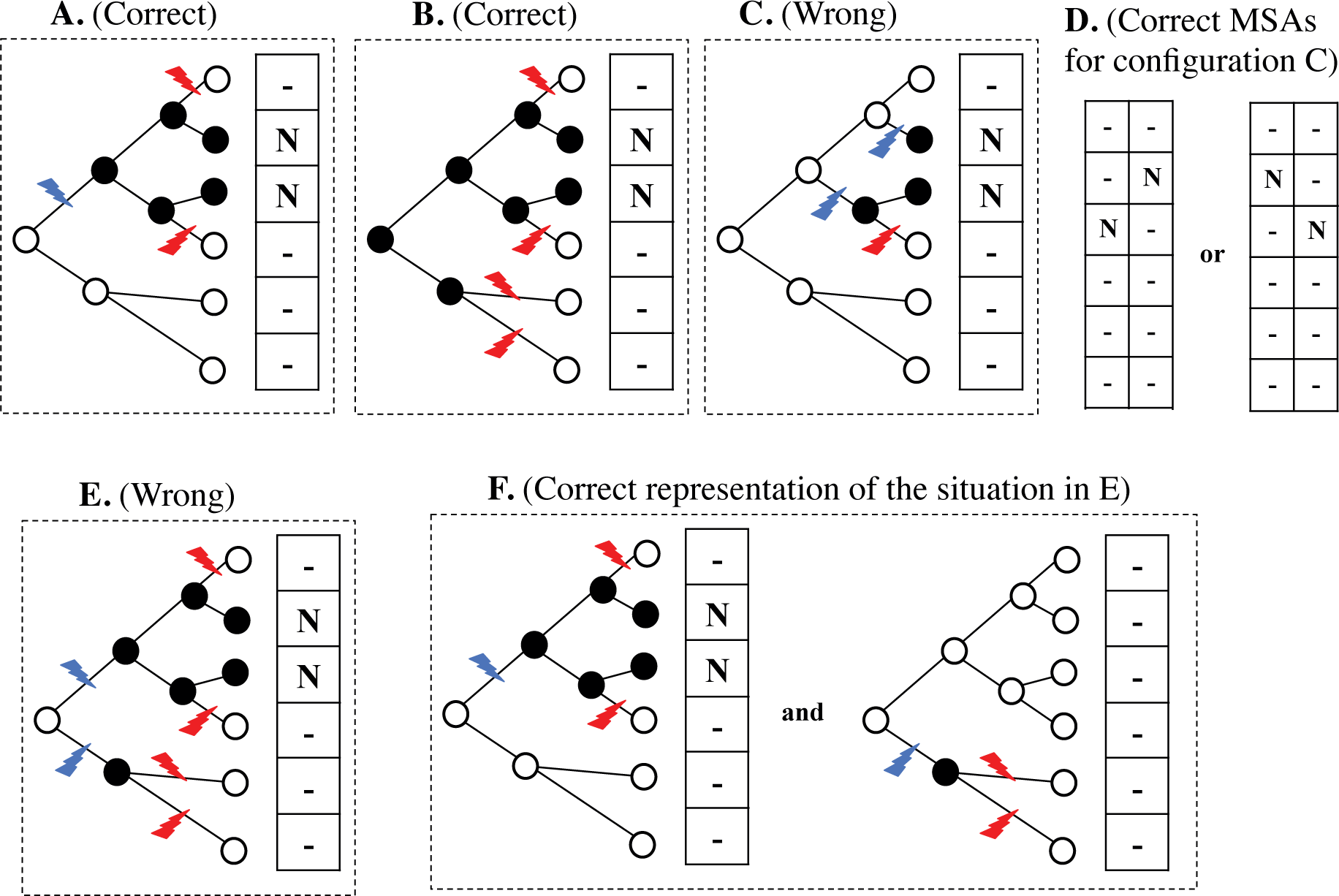
Phylogenetic correctness condition. **A.** This condition states that, if two sequences in a MSA hold homologous sites (the “N”s aligned in the column), all internal nodes along the path connecting the two sequences must also hold the site (the filled circles on the tree). **B.** Another phylogenetically correct configuration, which forms a single, connected “web” of nodes (and branches) holding the site. **C.** A phylogenetically wrong configuration, where there are two mutually disconnected “web”s, indicating two independent insertions (, one of which was followed by a deletion). Such a history must give rise to two independent columns, as in panel **D**. **E.** Another phylogenetically wrong configuration, which must be represented as two separate configurations, as in panel **F.** Each of the panels A, B, C and E consists of a tree and a MSA column enclosed by a dashed box. The ‘-‘ in each column represents a gap, meaning that the site is absent. The open circle in each tree represents the absence of the site from the sequence at the node. The red and the blue lightning bolts represent a deletion and an insertion, respectively.

#### 3.3. Equivalence classes of indel histories during time interval (II)

In Subsection 2.3, we introduced the local-history-set (LHS) equivalence between *global* indel histories as a set of histories that can be derived from the same set of local indel histories, through the unary equivalence relations, Eqs.(2.3.1a,b,c), and the binary equivalence relations, Eqs.(2.3.3a-d). A PWA cannot tell the relative time order between indel events in separate local histories. Therefore, if a global indel history can give rise to a given PWA, so can any other global histories that are LHS equivalent to it. Thus, the set, 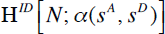, of all global histories of 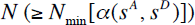 indel events consistent with a PWA, 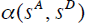, must be a union of (mutually disjoint) LHS equivalence classes of *N*-event histories consistent with 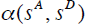. As discussed in Subsection 2.3, each LHS equivalence class can be represented by a set of local indel histories (local history set (LHS)), *e.g.*, 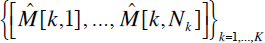. Here, let 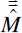 be a shorthand notation of such a LHS.

Let 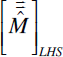 be the LHS equivalence class represented by 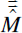. And let 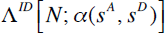 be the set of all local indel history sets that are consistent with 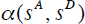 and each of which is made up of *N* events in total (*i.e.*, satisfying 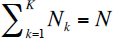 in the above example). Then, we have:

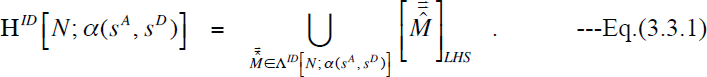

Next, let 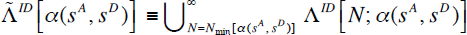 denote the set of *all* local indel history sets consistent with 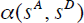. Then, from Eq.(3.3.1), we have, for the set of all global indel histories consistent with the PWA:

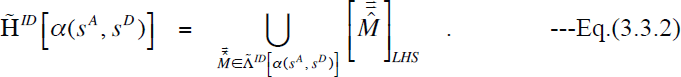

Thus the set of all global indel histories consistent with the PWA, 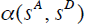, can be decomposed into the union of LHS equivalence classes. Next we compare different LHS equivalence classes that are components of 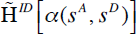. Because each equivalence class is represented by a set of local indel histories, we can focus on the differences between local indel histories that acted on the same region of the ancestral sequence, delimited either by a pair of preserved ancestral sites (PASs) or by 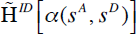 a PAS and a sequence end. By definition, all components of must give the same alignment, 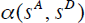. Thus, the local indel histories under comparison must also give the same *local* alignment, which must be a sub-region of 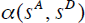 delimited in the same way as the local indel histories. Hence, in this sense, the local histories in question are equivalent. We already conjectured in Subsection 2.3 that such local histories must be connected with each other through a series of equivalence relations involving *overlapping* indel operators (non-exhaustively given in Appendix A1), and maybe additionally of some binary relations, Eq.(3.3.3a-d). Among them, we think that two cases require particular attentions: local histories that leaves no traces in the PWA, and histories that highlights the difference between the set of homology structures (see, e.g., Lunter et al. 2005) and the alignment of sequences as linear arrays of sites.

First, a series of events could have occurred between two adjacent PASs in a PWA, if it left no traces in either the ancestral or the descendant sequence. Such a local indel history needs to have started with an insertion between the PASs and ended with a deletion of everything that had been created in between (and excluding) them (see, e.g., Figure 3H). Thus, in order to estimate the probability of a PWA very accurately, such “null local histories” need also be taken into account in between each pair of PASs. Once the alignment probability is proven to be factorable, we can calculate the contributions of such null histories independently for each inter-site position, and thus the computation will be simplified considerably. To the best of our knowledge, no references thus far have *explicitly* discussed the effects of such null histories on the probability of a pairwise alignment. But it is almost certain that, although *implicitly*, the exact solutions of simple models (e.g., Thorne et al. 1991, 1992; Miklós and Toroczkai 2001) and the approximate likelihood of the “long indel” model (Miklós et al. 2004) incorporated this factor.

Second, consider the local PWA resulting, e.g., from a local history, 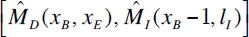. In this history (Figure 5A), a sub-sequence between (and including) the *x_B_* th and *x_E_* th sites is first deleted, and a sub-sequence of length *l_I_* is inserted exactly between the sites that flanked the deletion. This history could be represented by two alternative PWAs (panels B and C of Figure 5), because there is no *a priori* way to specify the relative spatial positioning of the deleted and inserted subsequences in this case. However, these two PWAs could also result from other local histories, different from the aforementioned one and also from each other; for example, Figure 5B could result also from 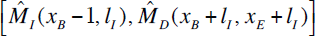 (Figure 5D), and Figure 5C could result also from 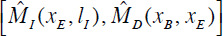 (Figure 5E).

**Figure 5.**
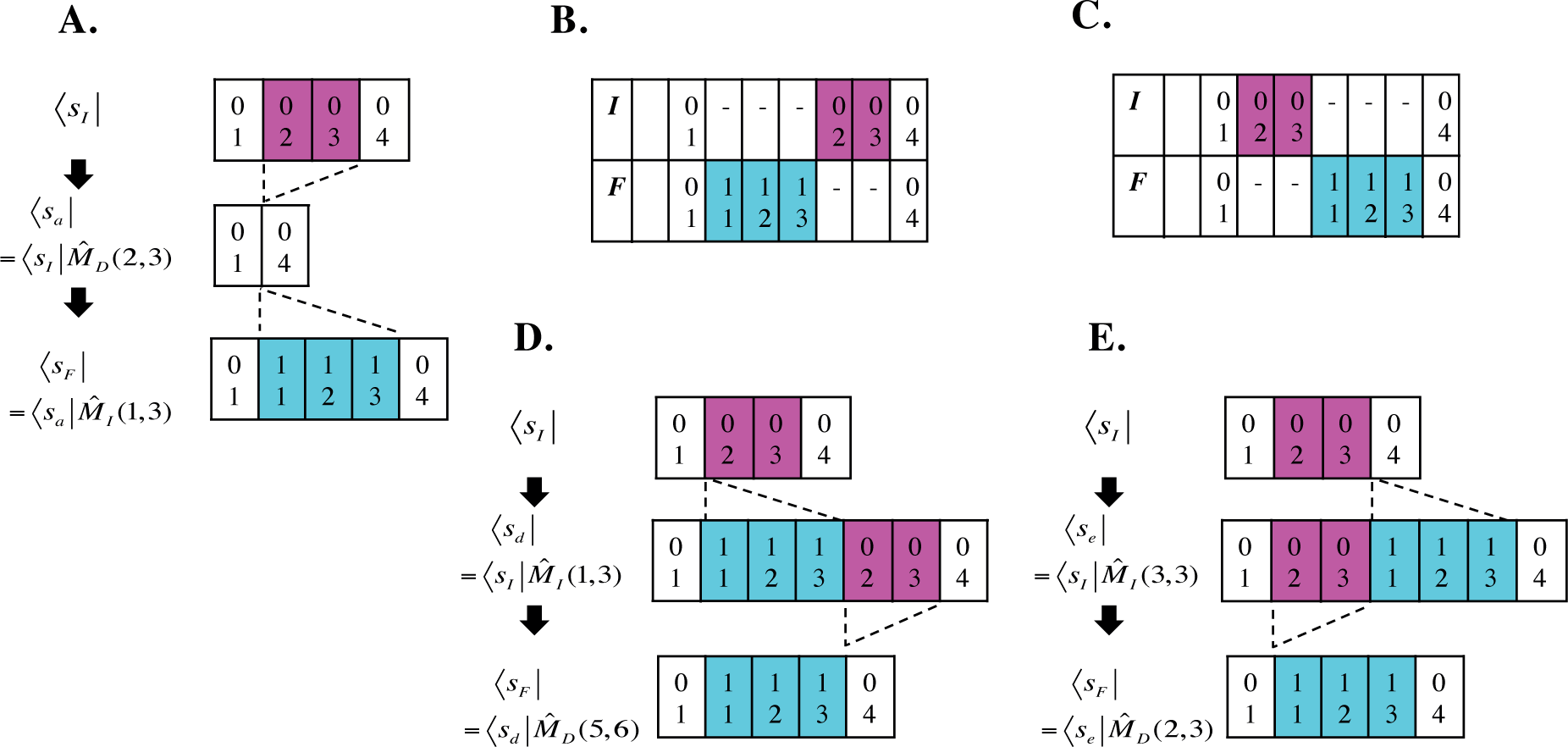
Ambiguities in interpretation of PWA. **A.** A deletion (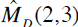) followed by an insertion between the deletion-flanking sites (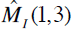). Two alternative PWAs that could result from this indel history are shown in panels **B** and **C**. **D.** An alternative indel history, 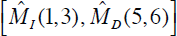, that can result in the PWA in panel B. **E.** An alternative indel history, 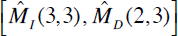, that can give rise to the PWA in panel C. We followed the same notations as in Figure 2.

These two local histories result in different intermediate states. Each of them have both inserted and deleted subsequences. However, the states in panels D and E have the inserted subsequence on the left and on the right, respectively, of the deleted subsequence. (For similar equivalence relations involving overlapping indels, see panels C, F, and G of Figure 3.) This difference might become important when discussing, e.g., possible functions of the intermediate sequences. Although these examples were on parsimonious indel histories, similar problems arise, likely more frequently, when we deal with non-parsimonious indel histories. Consider, *e.g.*, a three-event indel history, 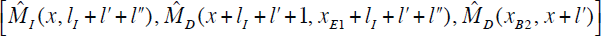 (with 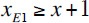, 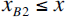 and 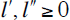 This history results in an inserted subsequence of length *l_1_* flanked from the left by a deletion between (and including) the *x*_*B*2_ th and *x* th ancestral sites and flanked from the right by a deletion between (and including) the *x*+1 th and *x*_*E*1_ th ancestral sites (panel A of Figure 6). Such positional relationships among indels would be revealed by the output of a simulator that faithfully records the actions of indels and their effects on the sequence states (Figure 6, panel B). However, even if we work in the space *S^III^*, just comparing the ancestral and descendant sequence states will never reveal such a linear structure among the responsible indels. Instead, it indicates that 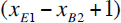 sites and *l_I_* sites are only in the ancestral state and the descendant state, respectively, in between a pair of neighboring (but not contiguous) PASs. In this situation, it is currently common to “parsimoniously” interpret it as, *e.g.*, a run of *l_I_* sites only in the ancestor followed by a run of 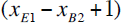 sites only in the descendant (panel C of Figure 6), which can be interpreted with histories with fewer indels (panels D, E, F), ignoring the possible histories as described above (panel A) (and accompanying intermediate sequence states including functions). Thus, depending on the situations (including parameter values), ignoring these issues could cause the probabilities of local PWAs to be misestimated. Thus far, it seems to have been a common practice to arrange or rearrange a set of inserted sites and a set of deleted sites into two blocks according to a pre-fixed order when inferring an optimum PWA or calculating the probabilities of possible PWAs from an input pair of sequences. We suppose that such a common practice is inevitable, considering that such arranged PWAs (e.g., panel C of Figure 6) are in general more likely than, *e.g.*, PWAs with an alternating run of multiple inserted and deleted segments uninterrupted by PASs (e.g., panel B). And such “parsimonious” interpretations should considerably save computational time and memory in general. Nevertheless, at least when interpreting the results of the analyses, it would be better to take account of the possibilities exemplified above, in order to avoid possibly serious errors. Similar issues arise also for indel histories giving rise to a MSA, as we will see in the next subsection.

**Figure 6.**
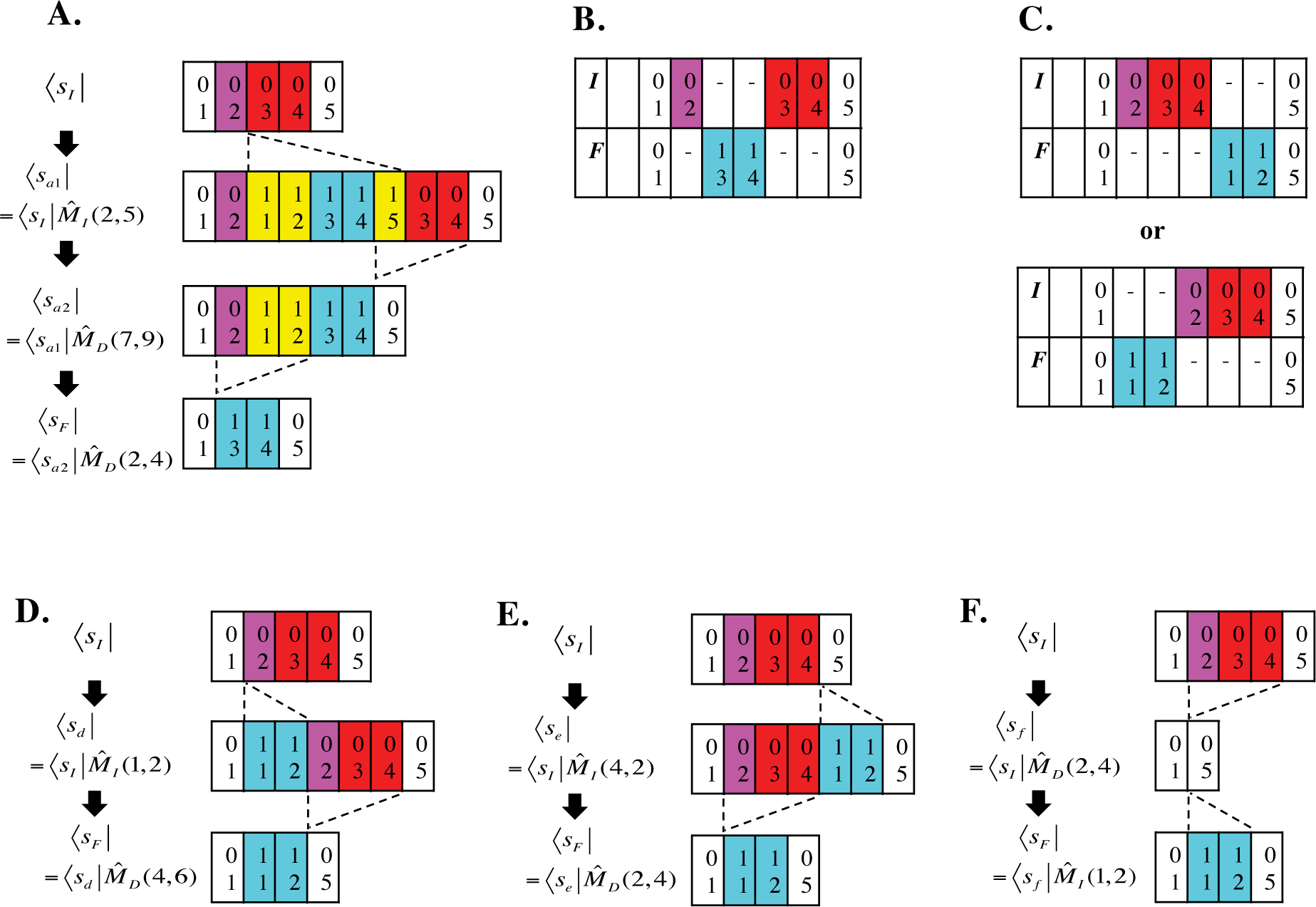
PWA representations of somewhat complex indel history. **A.** An example 3-event history, 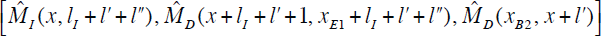, with 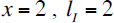, 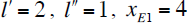, and 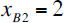. **B.** A PWA that would be output by a simulator that faithfully records the actually occurred indels and their outcomes. **C.** Two alternative “parsimonious” PWAs that would commonly be output by existing alignment programs, when the history in panel A actually occurred. **D, E**, and **F.** Three parsimonious interpretations of both of the PWAs in panel C.

#### 3.4. Equivalence classes of indel histories along phylogenetic tree

Here we will consider indel histories along a phylogenetic tree (including the initial sequence state at the root), 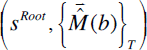’s, that are equivalent in the sense that they give the same MSA, 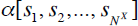. The largest such equivalence class would be 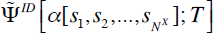, the set of all histories consistent with the MSA (in Eq.(3.2.10)). Here we consider some of its typical subsets that will help our theoretical calculations in the following sections. First, the concept of the local-history-set (LHS) equivalence could be extended to indel histories along a tree. As discussed in Subsection 3.2, an indel history along a tree is, after all, a set of histories along all branches, interdependent from the root down to the leaves (see Eq.(3.2.1)). Given a sequence state at the ancestral node, all histories belonging to a LHS equivalence class along each branch gives the same sequence structure at the descendant node, including the features that cannot be captured by PWAs output by commonly used aligners. Therefore, if we give particular LHS equivalence classes along all branches of the tree, as well as a particular root sequence state, they will result in a unique set of sequence state structures at the leaves of the tree, including a MSA of the sequences at the leaves. Thus, we define a LHS equivalence class *along a* tree, 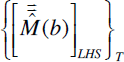, on a given root state 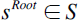 as follows:

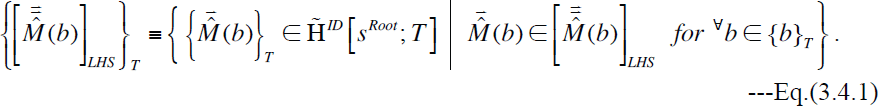

Here 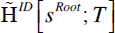 is the set of *all* indel histories along the tree *T* starting with the root state 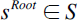 (see Eq.(3.2.4)). Using such equivalence classes, we can decompose 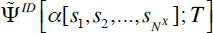. For this purpose, let 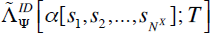 be the set of all pairs, each of which is composed of a root state 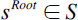 and a set, 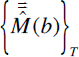, of local history sets along all branches, that are consistent with 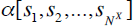. Then, similarly to Eq.(3.3.2), we have:

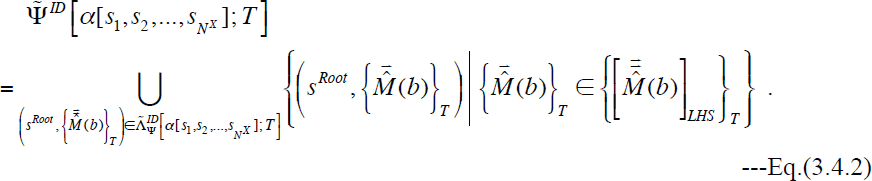

The equivalence through relations involving overlapping indels (*e.g.*, those in Appendix A1) also naturally defines the equivalence among histories along a tree, if we apply the relations to indel events along the same branch. More nontrivial relations are equivalence relations involving events along different branches. A clue comes from Eq.(3.2.12), which decomposes 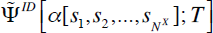 into a union of disjoint subsets over 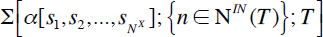, composed of sets of states at internal nodes consistent with 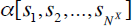. Broadly speaking, equivalence relations within each subset of 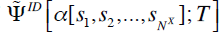 consistent with an element in 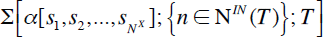 are covered by the LHS equivalence and other equivalences involving only indels along the same branch. Thus, all we have to explore here are indel histories giving rise to different sets of states at internal nodes consistent with 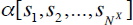. In Subsection 3.2, we also explained that, for sequence states at internal nodes to be consistent with 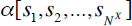, each site must be present in a set of nodes that form a *single* connected “web” in the tree (see, e.g., Figure 4), in order to satisfy the “phylogenetic correctness” condition (Chindelevitch et al. 2006; Diallo et al. 2007). Thus, by changing the states at internal nodes while keeping the web to be single and connected (i.e., while keeping it from splitting into two pieces) for each site, and by giving indel histories consistent with such states, we can move between histories via this new category of equivalence relations (e.g., Figure 7). Another important kind of move is to add or remove a “null local indel history” along the tree, which is consistent with a single, connected web consisting of at least one internal node but no external nodes (Figure 8). As far as we know, Rivas and Eddy (2008) were the first to explicitly consider these null local indel histories along the tree when calculating the probability of a MSA given a tree, albeit under a single-residue indel model. In our general continuous-time Markov model of indels, a (run of) column(s) corresponding to such a web with no external nodes could be joined with a run of gapped columns or flanking runs of gapped columns. This could enrich the repertoire of non-parsimonious local indel histories possibly responsible for the local MSA (Figure 9).

**Figure 7.**
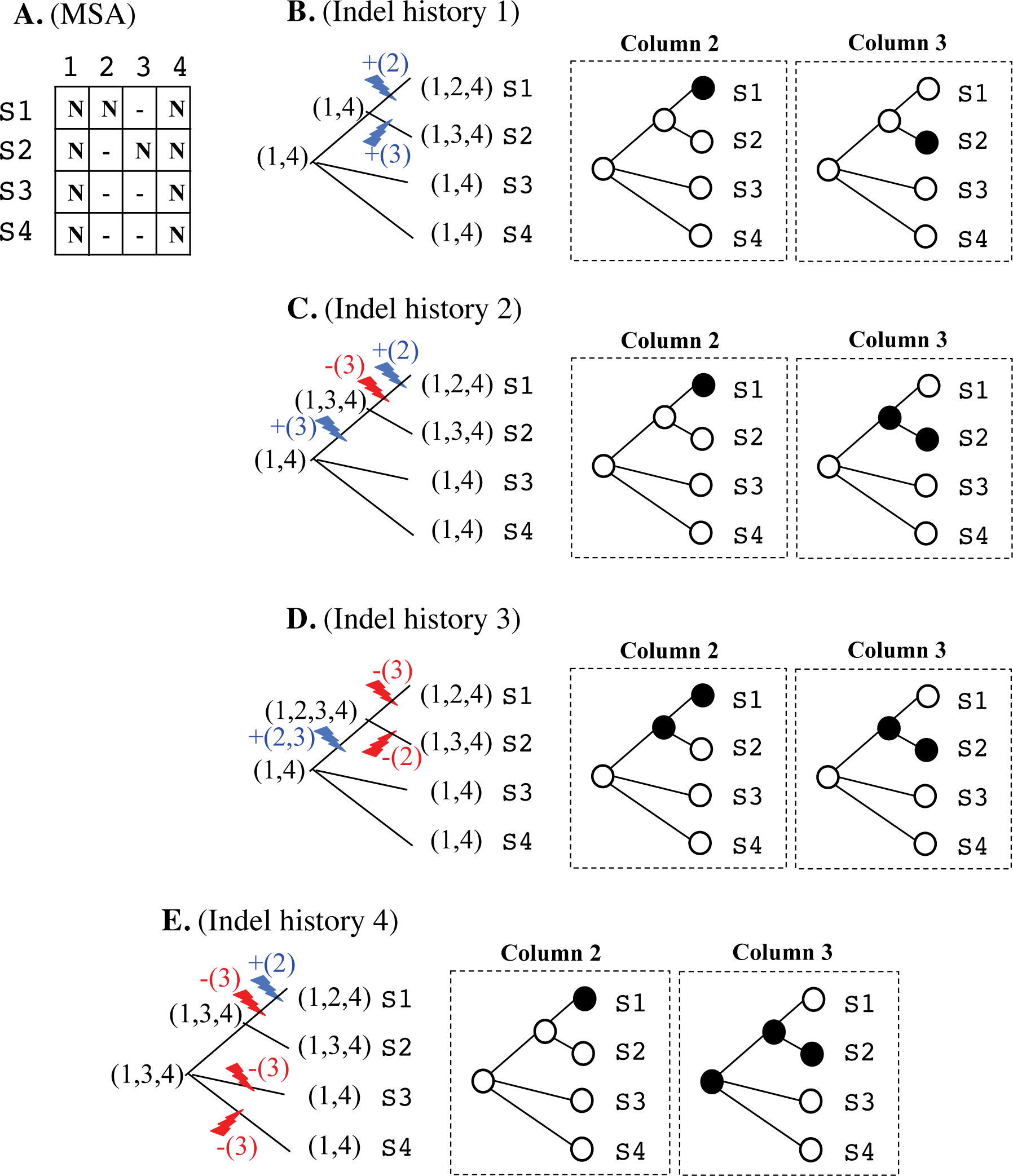
Equivalence relations between (local) indel histories along tree. Given a MSA [panel **A**], we can conceive of some indel histories consistent with it [shown, *e.g.*,in panels **B, C, D** and **E**]. These indel histories are equivalent, in the sense that they give rise to the same MSA. Each of panels **B, C, D** and **E** consists of an indel history mapped on the left tree, and two other trees (middle and right) showing whether the site corresponding to each MSA column is present in (a filled circle) or absent from (an open circle) the sequence at each node. In the left tree, the sequence state at each node is represented by the parenthesized list of MSA columns present in the sequence. And the set of blue/red parenthesized numbers, + / - (*x, y, z*), accompanying each blue/red lightning bolt represents the set of MSA columns inserted/deleted by the event. The move between the histories can be interpreted as a contraction or extension of the “web” of nodes (and branches) possessing each site, which changes the ancestral states at the internal nodes. Such a “web” transformation could be accompanied by an equivalence move between histories along each relevant branch as exemplified in Appendix A1 (not shown here). In all panels, S1-S4 are the sequence names, and the MSA columns are numbered 1-4.

**Figure 8.**
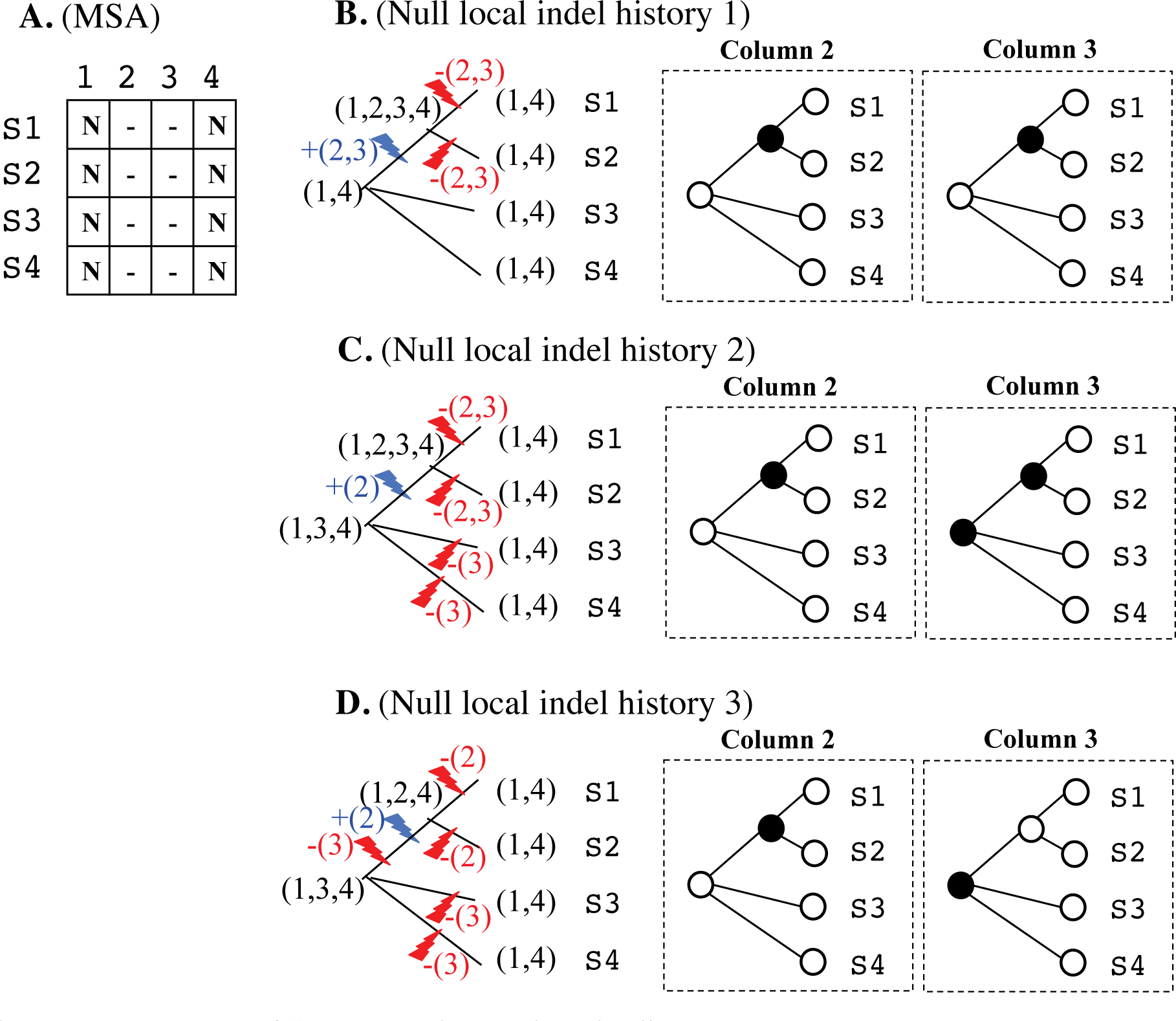
Examples of “null local indel histories.” **A.** Here, for clarity, the effect of null local indel histories is represented with a block of contiguous “null” MSA columns, each of which consists only of gaps (columns 2 and 3). **B, C, D.** Examples of null local indel histories that could give rise to the null MSA columns in panel A. This figure uses the same notation as Figure 7 does.

**Figure 9.**
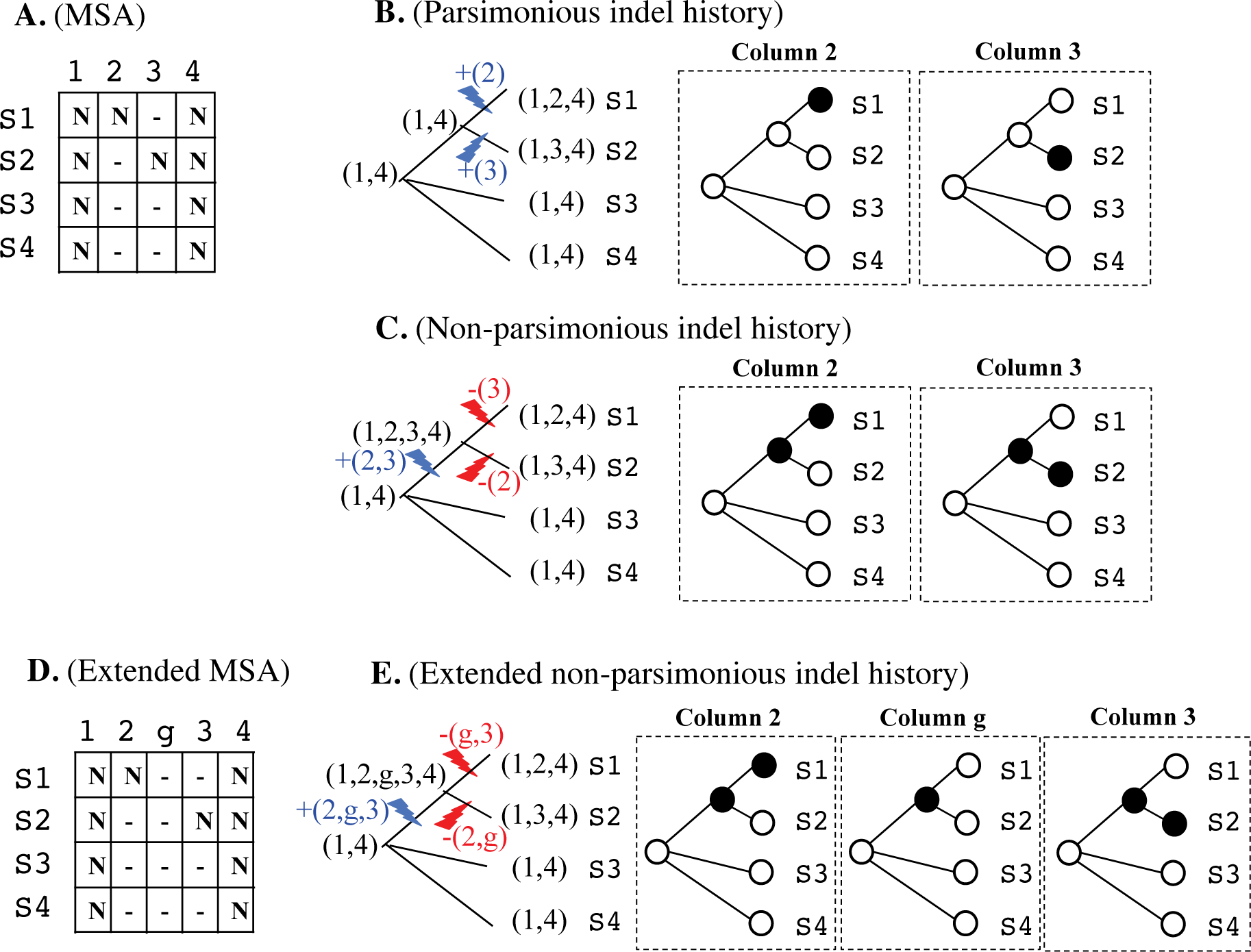
Enriched repertoire of non-parsimonious local indel histories by null local indel histories. Usually, the MSA in panel **A** most likely resulted from the parsimonious local indel history in panel **B.** Although the non-parsimonious local indel history in panel **C** could also give rise to the MSA in panel A, usually it is much less likely. However, if we notice that the extended MSA in panel **D** is also equivalent to that in panel A, we also notice that another class of non-parsimonious local indel histories shown in panel **E** could also result in the MSA. This could enhance the total likelihood that this class of non-parsimonious local indel histories is responsible for the MSA. In panels D and E, the ID “g” is assigned to the gap-only column, to facilitate the comparison between histories C and E.

In a MSA, a gapless column corresponds to a preserved ancestral site (PAS) in a PWA, because the existence of a gapless column means that the site was preserved in all compared sequences. Thus, by the “phylogenetic correctness” condition, a gapless column indicates that no indel events struck or penetrated the site throughout the evolutionary history along the phylogenetic tree. Hence, indel events that occurred in regions separated by more than one gapless column will never *physically* interfere with each other. This constraint enables us to deal with these events separately when considering the indel histories along the tree. However, this does not necessarily mean that we can always deal with them separately when calculating the probability of the indel histories. In Subsection 4.2, we will see under what conditions we can separate the contributions of such events.

### 4. Factorization of alignment probability

In the last section, we expressed the probability of a PWA and that of a MSA in perturbation expansions. These formulas, e.g., Eq.(3.1.10) for a PWA and Eq.3.2.11) for a MSA, could be immediately used to calculate the probability when the total number of indels along a branch (or, equivalently, during a time-interval) is at most, *e.g.*, ten. As they are, however, these formulas will be practically useless when there are more non-overlapping indels along a branch, in which case the probabilities must be summed over at least, e.g., 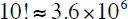 indel histories in the same LHS equivalence class. It would thus be convenient if the alignment probability can be factorized into a product of contributions from blocks (or segments) separated by preserved ancestral sites (PASs), even if it cannot be factorized into a product of column-wise contributions as in most HMMs or transducers. Such factorization has an additional benefit of potentially preventing a combinatorial explosion due to contributions from non-LHS equivalent indel histories. Miklós et al. (2004) conjectured a similar factorization when they calculate the probability of a given PWA under their “long-indel” model, but they did not explicitly prove it. Here, starting from Eq.(3.1.13) for a PWA probability under the general continuous-time Markov model describing the evolution of an *entire* sequence via insertions and deletions, we will examine whether and how the probability can indeed be factorized. We will also examine the conditions on the indel rate parameters under which the probability is factorable.

#### 4.1 Factorization of probability of PWA between descendant and ancestral sequences

Let us re-examine Eq.(3.1.13) for the probability of a given PWA, 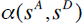, conditioned on an ancestral state, *s^A^*. Because we are interested only in whether it is factorable or not, the indel histories giving rise to 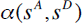 are assumed to contain at least two indel events separated by at least a PAS. It is immediately obvious that each component probability, 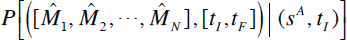, given by Eq.(3.1.8b), will *not* be factorable. This is because the multiple-time integral is over the region, 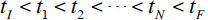, which *cannot* be expressed as a direct product of two or more regions. As mentioned in Subsection 3.3, however, each indel history, 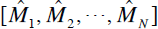 belongs to a LHS equivalence class, 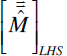 represented by a set of local indel histories, *e.g.*, 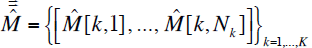, that satisfies the equivalence, 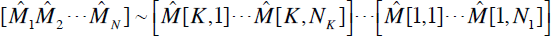, only through the binary equivalences Eqs.(3.3.3a-d) (and possibly the unary equivalences Eqs.(3.3.1a-c)). And the entire set 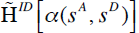, over which the summation in Eq.(3.1.13) is performed, was decomposed into the union of LHS equivalence classes in Eq.(3.3.2). Thus, we should prove the factorability of the PWA probability, Eq.(3.1.13), broadly in the following two steps. (i) Prove, under a certain set of conditions, the equation:

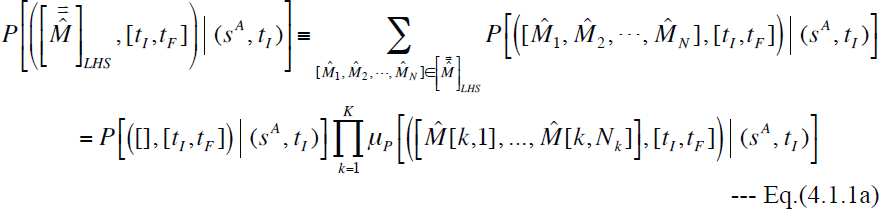

for each equivalence class definition: 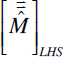 (with 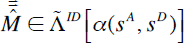). Here we used the

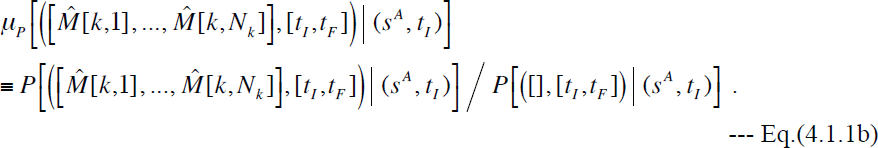

And (ii) put together Eq.(4.1.1a) over the set 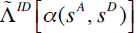 of LHS-equivalence classes, and lump together contributions from each of different regions separated by PASs in the PWA. Once step (i) is achieved, step (ii) is almost trivial. Thus, we will concentrate our efforts on finding out a set of conditions that enables the factorization, Eq.(4.1.1a). Actually, this two-step form of factorization, with the associated proof of Eq.(4.1.1a) given below, may be too restrictive compared to a possibly more general factorization of 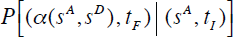. In general, Eq.(4.1.1a), or its derivative equations to be proved below, will not necessarily hold and yet the probability may be factorized via an intricate and miraculous cancellation among the contributions from indel histories in the same LHS-equivalence class, or even among contributions from different LHS-equivalence classes. In this sense, the conditions that we will find are regarded as “sufficient and *nearly* necessary” for the factorization. However, we believe that such “more general” factorizations, if at all, will be isolated exceptions, and that our proof will be general enough in practice.

Now we start proving Eq.(4.1.1a). First, substituting Eq.(3.1.8b) with some modifications into the rightmost side of Eq.(4.1.1a) divided by 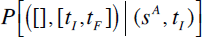, we have:

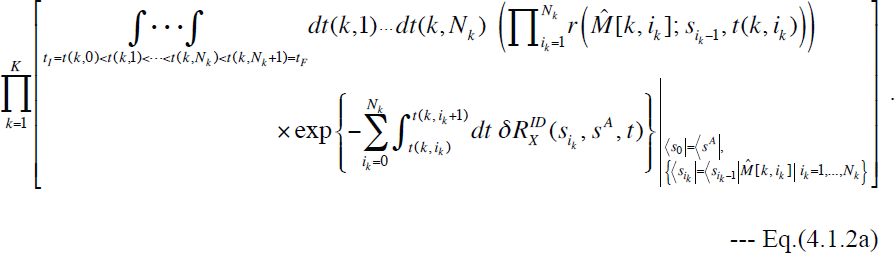

Here, 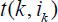 denotes the time at which the event 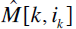 virtually occurred in the isolated *k* th local history, and we used a shorthand notation, 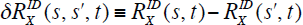. Second, we note that each LHS equivalence class, 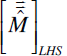, consists of 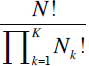 global indel histories. Each history corresponds to a map from each event in each local indel history (specified by *k*) to a temporal order within the global history:

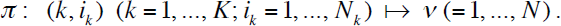

The map keeps the relative temporal order among indels in each local indel history. Then, 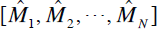 in the middle of Eq.(IV-1.1a) corresponding to the above *π* can be more precisely written as: 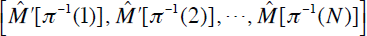. Here 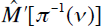 is an equivalent of 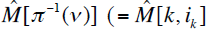 for 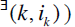 through a series of Eqs.(2.3.3a-d) to rearrange the events in 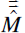 this way. Now, let 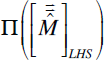 be the set of such 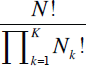 maps. Then, the expression in the middle of Eq.(4.1.1a) divided by 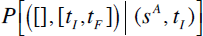 becomes:

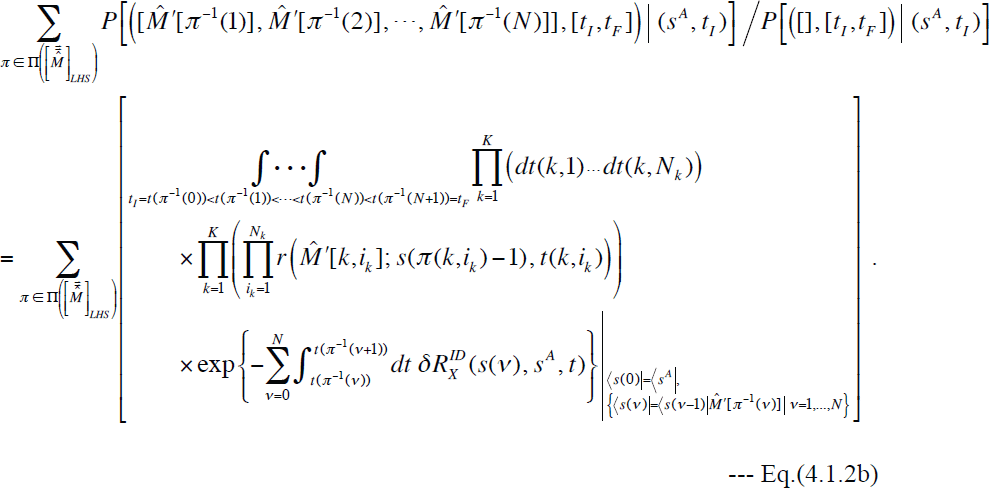

Comparing Eq.(4.1.2a) and Eq.(4.1.2b), we can see that Eq.(4.1.1) should hold if and *nearly* only if the following two equations are satisfied. (a) One is an equation between the integrands, *i.e.*,

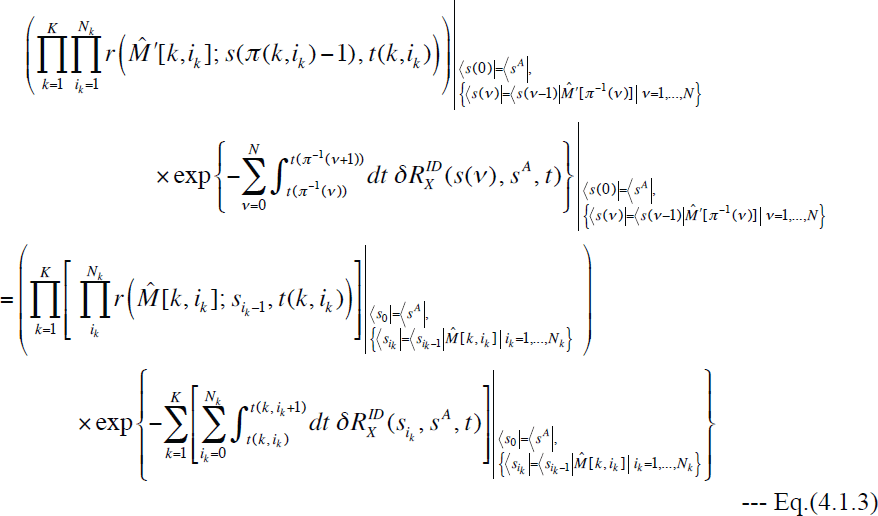

for each map 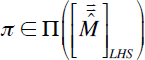 and its associated temporal order of events. And (b) the other is an equation between the multiple-time integration operations, *i.e.*,

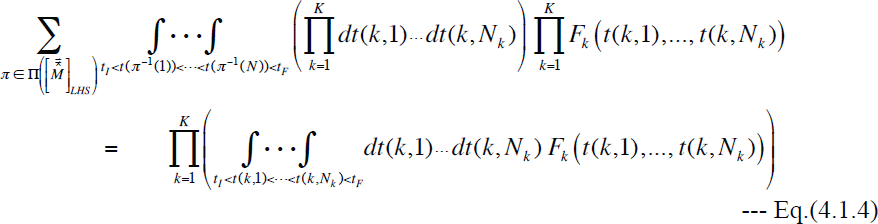

for any set of non-singular functions, 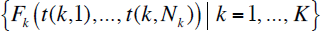. The first core equation, Eq.(4.1.3), holds only under an appropriate set of conditions on the indel rate parameters. The second core equation, Eq.(4.1.4), is an identity, which is intuitively plausible but whose rigorous proof is not so straightforward. Its rigorous proof is given in Appendix A4.

The both sides of Eq.(4.1.3) exhibit very similar forms. Each of them is a product of the rates of indels that actually occurred or their equivalents, multiplied by an exponential. And the exponent is the summation of time-integrated increments, of the exit rates of the states that the sequence actually (or virtually) went through, compared to the exit rate of the ancestral state. Thus, aside from miraculous, exceptional cases, it would be natural to expect the equations to be satisfied for each of the factors. This reasoning gives two types of equations. One is a set of equations for the factors in the product,

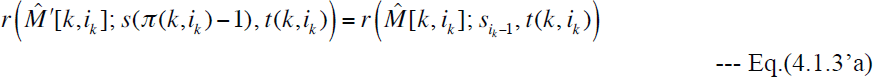

for 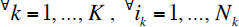, and 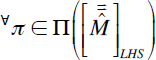. And the other is an equation for the exponent,

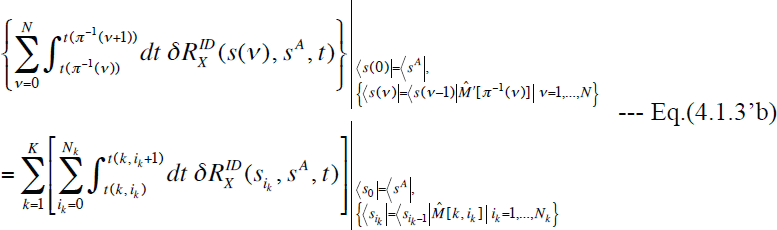

for 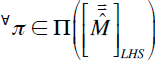. Here, we set 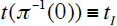 and 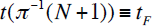. In Eq.(4.1.3’a), 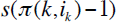 is the sequence state immediately before 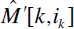 in the global indel history, and 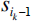 is the state immediately before 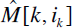 in the isolated *k* th local indel history. The only difference between both sides of Eq.(4.1.3’a) is in the states. In general, 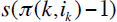 on the left hand side resulted from some of the events in the other local indel histories, on top of 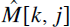 with 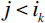. In contrast, 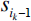 on the right hand side will never be impacted by the other local histories. Thus, Eq.(4.1.3’a) simply states, for the PWA probability to be factorized, “the rate parameter for each indel operator in each local indel history must never be influenced by the actions of any indels that occurred before *t*(*k*,*i_k_*) and that belong to any other local histories.” Meanwhile, Eq.(4.1.3’b) appear more formidable than Eq.(4.1.3’). Nevertheless, we can prove the following proposition.

**[Proposition 4.1.1]**

“Let 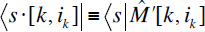 and 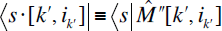 (with 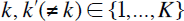 be the states resulting from the actions of the equivalents of events 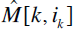 and 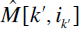, respectively, on 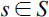 And let 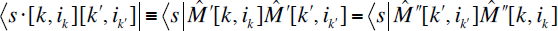 be the states resulting from the consecutive actions of the equivalents of 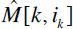 and 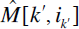, on *s*. The equation for the exponents, Eq.(4.1.3’b), holds for every global history 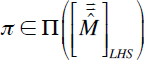 and for each of its sub-histories that could occur in any sub-interval, [*t_I_*, *t*] with 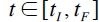, if and only if the equation,

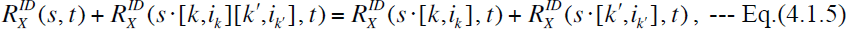

holds for every pair, 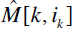 and 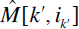 (with 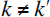), in the LHS 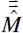, for every possible state 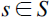 before both equivalents of 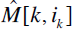 and of 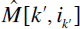 in the global histories in 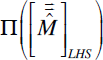, and at any time 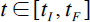.”

The detailed proof of this proposition is given in Appendix A5. In the proposition, the applicable scope of Eq.(4.1.3’b) was extended to all sub-histories of global histories belonging to 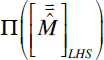 and to any sub-interval, [*t_I_*, *t*], of [*t_I_*, *t_F_*]. This extension would be acceptable in practical analyses, where what we actually want is to factorize *all* alignment probabilities during *any* time interval. We can clarify the meaning of Eq.(IV-1.5) by rewriting it as follows:

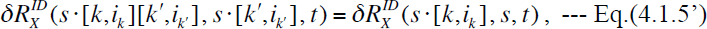

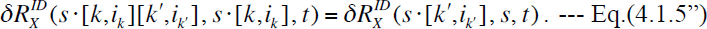

These equations mean that the increment of the exit rate due to an event in a local indel history must be independent of the effect of any event in any other local indel history.

To summarize, we derived a sufficient and nearly necessary set of conditions, Eq.(4.1.3’a) and Eq.(4.1.5), under which the integrand of the probability of an indel history can be factorized, as in Eq.(4.1.3). To clarify what these conditions mean, we here rephrase them in words. Eq.(4.1.3’a) can be rephrased as follows.

**Condition (i):** “The rate parameter, 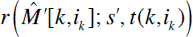, for each actually occurred indel event 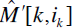 will not be influenced by the action of any indel events outside of the *k* th local history before 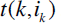.”

Second, we can rephrase Eq.(4.1.5) as follows.

**Condition (ii):** “Let 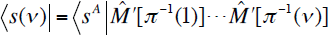 be the state resulting from the actions of events up to (and including) the *ν* th event in a global history corresponding to a map 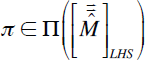, and let 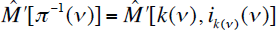 be the *ν* th event in the global history. Then, the increment of the exit rate, 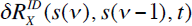, due to the event 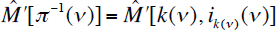, will not be influenced by the actions of any indel events outside of the *k*(*ν*) th local history before 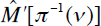.”

If this set of conditions is satisfied for all global indel histories in a LHS equivalence class 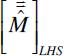, then, Eq.(4.1.3) holds for all integrands. This, combined with the identity on the domains of integration, Eq.(4.1.4), make the total probability of 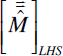 factorable, as in Eqs.(4.1.1a,b). (Someone might guess that the condition (ii) should follow from the condition (i) almost trivially. We will see that this guess is wrong in Section 5.)

Now, in terms of the probabilities of the LHS equivalence classes of global indel histories, we re-express Eq.(4.1.13) for the probability of a PWA as:

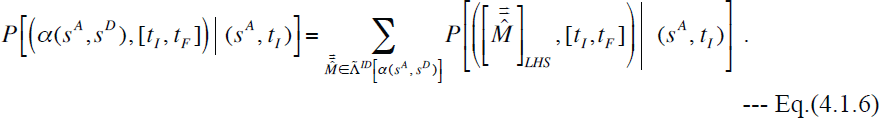

And we suppose that each term in the summation on the right was factorized as in Eqs.(4.1.1a,b). It should be noted that the number of local indel histories, *K*, could vary depending on the, 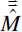. Here, we introduce the notation 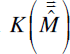, to remind this dependence of the number of local histories on the LHS. A local indel history could occur either between two or between a PAS and a sequence end. Thus, in principle, the largest possible set of regions that could potentially accommodate local indel histories consists of the region between the left-end of the PWA and the leftmost PAS, the regions each of which is between a PAS and the next PAS, and the region between the rightmost PAS and the right-end of the PWA. However, some of these regions may not be able to accommodate any local history because they do not have adequate nonzero indel rates. Or, local indel histories in some adjacent (but disconnected) regions may not be factorable from each other because either of the conditions (i) and (ii) is violated between them. In this case, the regions will be put together to form a single region to define local indel histories. Let 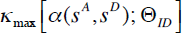, or 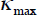 for short, be the number of regions in 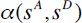 that can possibly accommodate local indel histories, given an indel model including the rate parameters, 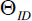. And let 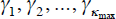 be such potentially local-history-accommodating regions in 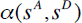, positioned from left to right along the PWA. First, we obviously have 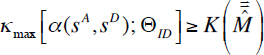 for 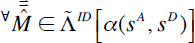, because each LHS defined under this PWA partitioning fills each region 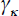 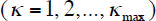 with at most one local history. Second, let 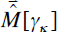 denote such a local history to fill 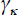. Then, we can represent any 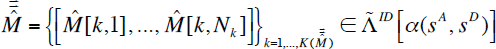 as a vector with 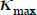 components: 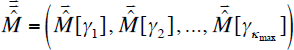. Here 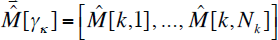 if the *k* th local history is confined in 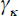, or 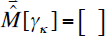 (empty) if no events in the LHS occurred in *γ*_κ_ (Figure 10). Using these notations, the factorization, Eq.(4.1.1a), of the probability of an LHS equivalence class 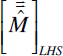 is re-expressed as:

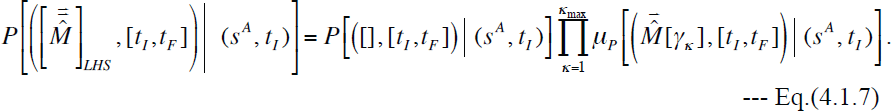

**Figure 10.**
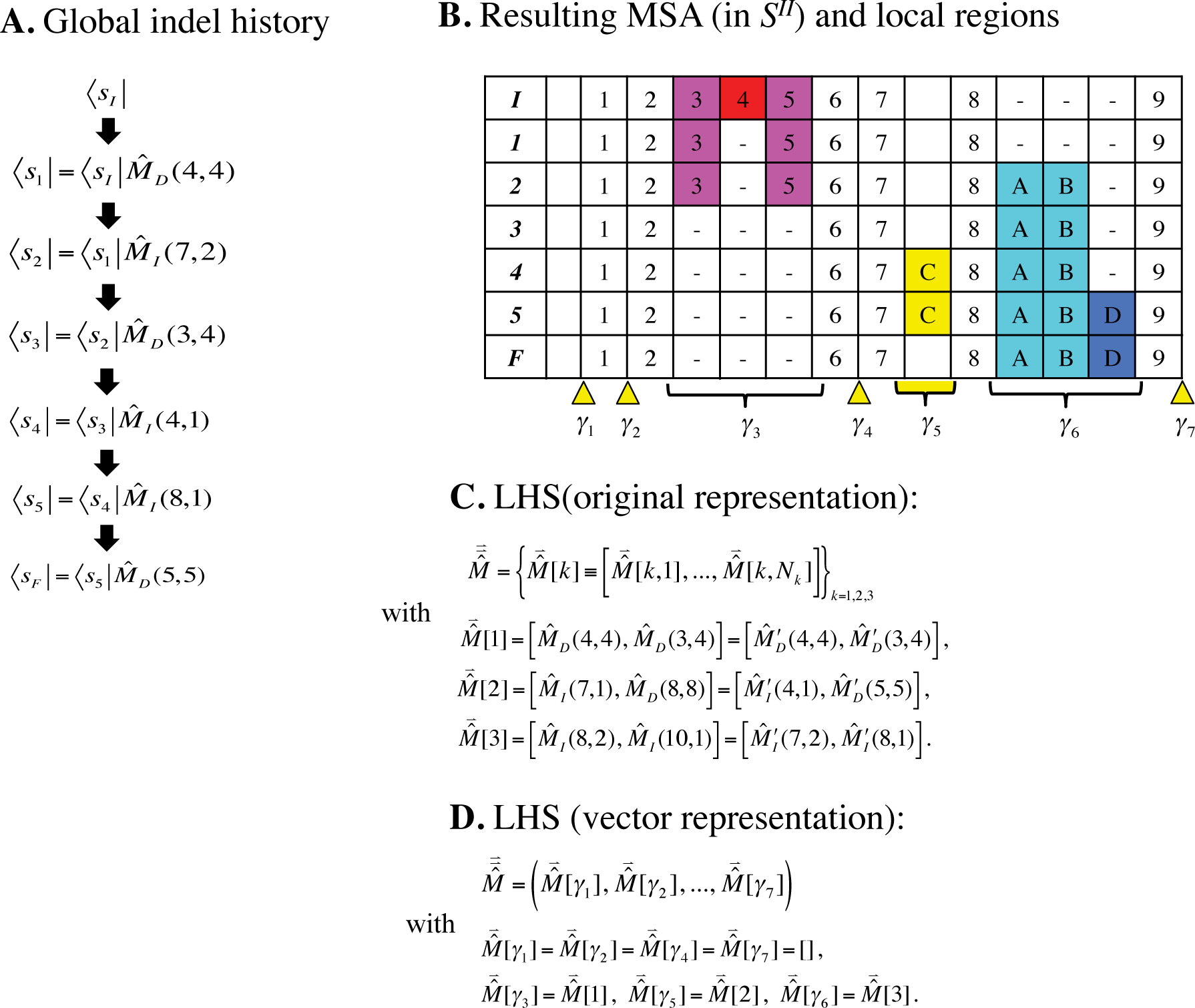
“Vector” representation of local history set (LHS) along time interval. **A.** An example global indel history, consisting of six indel events and seven resulting sequence states (including the initial state *S_I_*). **B.** The resulting MSA among the sequence states that the indel history went through. The boldface letters in the leftmost column indicates the sequence states in the global history (panel A). The 1-9,A-D in the cells are the ancestry identifiers of the sites (in the state space *S^II^*). The magenta and red cells represent the sites to be deleted. The cyan and blue cells represent the inserted sites. The yellow cells represent the inserted sites that are to be deleted. Below the MSA, the underbraces indicate the regions 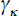 (κ = 3, 5, 6 in this example) that actually accommodate local indel histories. And the yellow wedges indicate the regions 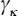 (κ = 1, 2, 4, 7 in this example) that can potentially accommodate local indel histories, but that actually do not. In this example, we have 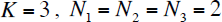 and 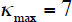. **C.** The original representation of the LHS. In each defining equation for 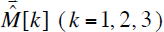, the expression in the middle is the local history represented by its action on the initial state *S_I_*. And on the rightmost side is the representation by the actual indel events in the global history (in panel A). The prime there indicates that each defining event is equivalent to, but not necessarily equal to, the corresponding event in the global history. **D.** The vector representation of the LHS. The ”[]” denotes an empty local history, in which no indel event took place.

Here 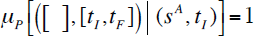 should be kept in mind. Now, consider the space 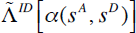 itself. Any two different LHSs in this space differ at least by a local history in some *γ_κ_*. Conversely, any given set of 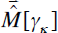’s in all 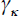’s, each of which is consistent with the PWA restricted in the region, defines a LHS in 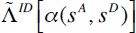. Thus, the set 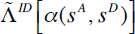 should be represented as a “direct product”: 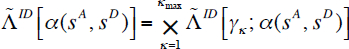, where 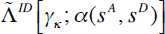 denotes the set of local indel histories in 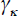 that are consistent with the sub-PWA of 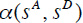 confined in 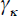. Using this structure of 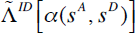 and substituting Eq. (IV-1.7) for each 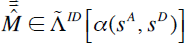 into Eq.(4.1.6), we finally get the desired factorization of the PWA probability:

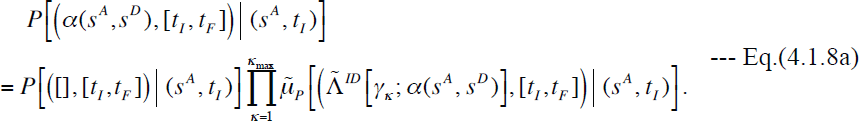

Here the multiplication factor,

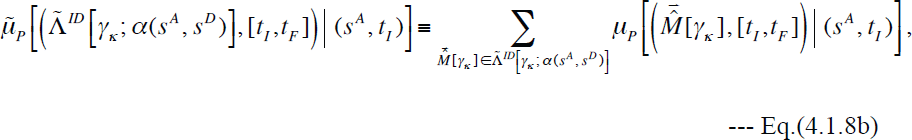

represents the total contribution to the PWA probability by *all* consistent local indel histories that can take place in 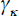.

#### 4.2. Factorization of probability of given MSA

We can use the results for the PWA probability in the last subsection to factorize 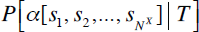, the probability of a given MSA (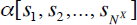) under a given phylogenetic tree (*T*). We could do this in two different ways, one directly starting from Eq.(3.2.11) accompanied by Eq.(3.2.3) and the other starting from Eqs.(3.2.13a,b’). It should be noted first that 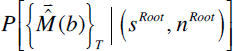, the probability of a given indel history along a tree 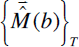 given by Eq.(3.2.3), is not factorable by itself, for a reason similar to that in the pairwise case. Thus, as in the pairwise case, let us consider the total probability of a LHS equivalence class *along T*, 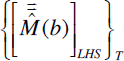, defined in Eq. (3.4.1):

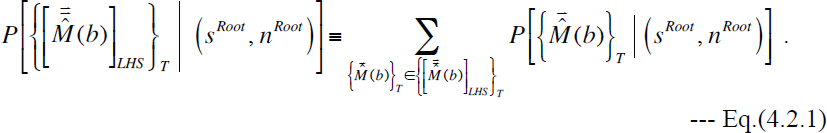

Using Eq.(3.2.3) that defines 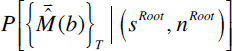 and Eq.(3.4.1) that defines 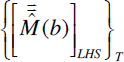, we have:

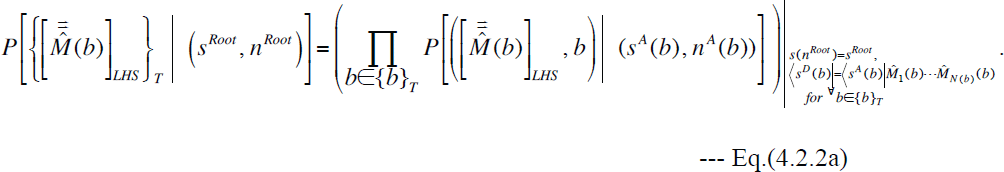

Here,

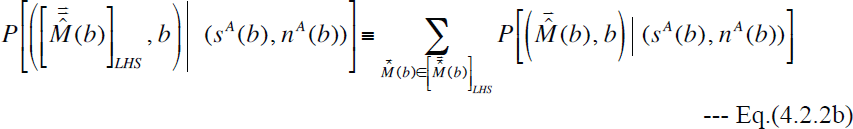

is an equivalent of Eq.(4.1.1a) along the branch *b*. Thus, under the same set of conditions, (i) and (ii), on the rate parameters and the exit rates, Eq.(4.2.2b) for each branch is factorable as in Eq.(4.1.7), giving:

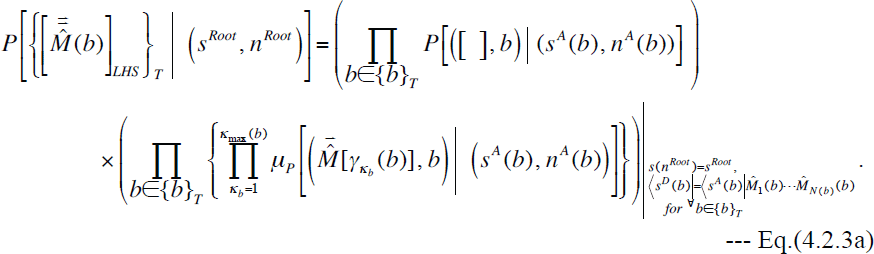

Here 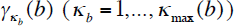 is a region that potentially accommodates a local indel history along branch *b*, and we made the replacements of the arguments for 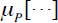 similar to those in Eq.(3.2.2). The first term on the right-hand side is actually an exponential:

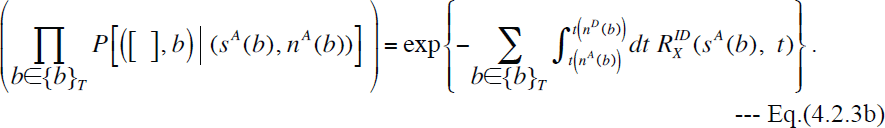

**Figure 11.**
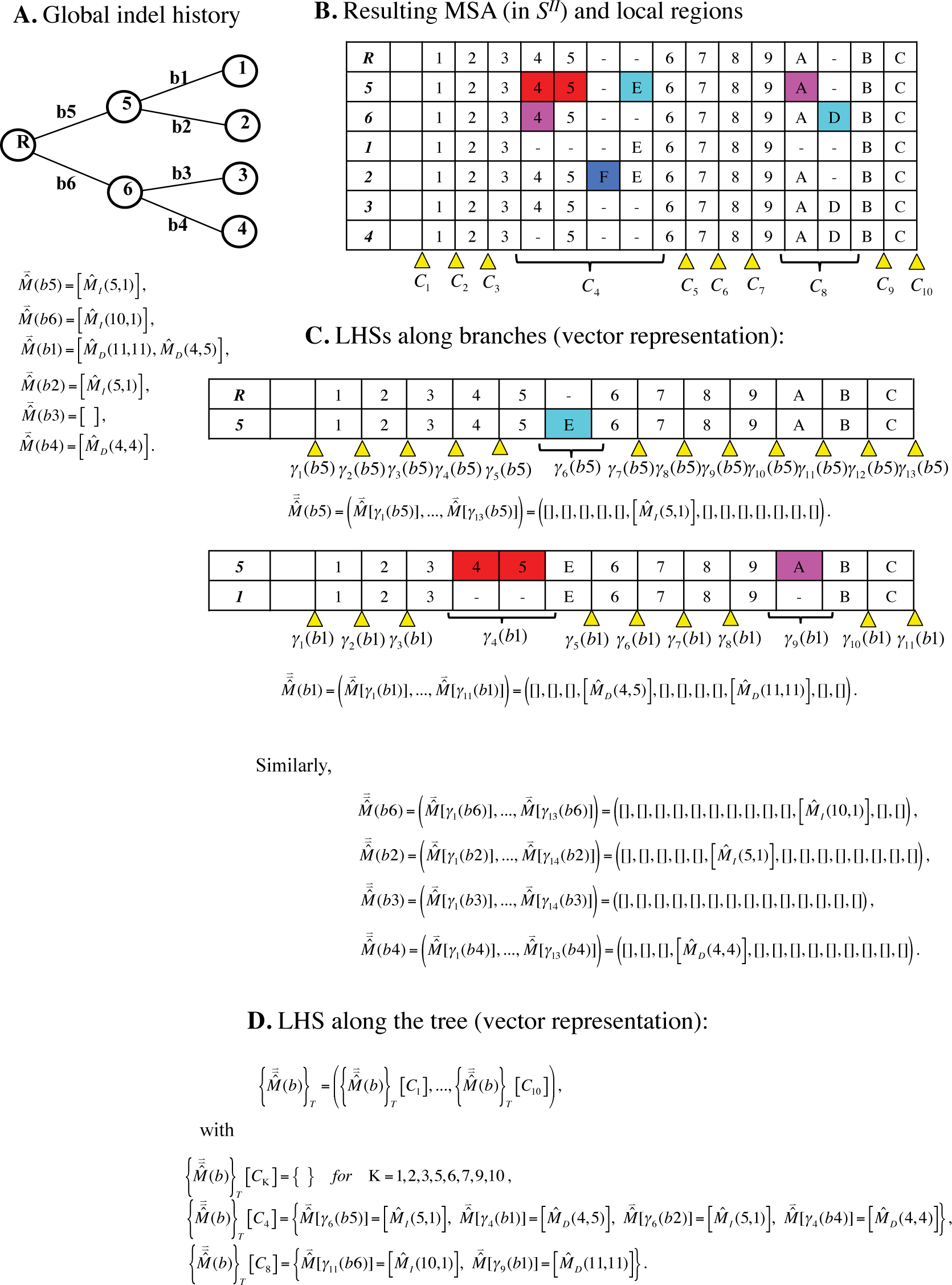
MSA regions potentially able to accommodate local indel histories along tree. **A.** A global indel history along a tree. Sequence IDs are assigned to the nodes. Each branch is accompanied by an ID (*b*1-*b*6) and a global indel history along it. The “R” stands for the root. **B.** Resulting MSA of the “extant" sequences at external nodes and the ancestral sequences at internal nodes. The boldface letters in the leftmost column are the node IDs. Below the MSA, the underbraces indicate the regions *C*_K_ (**K** = 4, 8 in this example) that actually accommodate local indel histories along the tree. And the yellow wedges indicate the regions *C*_K_ (Κ = 1, 2, 3, 5, 6, 7, 9, 10 in this example) that can potentially accommodate local indel histories along the tree, but that actually do not. In this example, we have **K**_max_ = 10. **C.** LHSs along the branches (in the vector representation). As examples, the PWAs along branches *b*_1_ and *b*_5_ are also shown, along with their own regions that can potentially accommodate local histories. **D.** The LHS along the tree (vector representation). Only the non-empty components were shown explicitly. This figure follows basically the same notation as Figure 10 does. A cell in the MSA is colored only if it is inserted/deleted along an adjacent branch.

To go further, we partition the MSA 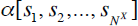 into regions between a gapless column and the next gapless column and regions on the left and right, respectively, of the left- and right-most gapless columns. But we lump together the regions into a single region if they fail to mutually satisfy either condition (i) or (ii) in Subsection 4.1. Let 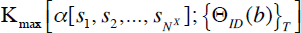, or 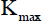 for short, be the number of all such potential host regions in 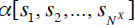 under a given set of rate parameters, 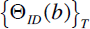. We always have 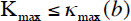 for 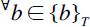. And let 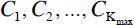 denote such regions. Each region, 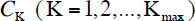, can potentially accommodate a local indel history along *T*, denoted as 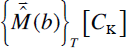, which is actually composed of local indel histories along all branches and confined in *C*_K_ (Figure 11). Thus, Eqs.(4.2.3a,b) can be rearranged as:

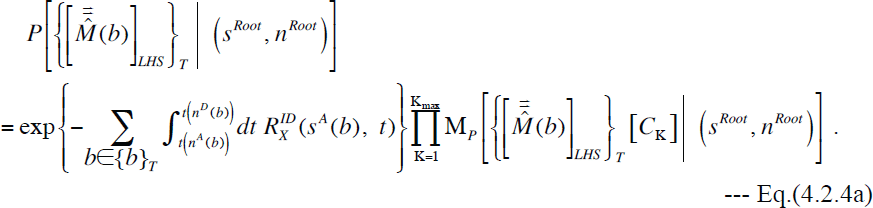

Here the multiplication factor,

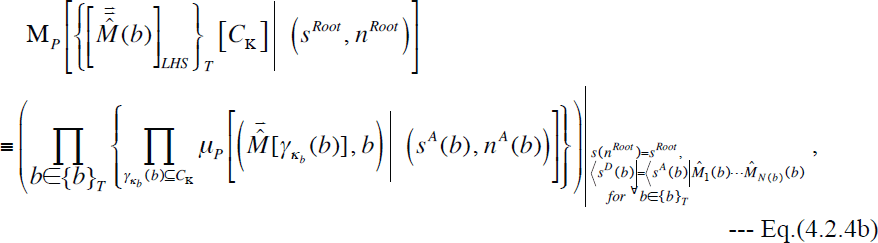

represents the total contribution from a LHS equivalence class, 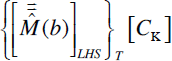, of local indel histories along *T* that are confined in *C*_K_. When factorizing the probability of a MSA, 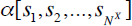, Eq.(4.2.4a) is not the final form of the factorization of 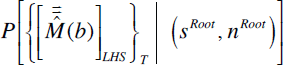, because the exit rates 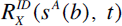 could vary depending on the LHS equivalence class 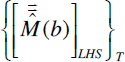. To finalize its factorization, we introduce a “reference” root sequence state, 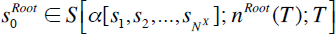. One good candidate for 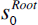 would be a root state obtained by applying the Dollo parsimony principle (Farris 1977) to each column of the MSA, because it is arguably the most readily available state that satisfies the phylogenetic correctness condition along the entire MSA. Given a reference, 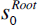, each ancestral state 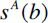 should differ from 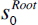 only within some *C*_K_’s. Moreover, the condition (ii) suggests that the impacts of their differences within separate *C*_K_’s on the exit rate should be independent of each other. Thus, we have:

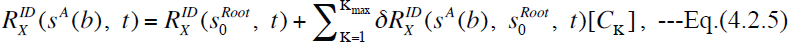

where 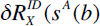, 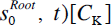 is the increment of the exit rate due to the difference between *s^A^*(*b*) and 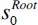 within the region *C*_K_. Especially, we have 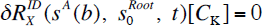 unless the states differ within *C*_K_. Substituting Eq.(4.2.5) into Eq.(4.2.4a),we get the desired factorization:

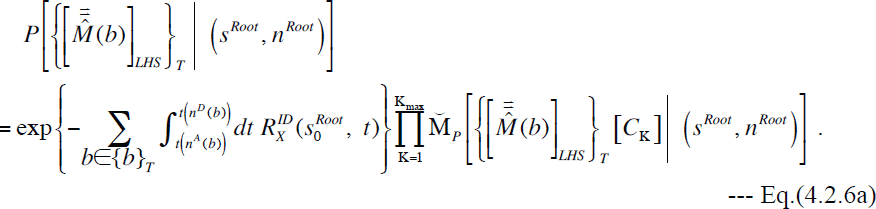

Here, we defined an augmented multiplication factor,

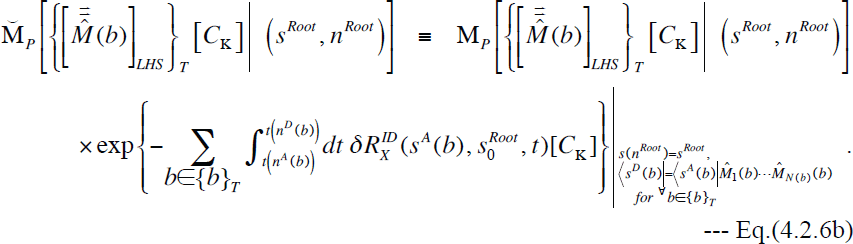

Now, using the decomposition, Eq.(3.4.2), of 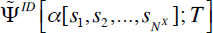, *i.e.*, the set of all pairs, each of an indel history and a root state, consistent with the MSA, Eq.(3.2.11) can be rewritten as:

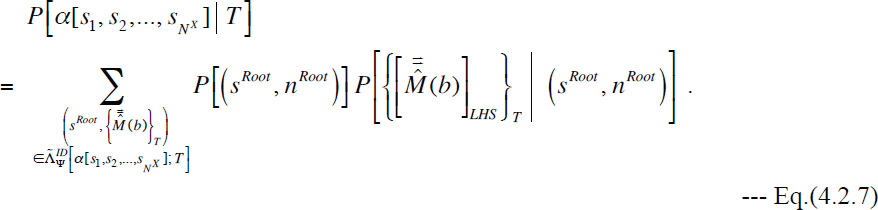

To go further, we here first assume that the following equation holds for the probability of the root state:

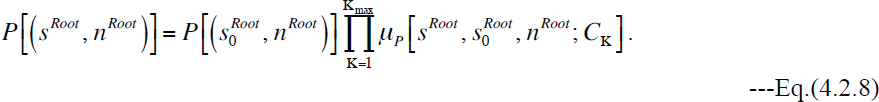

Here the multiplication factor 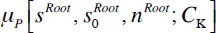 represents the change in the state probability at the root due to the difference between 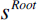 and 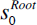 within *C*_K_. Eq.(4.2.8) holds, *e.g.*, when 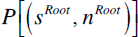 is a geometric distribution or a uniform distribution of the root sequence length, 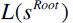. Geometric distributions of sequence lengths were commonly used by HMMs and by transducers. The uniform distribution may be a good approximation if we can assume that the ancestral sequence was sampled randomly from a chromosome of length *L_C_*. In this case, the distribution of the sequence length 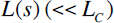 would be proportional to 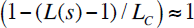. Second, similarly to 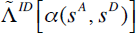 discussed above Eq.(IV-1.8a), we also express 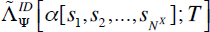 as a “direct product”: 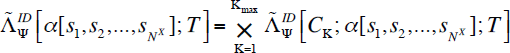, where 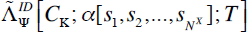 is the set of all local indel histories along *T* (each accompanying a root state) and within *C*_K_ that are consistent with the sub-MSA of 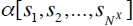 restricted to *C*_K_. Then, substituting Eq.(4.2.6a) and Eq.(4.2.8) into Eq.(4.2.7), and using the direct-product structure, we can finally factorize the probability of 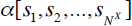:

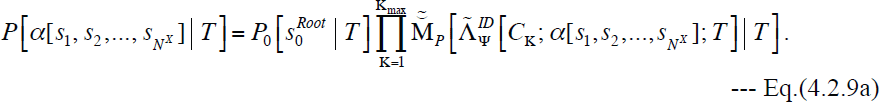

Here,

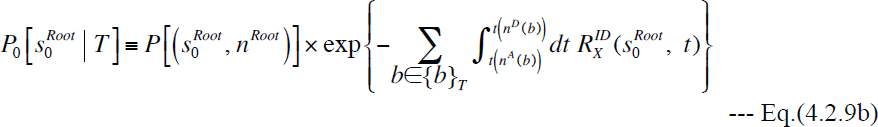

is the probability that the reference root state 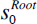 was present at the root and underwent no indel events throughout the evolutionary history along the tree. The augmented multiplication factor,

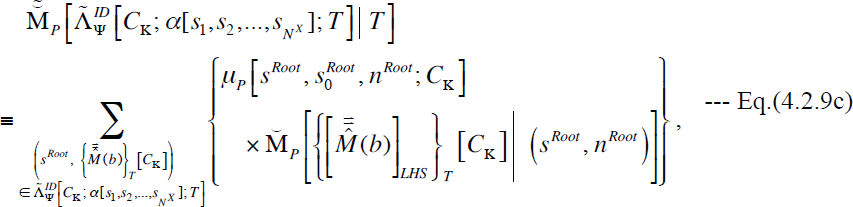

provides the total probability change due to the MSA-consistent local histories along the tree and confined in *C*_K_.

Now, we briefly explain how we can achieve the MSA probability factorization along the other route starting from Eqs.(3.2.13a,b’). Each term in Eq.(3.2.13a) is the probability of MSA-consistent indel histories with a fixed set, 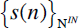, of sequence states at internal nodes, and Eq.(3.2.13b’) expresses the term as a product of the probabilities of PWAs, each between the fixed ancestral and descendant states along a branch. Such probabilities of PWAs can be factorized using Eq.(4.1.8a), and we could lump together the multiplication factors within the same region, e.g., *C*_K_, but along different branches, into a single factor representing the total probability change contributed from *C*_K_. The product of 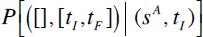’s can be re-expressed as an exponential just as in Eq.(4.2.3b), and then processed just like from Eq.(4.2.4a) to Eq.(4.2.6a), using Eq.(4.2.5). Then, we use Eq.(4.2.8) to factorize the root state probability in Eq.(3.2.13b’). Then, we introduce the direct product structure of the set of MSA-consistent internal node states:

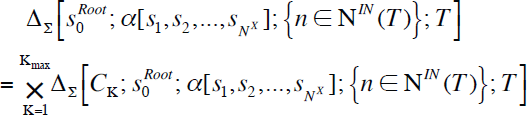

Here 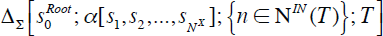 is the space of *deviations* of MSA-consistent internal sequence states from the reference state 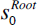, and 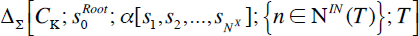 is the space of such deviations *within* the region *C*_K_. Using this direct product structure, the MAS probability can be factorized similarly to Eqs.(4.2.9a,b,c) but with the multiplication factor from *C*_K_ organized differently from that in Eq.(4.2.9c). The difference arises because the first route treats the (local) indel histories as fundamental building blocks, whereas the second route focuses on the (local) sequence states at the internal nodes.

### 5. Indel models with factorable alignment probabilities

In the previous section, we derived a sufficient and nearly necessary set of conditions for the factorability of PWA probabilities. The conditions are briefly stated as follows.

**Condition (i):** “The rate parameter, 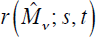, for each indel event (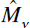) in every PWA-consistent global history will not be influenced by the actions of any indel events that occurred before 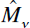 and outside of the local history to which 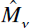 belongs.”

**Condition (ii):** “The increment of the exit rate, 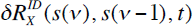, due to the event 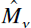 (with 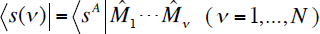 for a global history 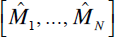), will not be influenced by the action of any indel events that occurred before 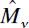 and outside of the local history to which 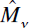 belongs.”

In this section, we will actually see some example indel models that indeed satisfy, or do not satisfy, these conditions. Before going into specific examples, we here note that models satisfying the above conditions are distinct from an indel model described by the context-independent rate grammar proposed by Miklós et al. (2004), although they are somewhat similar to each other. First, as already explained in Subsection 2.1, our state space could be more general than that for (the indel component of) the rate grammar. Second, the condition (i) could be more liberal than the context-independence condition, in the sense that the former could allow the rates to depend on the state of a *close vicinity of* the insertion position or the deleted subsequence. And third, as we will see in Subsection 5.2, the condition (ii) may not necessarily be satisfied even if the context-independence is satisfied by the rates of all indel events in the alignment-consistent histories.

#### 5.1. Space-homogeneous models

The simplest conceivable indel models would impose that the indel rate parameters be space-homogeneous, *i.e.*, independent of the positions where the indels occur:

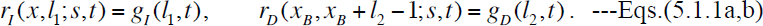

In fully space-homogeneous models, these equations hold for 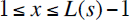, 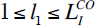, 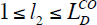, and 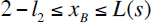. (Depending on the model, 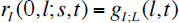 and 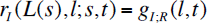 could differ from 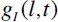 in Eq.(5.1.1a).) In fact, these conditions were imposed by nearly all continuous-time Markov models of indels that were studied in the past (except, *e.g.*, the TKF92 model (Thorne et al. 1992)). Note that the rate parameters in Eqs.(5.1.1a,b) could depend on time, although most models used thus far imposed that the rates be time-independent as well. Eq.(5.1.1a,b) automatically guarantees the condition (i). Thus, all we have to do is to check whether or not the condition (ii) is also satisfied. Indeed, we can show it is. The exit rate from Eq.(5.1.1a,b) is calculated exactly in the same way how we derived Eq.(2.4.7e), and we find that it is an affine function of the sequence length (*L*):

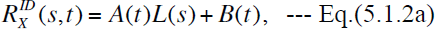

with 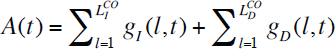 and 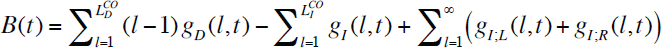. If the exit rate is affine, we have:

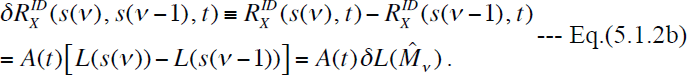

Here 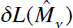 is the length change caused by the event 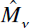. The rightmost hand side of this equation depends only on 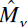 and the time it occurred, but not on the other events in the indel history. Thus, the condition (ii) is always satisfied under fully space-homogenous models, which means that alignment probabilities calculated *ab initio* (as in Section 3) under such models are factorable, as shown in Section 4.

An important special case of the space-homogeneous model, Eqs.(5.1.1a,b), is the indel model used by Dawg (Cartwright 2005), whose indel rate parameters were already given in Eqs.(2.4.4a,b). This is a special case of Eqs.(5.1.1a,b) with time-independent indel rates, and thus provides factorable alignment probabilities. This model is probably among the most flexible ones used thus far. The model accommodates any distributions of indel lengths, and allows independent length distributions, and independent total rates, for insertions and deletions. In parts II and III (Ezawa, Graur and Landan 2015a,b), we will base our calculations mostly on this model.

Another important special case is the “long indel” model (Miklós et al. 2004), whose rate parameters are given by Eqs.(2.4.5a-e), which are also time-independent. This model is less flexible than Dawg’s model, because its indel rates are subject to the detailed-balance conditions, Eq.(2.4.6a-d). Like Dawg’s model, this model is a special case of the model defined by Eqs.(5.1.1a,b). Thus, the alignment probabilities calculated under it are indeed factorable, as Miklós et al. (2004) conjectured. Indeed, we can show that, as far as each LHS equivalence class is concerned, the indel component of its probability calculated according to the recipe of Miklós et al. (2004) equals Eq.(4.1.7), *i.e.*, the probability of the LHS equivalence class via our *ab initio* formulation, calculated with the indel rate parameters Eq.(2.4.5a-e). The proof is given in Appendix A6. This equivalence is likely because the contribution from each local indel history, e.g., the expression in square brackets in Eq.(4.1.2a), is calculated essentially from the increments of the exit rate for the entire sequence, as well as from the rates of indels in the local history. In the special case with space-homogeneous indel rates, the exit-rate increment for an entire sequence coincides with the exit-rate increment calculated for the “chop-zone” according to Miklós et al.’s recipe. Thus, under the long-indel model, Eqs.(2.4.5a-e), (or actually under any space-homogeneous model,) the indel component of the probability of a PWA calculated according to the prescription of Miklós et al. (2004) should also be equivalent to that calculated via our *ab initio* formulation given in Subsection 4.1, *as long as the contributing local indel histories are correctly enumerated*. Actually, it is not so straightforward to enumerate *all*(local) indel histories consistent with a (local) PWA, as we will see in Subsection 1.2 in part II (Ezawa, Graur and Landan 2015a), due to the complexities on the equivalent local histories explained in Subsection 3.3.

Regarding the insertion rates, we could relax the condition of space-homogeneity without compromising the factorability of alignment probabilities. For example, we could consider the rates of insertions between the *x* th and *x* + 1 th sites along the sequence *s* as a function of the ancestries of these sites, *υ*(*s*,*x*)and *υ*;(*s*, *x* + 1):

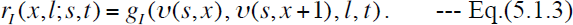

Of course, these rates satisfy the condition (i). And Eq.(5.1.3) and the space-homogeneous deletion rates, Eq.(5.1.1b), still gives an exit rate whose increment due to an indel event depends only on the inserted/deleted sub-sequence (and flanking sites) but not on the regions separated from it by at least a PAS. This means that the model also satisfies the condition (ii). Thus, the alignment probabilities should be factorable also under this model with somewhat generalized insertion rates. Relaxing the space-homogeneity of deletion rates, however, is somewhat difficult, particularly because of the condition (ii). In the following subsections, we will attempt to do it.

#### 5.2. Indel models containing biologically essential regions

The space-homogeneous models discussed above, including Dawg’s model and the long-indel model, may decently approximate the neutral evolution of a sequence region under no selective pressure. A real genome, however, is scattered with regions and sites under strong or weak purifying selection.

First, we consider a simplest model that implement such a situation, where a neutrally evolving region is left-flanked by a region (or a site) that is biologically essential. This situation could be implemented with the rate parameters given by Eq.(5.1.1) for the same domains as in the fully space-homogeneous models, except that the domain for *x_B_* changed to 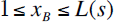. In other words, we have

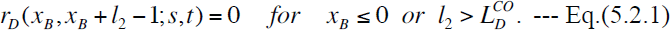

In this case, the exit rate is given by an affine form, Eq.(5.1.2a), with *A*(*t*) exactly the same as for the fully space-homogeneous case, and with 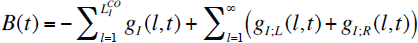. Because the exit rate is affine, this model satisfies the condition (ii). The condition (i) is also satisfied, because the rate parameter of an event will remain unaffected by the events outside of the local history it belongs to. Thus, the alignment probabilities under this model should also be factorable. By symmetry, we expect that alignment probabilities should also be factorable under a model where a neutrally evolving region is right-flanked by a region (or a site) that is biologically essential. This situation can be implemented by the insertion rates, Eq.(5.1.1a) for exactly the same domain as before, and the deletion rates:

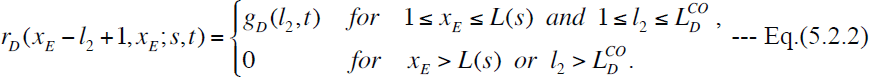

The exit rate is exactly the same as in the left-flanking case, and thus the alignment probabilities are indeed factorable under this model as well.

Second, we consider a model where a neutrally evolving region is flanked by biologically essential regions (or sites) from both sides. The insertion rates of this model are given by Eq.(5.1.1a) with the same domain, and the deletion rates are:

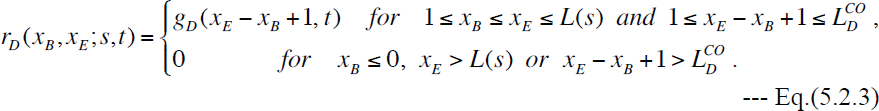

The exit rate for this model is calculated as:

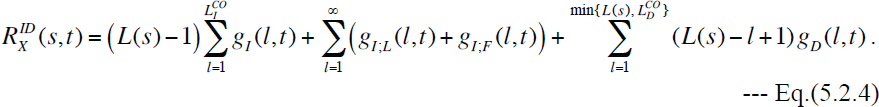

For 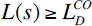, this is affine, and given by Eq.(5.1.2a), with exactly the same *A*(*t*) as before and 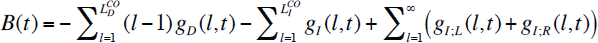.

Therefore, if the sequence length remains greater than or equal to 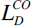 throughout all indel histories that could give rise to the alignment in question, the alignment probability is still factorable even under this model. For 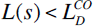, in contrast, it exhibits a *non-affine* form:

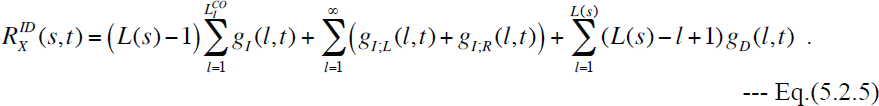

Thus,in this case, the condition (ii) will not be satisfied in general. As an example, let us consider a sequence state 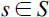 with 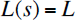, and the action of two separated deletions, 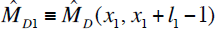 and 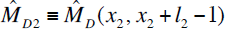 with 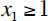 and 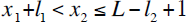. And we use the notations, 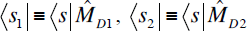, and 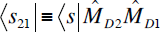. Then, substituting 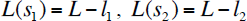, and 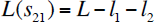 into Eq.(5.2.5), we have:

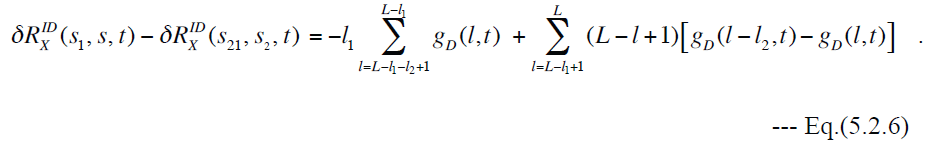

(The derivation is in Appendix A7.) Although the terms on the right-hand side of Eq.(5.2.6) might exactly cancel out for a special form of *g_D_*(*l,t*) (and for special values of *L*, *l*_1_, and *l*_2_), they will not in general. Thus, although the indel events in other local histories do not impact the rate of the indel in question,they do impact the increment of the exit rate the event causes. Therefore, in this case, the alignment probabilities are *not* factorable. Because the cut-off lengths, 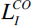 and 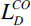, were originally introduced as a proxy of the collective effect of physical and biological constraints on the indel size, it would accord with the common sense to assume them to be longer than a neutrally evolving region, *i.e.*, 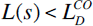. Then, the above argument implies that the alignment probabilities are unlikely to be *exactly* factorable. Nevertheless, they may be *approximately* factorable, if the “difference between differences” as in Eq.(5.2.6) is much less than 1. This could happen, *e.g.*, when 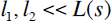 so that 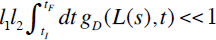 for all pairs of indels belonging to different local histories.

Third, we consider a model where a sequence contains one or more conserved regions. In this case, we need to work with the state space *S^II^* or *S^III^*, because it is essential to keep track of ancestral residues, which the simple structure of *S^I^* cannot do. It should be understood that the argument unfolded below is implicitly mediated by the ancestries naturally assigned to the sites by the state space *S^II^* or *S^III^*, although we will introduce a more convenient notation to keep track of ancestral sites and to figure out the positioning of inserted sites relative to them. Let 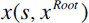 be the site number (*i.e.*, the coordinate),in a sequence *s*, of the site whose site number was *x^Root^* in the ancestral sequence *s^Root^*. And assume that the sequence has *Y* (≥1) conserved region(s) defined by the closed interval(s), 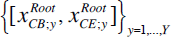, in *s^Root^*. (Assume 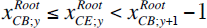 for 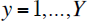, with 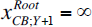.) In this situation, the indel rates are constrained as:

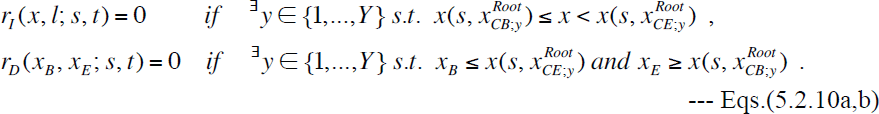

In other words, the indel rate could be nonzero only if the insertion position or the deleted subsequence does not overlap any conserved region The exit rate is then decomposed as:

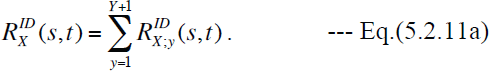

Here

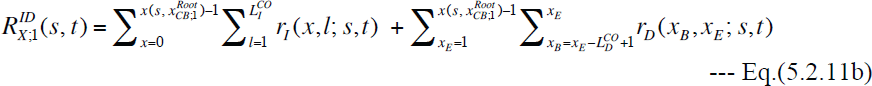

is the exit rate for the region on the left of the leftmost conserved region.

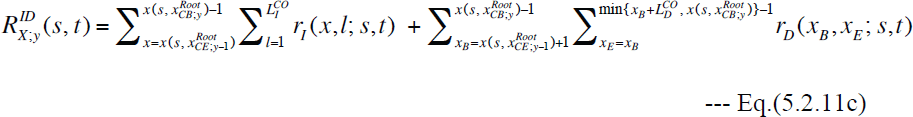

is the exit rate for the region between the *y*−1 th and *y* th conserved regions (*y* = 2, …, *Y*). And

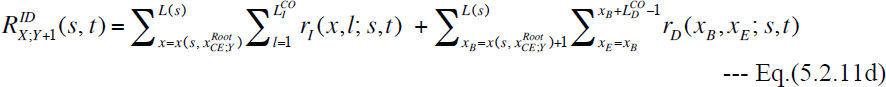

>is the exit rate for the region on the right of the rightmost conserved region. Thus, if the indel rates in each evolvable region do not depend on the portion of the sequence states inany other evolvable region, the different evolvable regions are completely decoupled regarding evolution via indels. Thus, in this case, an alignment probability can be factorized into the product of contributions from these evolvable regions. (Actually, we could factorize the stochastic evolution operator itself into a tensor product form, if desired.) The contributions from the both ends of the sequence will be further factorable if we assume homogeneous indel rates in these regions, as in the first case (Eq.(5.2.1) or Eq.(5.2.2)), where a neutral sequence is flanked from only one side by a conserved region. The region between two neighboring conserved regions, however, is essentially the same as the second case (Eq.(5.2.3)), whose alignment probabilities are not exactly factorable in general even if the indel rates are space-homogeneous. Thus, in this situation, it may not be so meaningful to further restrict the functional forms of indel rates in each evolvable region. This means that, if desired, we could freely fit the rate parameters to approximate the real position-dependent indel rates in the region.

#### 5.3. More general model

The models considered thus far contained only sites of somewhat extreme biological importance: either essential (*i.e.*, completely conserved) or unimportant (*i.e.*, neutrally evolving). However, the sites of a real sequence should have a wide variety of biological importance, and evolve under different levels of selective pressures. Besides, different regions may also have different mutation rates, depending on the sequence or epigenetic contexts. Thus, it would be preferable if we can allow indel rates to vary more flexibly across regions while keeping alignment probabilities somewhat factorable. A first clue comes from the fact that, if two different sets of indel rates satisfy the conditions (i) and (ii) for a given LHS, a linear combination of the two sets also satisfies the conditions. Another important clue is that the set of indel rates in the last example in Subsection 5.2 could be considered as composed of different sets of indel rates. Each of them is confined in an evolvable region, 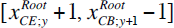 (*y* = 0, 1, …, *Y*), and depends only on the portion of the sequence state within the region. (Here we considered 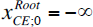 and 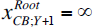.) Inspired by these two clues, we first define a set of non-overlapping regions, 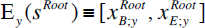, that existed in (or beyond the boundaries of) the root sequence 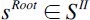 or *S^III^*. We define the “descendant,” E*_y_* (*s*), of 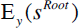 in a descendant state (*s*) by the closed interval, 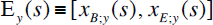, where 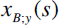 and 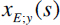 are the leftmost and the rightmost sites, respectively, among those descended from the sites in 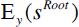. Then, based on them, we define an indel model whose rate parameters are given by:

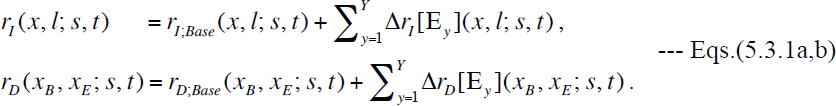

Here, the “baseline” indel rates, 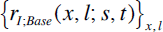, and 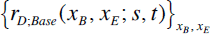, are given by the flanking-site-dependent insertion rates Eq.(5.1.3) and the space-homogeneous deletion rates Eq.(5.1.1b), as in the bottom of Subsection 5.1. The region-specific increments of the indel rates, 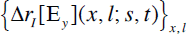 and 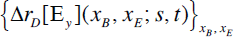, can be non-zero *only within* the region, 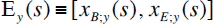, defined above (panel A of Figure 12). Moreover, the increments can depend only on the portion of the sequence state within E*_y_*(*S*). The increments can be negative, as long as the entire rates, Eqs.(5.3.1a,b), are non-negative. From Eqs.(5.3.1a,b), the exit rates can be decomposed as:

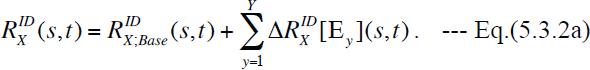

**Figure 12.**
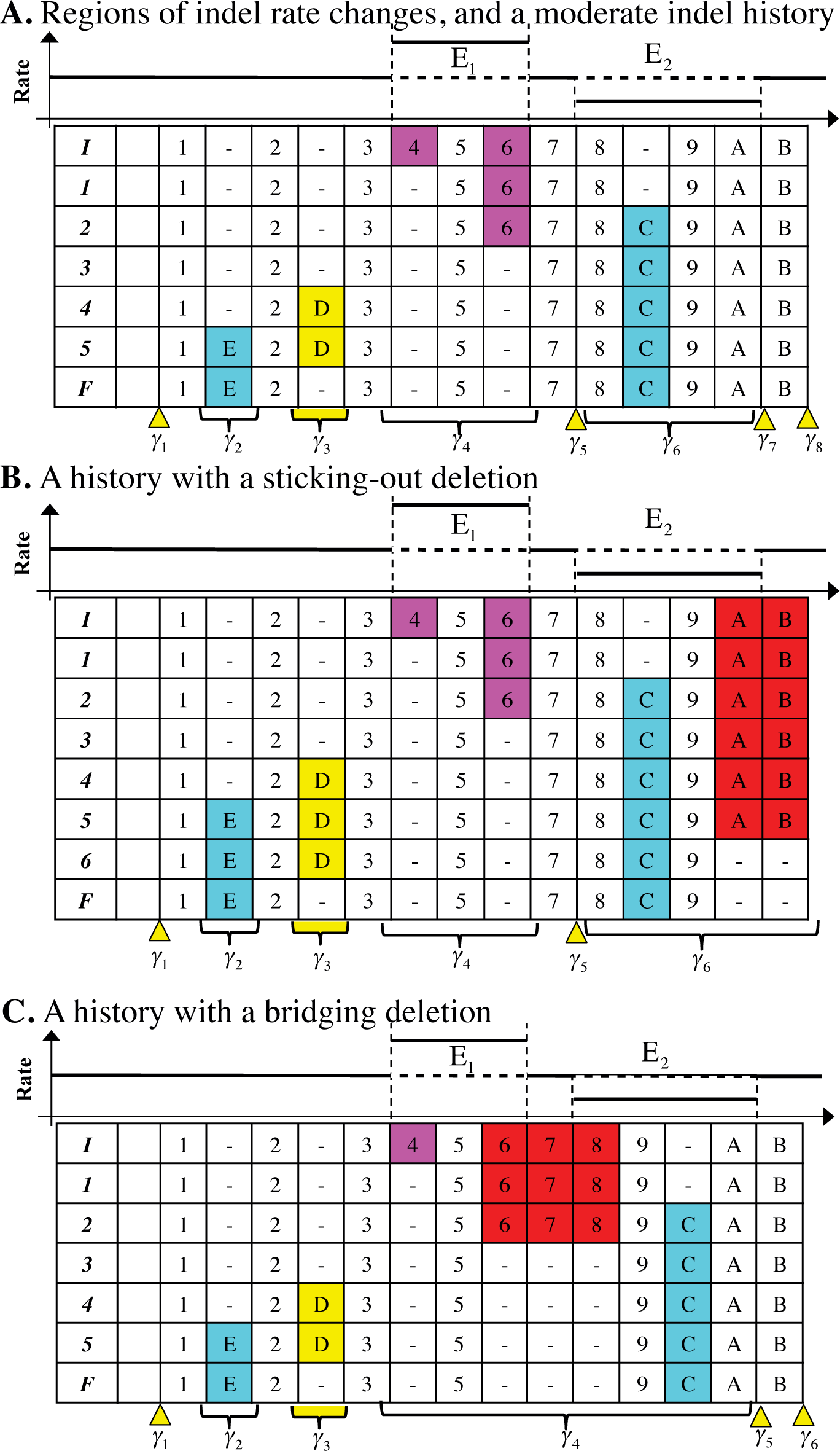
Example of partially factorable indel model, Eqs.(5.3.1a,b). In each panel, the graph above the MSA schematically shows the indel rates of the regions. In the example here, indel rate increments are confined in two regions, *E*_1_ and *E*_2_. Other than that, the figure uses the same notation as in Figure 10. **A.** When all indels are either completely within or completely outside of the regions. Although the deletion of a site with the ancestry ‘4’ and the deletion of a site with the ancestry ’6’ are separated via a preserved ancestral site (with the ancestry ‘5’), they are lumped together into a single local indel history, because they are both within the region *E*_1_. **B.** When a deletion sticks out of the region of an indel rate increment. The deletion of the two sites (with the ancestries ‘A’ and ‘B’) sticks out of the region *E*_2_. In this case, γ_6_ is extended to encompass this deletion, and end up engulfing the original γ_7_ and γ_8_ (in panel A). All indel events within this extended γ_6_ define a single local indel history. **C.** When a deletion bridges the two regions of indel rate increments. The deletion of the three sites (with the ancestries ‘6’, ‘7’ and ‘8’) bridges the regions *E*_1_ and *E*_2_. In this case, the regions *E*_1_ and *E*_2_, as well as the spacer region between them, are put together to form a “meta-region,” which in turn determines a revised γ_4_, and the indel events within it are lumped together to form a single local indel history.

Here,

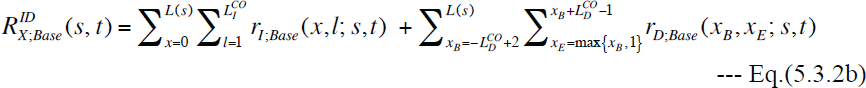

is the baseline exit rate. And

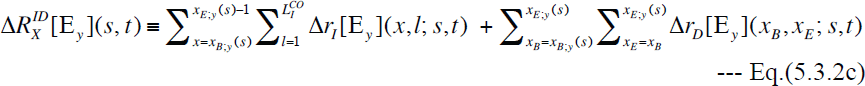

is the increment of the exit rate confined in, and dependent only on, the region E*_y_* (*s*) (*y* = 1, …, *Y*). As explained at the bottom of Subsection 5.1, 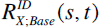 alone gives factorable alignment probabilities. And the increments, 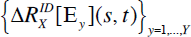, behave independently of each other, as well as of the portions of sequence states in the remaining regions. Thus, similarly to the final model in Subsection 5.2 (called the “multi-conservation model” here),if each indel event is completely confined in any of the E*_y_* (*s*)’s or in any spacer regions between neighboring E*_y_* (*s*)’s (Figure 12, panel A), the alignment probability can be expressed as the product of the overall factor and the contributions from all the E*_y_* (*s*)’s and all the local indel histories within spacer regions. And, also similarly to the multi-conservation model, even if some events within a region E*_y_* (*s*) are separated from the others by at least a PAS, they must be put together into asingle local indel history (panel A). One major difference from the multi-conservation model is that the current model allows deletions to stick out of a region (E*_y_*(*s*))or even bridge between two or more regions (panels B and C). The rates of such deletions and indels that are completely outside of the regions are given by the baseline rates. When a deletion sticks out of a region, the region will be extended to encompass the deletion, and all events within the extended region are lumped into a single local indel history (panel B). When a deletion bridges between two or more regions, a “meta-region” encompassing all bridged regions is defined, and all events within the meta-region will form a single local indel history (panel C). In contrast, the indels completely outside of the regions should be independent of each other as long as they are separated by at least a PAS. Hence, under this model, the PWA probabilities are “factorable” in this somewhat non-trivial sense.

In Appendix A6, we proved that, under a space-homogeneous continuous-time Markov model of indels, the total probability of each LHS equivalence class of indel histories (during a time interval)calculated via the method of Miklós et al. (2004) is identical to that calculated via our *ab initio* formulation. Although we will not explicitly prove here, we believe that the proof can be extended to the indel model given in this subsection as well, if we re-define a “chop-zone” as a region that can potentially accommodate a local indel history (as defined here)plus its right-flanking PAS.

## Discussion

Here we will discuss some issues we did not elaborate on in Results or Appendix.

In this study, we only considered simple boundary conditions. Each sequence end was either freely mutable or flanked by a biologically essential region that allows no indels. These boundary conditions may remain good approximations if the subject sequences were extracted from well-characterized genomic regions. In real sequence analyses, however, the situations are unlikely to be so simple. This is because the ends of the aligned sequences are often determined by artificial factors, such as the methods to sequence the genome, to detect sequence homology, and to annotate the sequences. Moreover, the constant cutoff lengths (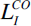 and 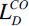) were introduced just for the sake of simplicity, to broadly take account of the effects of various factors that suppress very long indels (such as selection, chromosome size, genome stability, etc.). In reality, it is much more likely that the cutoff lengths would vary across regions. Then, the alignment probabilities would be only approximately factorable, as in the second example model discussed in Subsection 5.2 of Results. In order to pursue further biological realism and to enable further accurate sequence analyses, it would be inevitable to address these issues seriously.

We developed our *ab initio* perturbative formulation aiming to calculate the probabilities of given alignments, especially MSAs, quite accurately, with the ultimate goal of applying it to the reconstruction of a fairly accurate probability distribution of candidate MSAs from an input set of homologous sequences. And, as you will see in parts II and III (Ezawa, Graur and Landan 2015a,b), we actually developed some analytical and computational methods to calculate alignment probabilities via our formulation.

At the same time, however, we strongly caution the readers that, at this point, a naïve application of these methods to a *reconstructed* MSA is fraught with high risks of incorrect predictions of indel histories,*etc*. This is because *reconstructed* MSAs, *even if they were reconstructed via state-of-the-art aligners* (reviewed, *e.g.*, in Notredame 2007), are known to be considerably erroneous (*e.g.*, Löytynoja and Goldman 2008; Landan and Graur 2009). Thus, it would be preferable to first develop a method or a program that accurately assesses and rectifies alignment errors, before using our formulation to make some evolutionary or biological predictions. These topics will be discussed in more details in part III (Ezawa, Graur and Landan 2015b).

When reconstructing the probability distribution of candidate MSAs, quite fast MSA samplers will be necessary. In Appendix A6, we demonstrated that, as far as each LHS equivalence class is concerned, the probability calculated via the method of Miklós et al. (2004) is equal to that calculated via our formulation, at least under their space- and time-homogeneous indel model. Thus,our formulation could use at least the dynamic programming (DP) of Miklós et al.(2004), possibly with some modifications, both to identify the optimum PWA and to sum the probabilities over candidate PWAs. A problem would be that the full version of their DP is quite slow, with the time complexity of *O*(*L*^4^), where *L* represents the sequence length. Although the rough version of their DP is *O*(*L*^2^), we are currently not sure whether it is compatible with biologically realistic situations. Therefore, it would be preferable if we can devise a sampling method that is smarter and more suitable for our formulation. Part III (Ezawa, Graur and Landan 2015b) will also discuss this topic and some other possible applications in more details.

## Conclusions

To the best of our knowledge, this is the first absolutely orthodox study to theoretically dissect the calculation of the probabilities of the alignments (whether they are PWAs or MSAs) *purely from the first principle*, under a *genuine* stochastic evolutionary model, which describes the evolution of an *entire* sequence via insertions and deletions (indels) along the time axis. The model handled here extends the previously most general evolutionary model, *i.e.*, the general form of the “substitution/inserton/deletion models” proposed by Miklós et al.(2004). It should be noted that we did not impose any unnatural restrictions such as the prohibition of overlapping indels. Nor did we make the pre-proof assumption that the probability is factorable into the product of column-wise or block-wise contributions. The only tricks that we took advantage of are the techniques that were essential for the advances of the theoretical physics in the 20th century, namely, the bra-ket notation of state vectors, the operator representation of the actions of indels, and the perturbation expansion (*e.g.*, Dirac 1958; Messiah 1961a, 1961b). We slightly modified them here so that they will be applicable to the finite-time stochastic evolution operator. Using these techniques, we formally showed that the probability of an alignment can indeed be expressed as a summation of the probabilities over all global indel histories consistent with the alignment. Our derivation via the perturbation expansion serves as a bridge between Gillespie’s (1977) intuitive derivation of his own method for stochastic evolutionary simulations and Feller’s (1940) mathematically rigorous theorems underpinning the method. Then, under a most general set of indel rate parameters, we went on to find a sufficient and nearly necessary set of conditions on the indel rate parameters and exit rates under which the alignment probability can be factorized into the product of an overall factor and the contributions from regions separated by gapless columns (or preserved ancestral sites). We also showed that quite a wide variety of indel models could satisfy this set of conditions. Such models include not only the “long indel” model (Miklós et al. 2004) and the indel model of a genuine molecular evolution simulator, Dawg (Cartwright 2005), but also some sorts of models with rate variation across regions. Moreover, we proved that, as far as each LHS equivalence class is concerned, the probability calculated via the method of Miklós et al. (2004) is equivalent to that calculated via our *ab initio* formulation under their spatiotemporally homogeneous indel model.

To summarize, by depending purely on the first principle, this study established firm theoretical grounds on which other approximate indel probabilistic models can be based. And, as will be demonstrated in the subsequent papers (Ezawa, Graur and Landan 2015a,b), it also provides a sound reference point to which other indel models can be compared in order to see when and how well they can approximate the true alignment probabilities.

## Acknowledgements

This study is dedicated to the late Dr. Keiji Kikkawa, who was a renowned theoretical physicist,one of the key pioneers of the string field theory of the elementary particle physics, and the best ever mentor of K.E. We are grateful to Dr. R. A. Cartwright at Arizona State University for his useful information and discussions that inspired this study. We appreciate the logistic support and the feedback of Dr. Tetsushi Yada at the Kyushu Institute of Technology. We would also like to thank the three anonymous referees of the predecessor manuscript entitled: “Framework that enables approximate lilelihood analysis of insertions/deletions on multiple sequence alignment.” Their comments helped drastically improve the study itself, not to mention the manuscript. We would like to thank Dr. Toshiaki Kakitani, a professor emeritus at Nagoya University, for his encouragement and feedback on the manuscript. We are also grateful to Dr. Ian Holmes at University of Califonia, Berkeley, for his critical comments on a different manuscript on this work and useful information on some relevant previous studies. This work was a part of the project,“Error Correction in Multiple Sequence Alignments,” which was funded by US National Library of Medicine [grant number LM010009-01 to Dan Graur and Giddy Landan at the University of Houston]. The later stage of this work was also supported by Grants-in-Aid No. 221S0002, which was awarded to Tetsushi Yada by the Ministry of Education, Culture, Sports, Science and Technology of Japan.

### Authors’ contributions

KE conceived of and mathematically formulated the theoretical framework in this paper, participated in designing the study, performed all the mathematical analyses, and drafted the manuscript. DG and GL participated in designing the study, helped with the interpretation of the data, and helped with the drafting of the manuscript.

## Appendix

**A1. Equivalence relations between products of operators representing overlapping indels**

In Subsection 2.3 of Results, we mainly discussed the equivalence relations between the products of operators representing non-overlapping indels, based on the fundamental binary equivalence relations, Eqs.(2.3.3a-d). Here, we will give some typical equivalence relations involving indels that overlap each other.

First, consider the action of two indels that are spatially nested or adjacent to each other (panel A of Figure 3). Because such actions are indistinguishable from the action of a single deletion, we have:

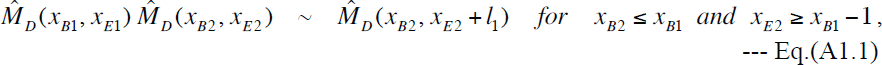

where 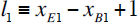. When 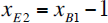 and 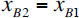, the 2nd event deletes a subsequence on the immediate left and on the immediate right, respectively, of the subsequence deleted by the 1st event.

Second, consider the action of two insertions that are spatially nested or adjacent to each other (panel B). In the state space *S^I^* or *S^III^*, we cannotdistinguish the action from the action of a single insertion. Thus, we have:

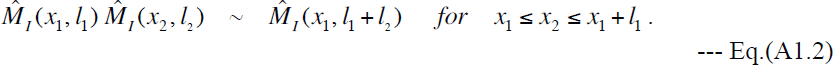

When 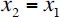 and 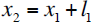, the 2nd event inserts a subsequence on the immediate left and on the immediate right, respectively, of the subsequence inserted by the 1st event. In the state space *S^III^*, however, the left-hand side of Eq.(A1.2) is distinguishable from the right-hand side, or the left-hand side with different *x*_2_ or *l*_2_ (while keeping the same *l*_l_ + *1*_2_) are distinguishable from each other.

Third, consider the action of a deletion and an insertion that overlap with each other. There are several different patterns of such cases. A deletion and a subsequent insertion can overlap or touch each other only when the insertion occurs *exactly* between the sites that flanked the deleted subsequence (Figure 3, panel C). Thus, we can differentiate these cases only through the patterns of an insertion followed by a deletion (panels D, E, F, G and H). There are four possible patterns: (a) cases where the deleted region completely encompasses the inserted subsequence (panel D); (b) cases where the deleted region is completely nested within the inserted subsequence (panel E); (c) cases where the deleted region overlaps the left fragment of the inserted subsequence (panel F); and (d) cases where the deleted region overlaps the right fragment of the inserted subsequence (panel G). The equivalence relations for these cases are as follows:

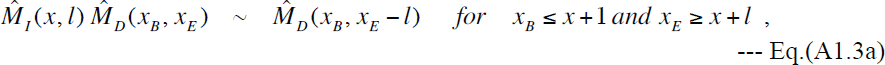

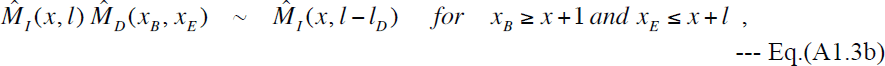

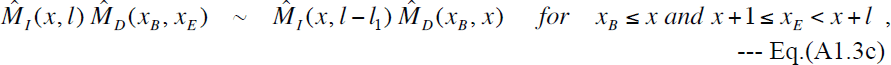

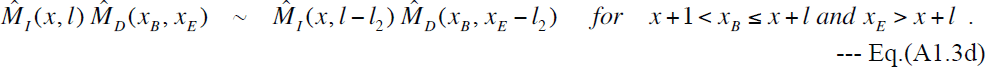

Here 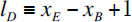, 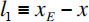, and 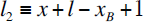.

Eqs.(A1.3a,b) exclude the case with 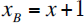 and 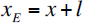, which yields a crucial equivalence relation (Figure 3, panel H):

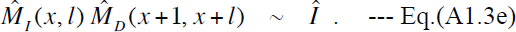

And the right-hand sides of Eqs.(A1.3c,d) are also equivalent to the action of a deletion followed by an insertion exactly between the sites flanking the deleted subsequence (panel C). More precisely,

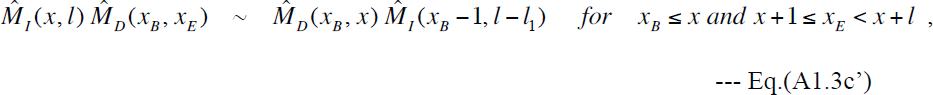

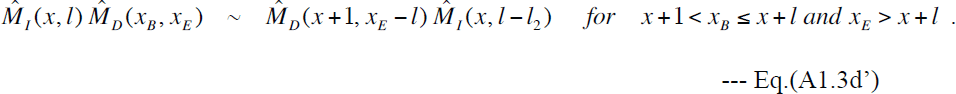

Among these equations, Eqs.(A1.3a,d,e,d’) hold in any state space, *S^I^*, *S^II^*, or *S^III^*, whereas Eqs.(A1.3b,c,c’) hold only in *S^I^* or *S^II^* but not in *S^III^ in its strict sense*. (But Eqs.(A1.3c,c’) could hold also in *S^III^* if the space’s sense is broadened.)

Almost all the equivalence relations between local indel histories on the same region should be derived from serial applications of these equivalence relations, Eq.(A1.1), Eq.(A1.2), and Eqs.(A1.3a-e,c’,d’), possibly supplemented by the binary equivalences, Eqs.(2.3.3a-d), and the unary equivalences, Eqs.(2.3.1a-c).

**A2. “Decomposition” of 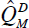, deletion component of rate operator**

Using the unary equivalence relations, Eqs.(2.3.1a,b,c), we can further rewrite the definition of 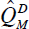, Eq.(2.4.2c’), into a summation of contributions from the deletions in the middle of the sequence (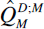), on the left-end (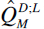), on the right-end (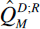), and from the whole-sequence deletions (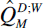), as follows:

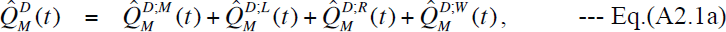

where

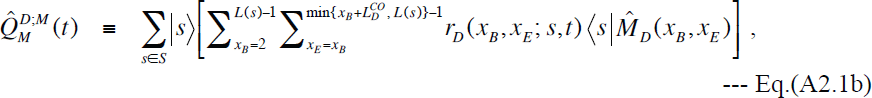

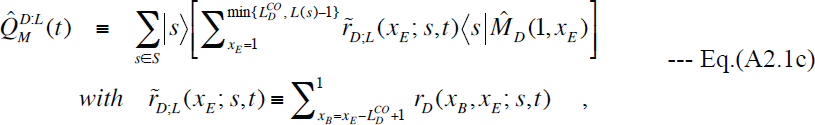

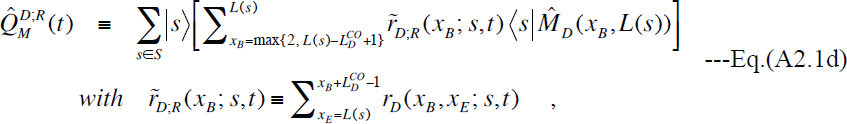

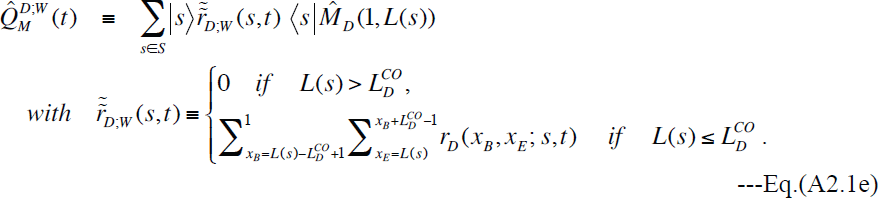

Eqs.(A2.1a-e) could sometimes simplify theoretical thinking and also save computational costs by doing away with deletions that stick out of the boundaries of the sequence under consideration.

**A3. Multiplicativity of perturbation expansion: details**

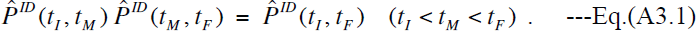

Let’s see how this condition is satisfied by the perturbation expansion, Eq.(3.1.3), that is,

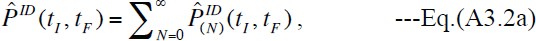

With

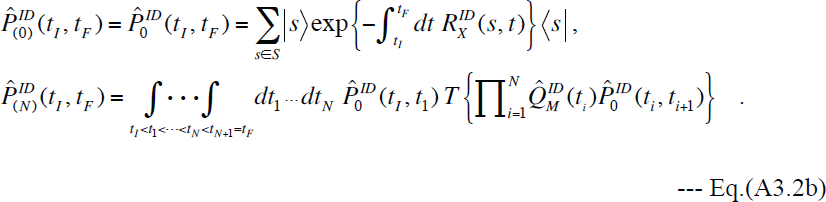

Substituting Eq.(A3.2a) into Eq.(A3.1) and comparing the terms with the same number of indel operators, we find that the following equation must be satisfied for *N* = 0, 1, …:

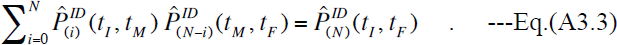

We will prove this by induction. For *N*=0, the equation can be proven easily:

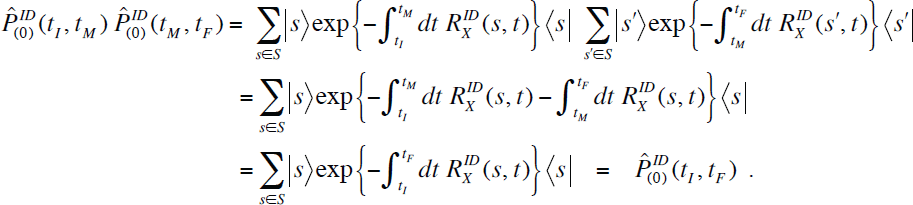

Next, assume that Eq.(A3.3) holds for a particular non-negative integer *N*. The left hand side of Eq.(A3.3) with *N* replaced by *N* + 1 can be rewritten as:

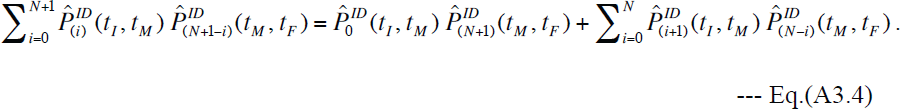

To go further, we first notice that the following equation holds from Eq.(A3.2b):

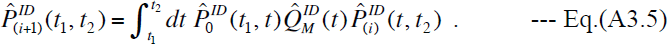

Substituting it into the first term of the right-hand side of Eq.(A3.4), we have:

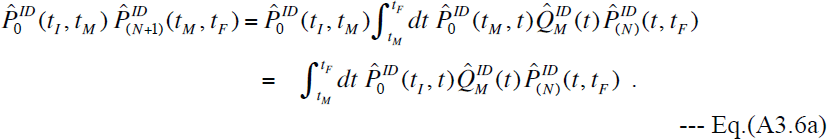

Meanwhile, the second term of the right-hand side of Eq.(A3.4) can be rewritten as:

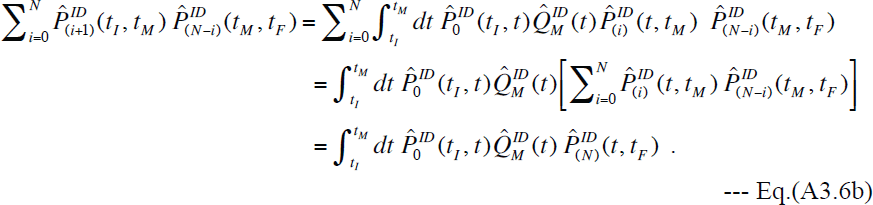

To get the last equation, the assumed Eq.(A3.3) for *N* was used. Summing Eq.(A3.6a) and Eq.(A3.6b), we see that the right-hand side of Eq.(A3.4) becomes:

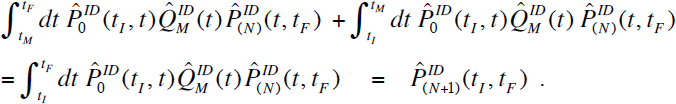

To get the last equation, Eq.(A3.5) for *i* = *N* was used. Thus, if Eq.(A3.4) holds for a particular *N*, then it holds also for *N* + 1. Therefore, Eq.(A3.4)holds for every non;negative integer *N*, which guarantees that our stochastic evolution operator, Eq.(3.1.3), and its more specific representation, Eq.(3.1.8), do indeed satisfy the Chapman-Kolmogorov equation, up to a desired degree in the perturbation expansion. [NOTE ADDED: The proof given in this subsection is essentially equivalent to the proof of Theorem 5 in Feller (1940). In a sense, this subsection could be considered as fully explaining his rather blunt proof (as well as the context the theorem is in) and recasting the explanation into the operator representation. In general, the operator representation gives a clearer intuitive picture than the probability representation used by Feller.]

**A4. Proof of factorization of multiple-time integration, Eq.(4.1.4)**

The identity,Eq.(4.1.4),

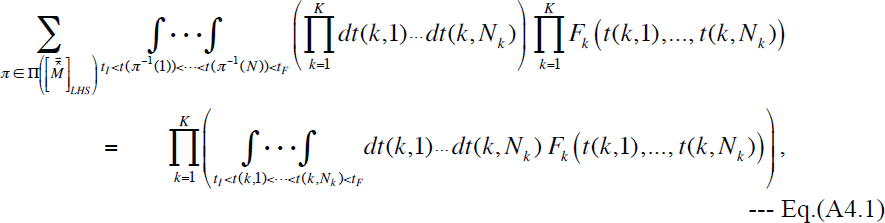

where 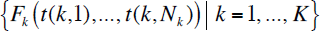 is any set of non-singular functions of multiple-time points, is one of the two essential elements for obtaining our sufficient and nearly necessary set of conditions for the factorability of the PWA probability. The identity states that, if we sum the multiple-time integration operations for global indel histories over a LHS equivalence class, it can be factorized into the product of multiple-time integration operations, each for a local indel history, over the LHS. Here, we prove this identity in a mathematically rigorous manner.

Let us remember here that 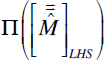 denotes the set of maps that correspond to global indel histories in a LHS equivalence class. Each of its elements is expressed as:

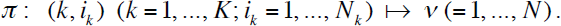

Then, we first note that, because the integrands and the sets of variables of integration are identical on both sides of Eq.(A4.1), proving this identity is equivalent to proving the equality (*modulo differences of measure zero*) between the domains of integration:

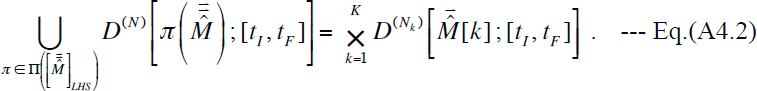

Here, 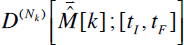 is the domain of integration for the *k* th local indel history, 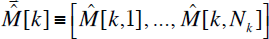:

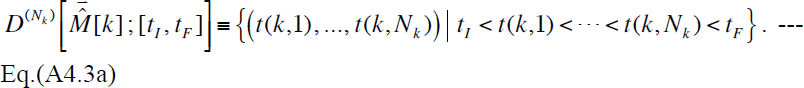

And 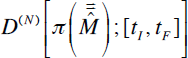 is the domain of integration for the global indel history, 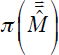:

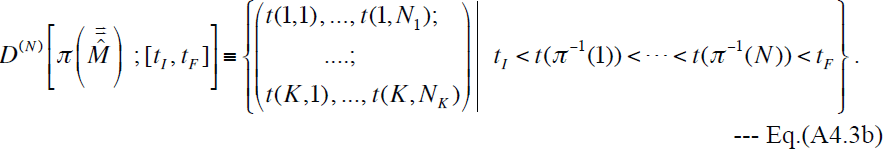

To go further, let us introduce a new notation, 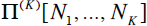, that represents 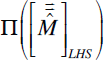 to remind that each of its 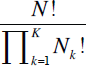 elements can be re-interpreted as a rearrangement of *K* sets, whose sizes are 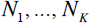, into a single set of size 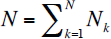 Then, each map 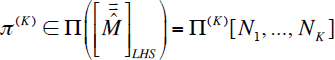 can be re-expressed as a composite map, 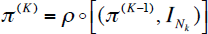. Here 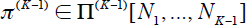 is a rearrangement of *K* ̶ 1 of the original *K* sets excluding the *K* th set, 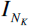 is the identity map from the *N_k_* elements in the *K* th set to themselves, and 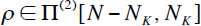 is a rearrangement of the *K* th set and the remainder made from the *K* ̶ 1 sets. The numbers of the elements exactly match,because we have 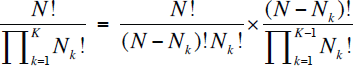. Provided that the binary (*i.e., K* = 2)version of Eq.(A4.2)is proved, then we can apply them for each fixed 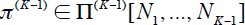 and all 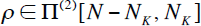, and we can factor out the contribution from the *K* th local (or “separated”) indel history.

This is formally proved as follows. First, the left-hand side of Eq.(A4.2) is re-expressed as:

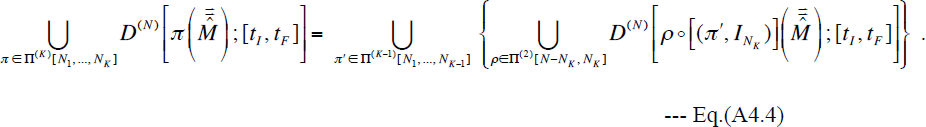

On the right-hand side, we have 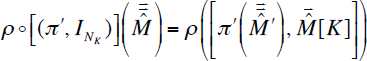 by definition. Here, 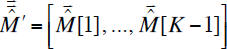 is the “reduced ” LHS consisting of *K* ̶ 1 out of the original *K* local indel histories in 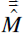, excluding the *K* th local history, 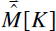. Substituting this into Eq.(A4.4), and assuming that Eq.(A4.2) holds with *K* = 2, we have:

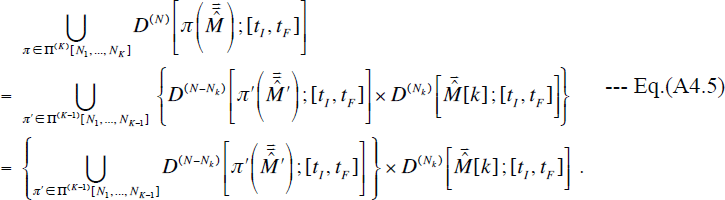

This series of equations re-expresses the above verbal reasoning in clear mathematical terms, and formally demonstrates that the domain of integration for the rightmost local indel history (*i.e.*, the *K* th local history) is indeed factored out. Iteratively applying the above reasoning to the remaining set of *K* ̶ 1 local indel histories, we can prove that the domains of integration for all local indel histories can be factored out. This finally gives Eq.(A4.2) and thus proves the identity, Eq.(A4.1), *i.e.*, Eq.(4.1.4). Thus, the problem at hand was reduced to proving Eq.(A4.2) with *K* = 2, which we will call the “binary domain identity” here. It is rewritten as:

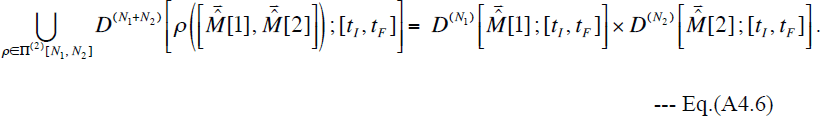

Using Eqs.(A4.3a,b), and setting 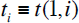 and 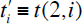 it can be rewritten further as:

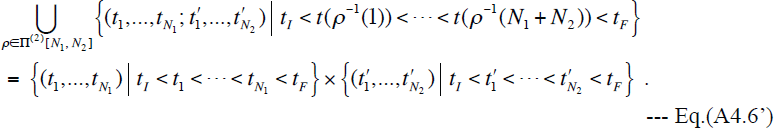

(In this equation and hereafter in this subsection, the identities are considered *modulo differences of measure zero*.)

We will prove this identity, Eq.(A4.6’), *via* mathematical induction regarding *N*_2_. First, we show Eq.(A4.6) with *N*_2_ = 1 holds for every fixed positive integer *N*_1_. In this case, 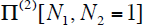 consists of *N*_1_ + 1 elements, each of which inserts the event in the 2nd local history between the *i* th and *i* + 1 th events in the 1st local history 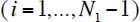, or places it before or after all events in the 1st local history.

Thus, we have:

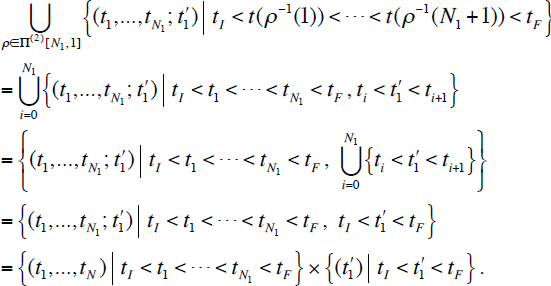

Here we set 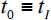 and 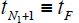. This shows that Eq.(A4.6’) with *N*_2_ = 1 holds for every 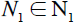.

Next, let us assume that the binary domain identity, Eq.(A4.6’), holds for 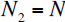 and for every 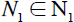 (N_1_ is the set of positive integers),and see if the identity also holds for 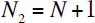. For this purpose, we classify 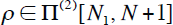 according to the position of 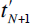 relative to 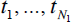, and let 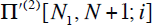 (with 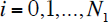) be the subset of 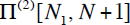 whose elements satisfy 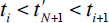. Here we set 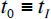 and 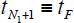 again. For every 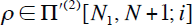, there exist a unique 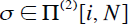 such that: 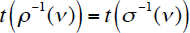 for 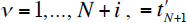 for 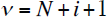, and 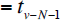 for 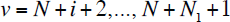. It could also be represented as:

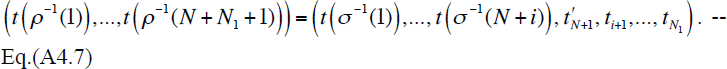

Thus, 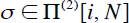 corresponds to the local sub-history before 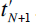. Taking advantage of these facts, the left-hand side of Eq.(A4.6’) with 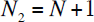 is re-expressed as:

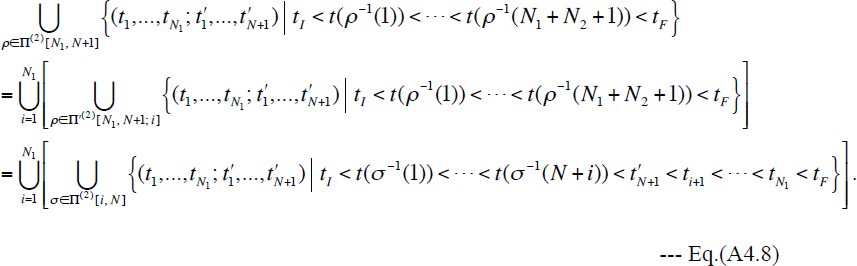

Applying the assumed Eq.(A4.6’) with *N*_2_ = *N* and *N*_1_= *i*, and with *t_F_* replaced by 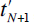, to the expression in the square brackets on the rightmost hand side of Eq.(A4.8), we have:

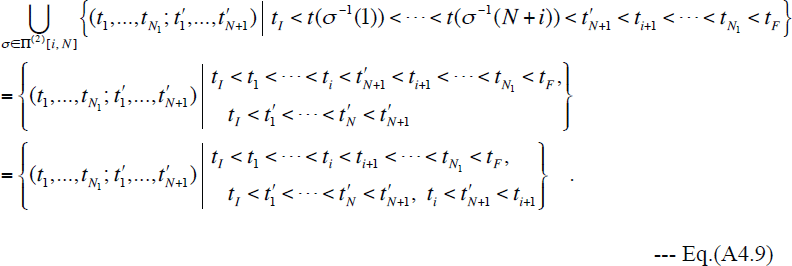

Substituting Eq.(A4.9) back into the rightmost hand side of Eq.(A4.8), we finally get:

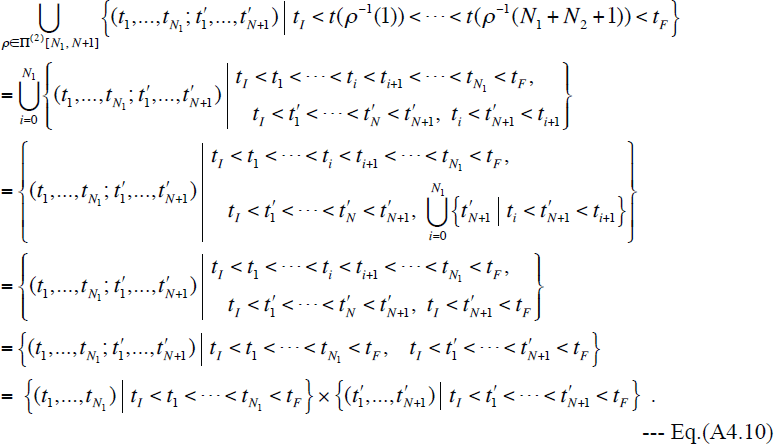

This final expression is nothing other than the right-hand side of Eq.(A4.6’) with *N*_2_ = *N* + 1. Thus, assuming that Eq.(A4.6’) holds for *N*_2_ = *N* and for every 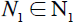, we did indeed show that it also holds for *N*_2_ = *N* + 1 and for every 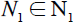. Therefore, the binary domain identity, Eq.(A4.6’), holds for every pair, 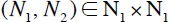. This completes the proof of our key identity, Eq.(A4.2), and therefore the proof of the factorization of the multiple-time integration, Eq.(A4.1).

**A5. Proof of proposition 4.1.1 for factorization of exponent**

The other core element is the proposition 4.1.1:

“Let 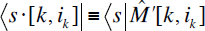 and 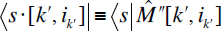 be the states resulting from the actions of the equivalents of events 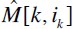 and 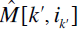 respectively, on 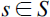. And let 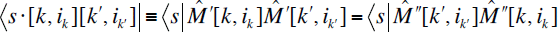 be the state resulting from the consecutive actions of the equivalents of 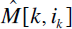 and 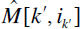 on *s*. The equation for the exponents, Eq.(4.1.3’b), holds for every global history 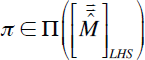 and for each of its sub-histories that could occur in any sub-interval, [*t_I_*, *t*] with 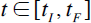 if and only if the equation,

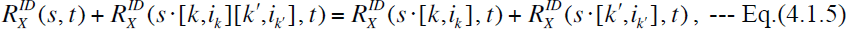

holds for every pair, 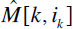 and 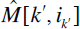 (with 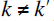), in the LHS 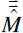, for every possible state 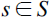 before the equivalents of 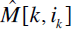 and 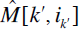 in the global histories in 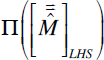 and at any time 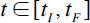.”

It provides an essential part of our sufficient and nearly necessary set of conditions for the factorability of the PWA probability. Here, we prove this proposition via mathematical induction, similarly to the proof in Appendix A4.

We first reduce the problem into a binary one by mathematical induction regarding the number of local indel histories, *K*. As in Appendix A4, let 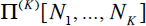 denote the set of maps, 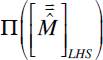, each of whose elements is a rearrangement of *K* sets, of sizes 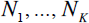, into a single set of size 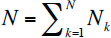. And re-express each map 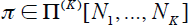 as a composite map, 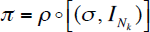 where 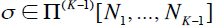 and 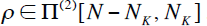. Then, also as in Appendix A4, we have 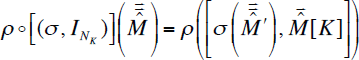 by definition, where 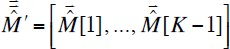 is the reduced LHS consisting of *K* −1 out of the original *K* local indel histories in the LHS 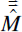, excluding 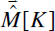. Thus, if the binary version of the proposition 4.1.1, with 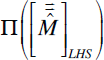 replaced by 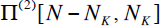, is true for each fixed 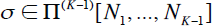, we have the binary version of the factorization, Eq.(4.1.3’b):

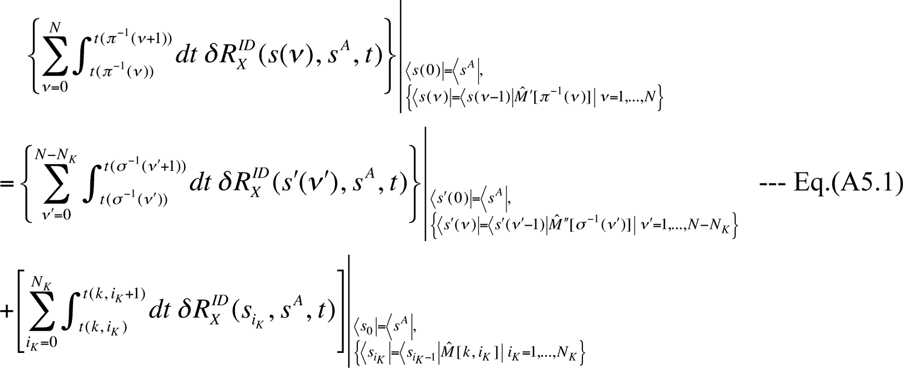

for every possible 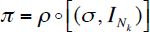 with the fixed σ and any 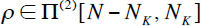. The first summation on the right-hand side is the left-hand side of Eq.(4.1.3’b) with 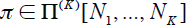 replaced by 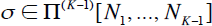. Thus, the problem was reduced to that of the factorization for the global indel histories equivalent to a set of *K* -1 local indel histories. By iteratively applying the binary version of the proposition 4.1.1 to the reduced problems, we will finally obtain the fully factorized form, *i.e.*, the right-hand side of Eq.(4.1.3’b).

Therefore, all we have to do is to prove the binary version of the proposition 4.1.1. To do so, we will rewrite it into a more tractable form. We first pick two integers, 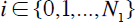 and 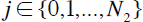, and consider all sub-histories of indels composed of two local sub-histories, 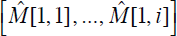 and 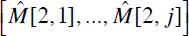. (If *i* = 0 or *j* = 0, the corresponding local sub-history is considered as empty.) Each such sub-history corresponds to a map, 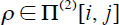 and the state resulting from the action of this sub-history on the state 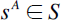 is represented, *e.g.*, as: 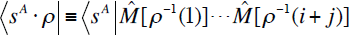. As in Subsection 2.3, through the binary equivalence relations, Eq.(2.3.3a-d), we can show that 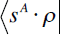 for each sub-history 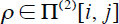 is in fact equal to the state:

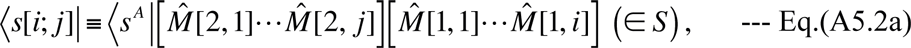

that is uniquely determined solely by the local sub-histories, 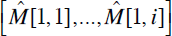 and 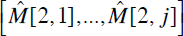, and the initial state, 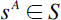. That is, the state 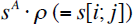 is independent of further details of 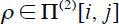. (Naturally, we have 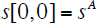.) Thus, the binary version of the proposition 4.1.1 is rephrased as follows.

**[Proposition A5.1]**

“Eq.(4.1.3’b) with *K* = 2 holds true for 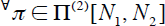 and for each of their sub-histories during [*t_I_*, *t*] with 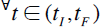 if and only if the equation,

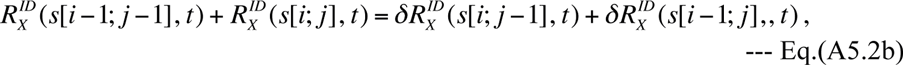

holds for 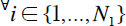, 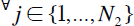, and for 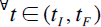.”

Here comes the proof of the proposition A5.1. First of all, we rewrite Eq. (A5.2b) in two different ways, as:

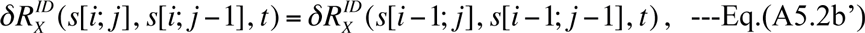

and

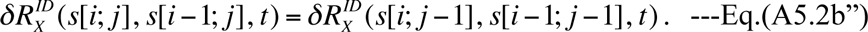

These equations collectively indicate that the increment of the exit rate due to an indel event in one local indel history will not be influenced by the past events in the other local history. Indeed, these equations can be “solved” to give:

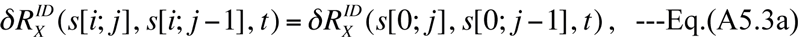

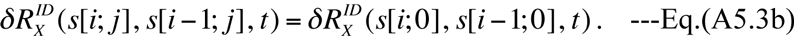

The right-hand sides of Eq.(A5.3a) and Eq.(A5.3b) are, respectively, the increment purely within the 2nd local history and that purely within the 1st local history.

Replacing *i* with *i′* in Eq.(A5.3b), and summing the result over 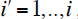, we find:

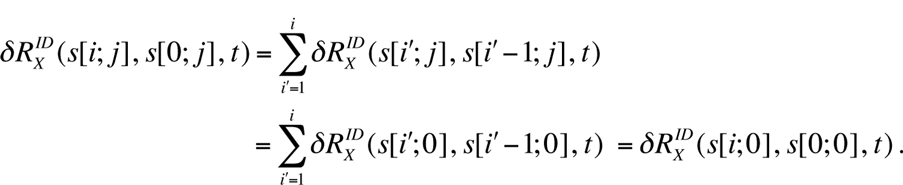

Using 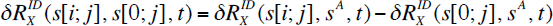 and 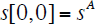, we get a key equation:

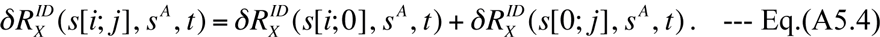

This means that the increment of the exit rate by a sub-history 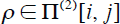 is decomposed as the summation of two increments, each by one of the local sub-histories, 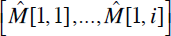 and 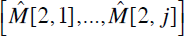.

Now, pick an indel history corresponding to a map 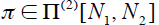, and consider the left-hand side of Eq.(A1.3’b) with *K* = 2, *i.e.*, 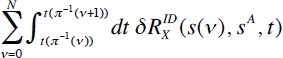 with 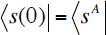 and 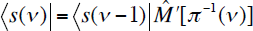 for 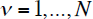. Let 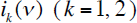 be the number of events in the local history 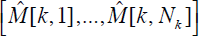 that are equivalent to those included in the sub-history 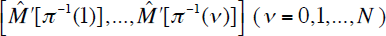. Then, we have 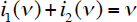, and 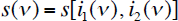. Thus, using Eq.(A5.4), the left-hand side of Eq.(A1.3’b) with *K* = 2 can be decomposed into the contributions from two local sub-histories:

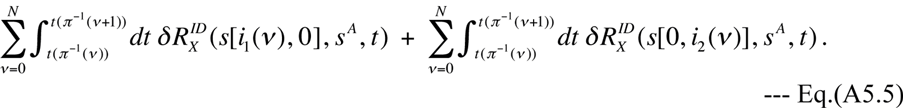

In each summation, *i_k_*(ν) remains a particular value, *e.g.*, *i_k_*, since 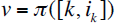 until (and excluding) 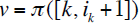 (for *k* = 1, 2). Taking account of it, Eq.(A5.5) becomes:

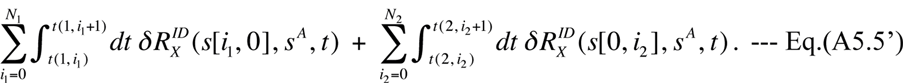

From the definition of *s*[*i*; *j*], Eq.(A5.2a), we can see that Eq.(A5.5’) is nothing other than the right-hand side of Eq.(4.1.3’b) with *K* = 2. The argument after Eq.(A5.4) applies to every history corresponding to 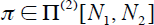. Thus, we proved that Eq.(4.1.3’b) with *K* = 2 holds if Eq.(A5.2b) holds.

To prove the converse, we now assume that Eq.(4.1.3’b) with *K* = 2 holds for the indel history corresponding to every 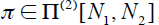, as well as for each of its sub-histories during [*t_I_*, *t*] with 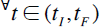. Then, taking the time-derivative of both sides of Eq.(4.1.3’b) with *K* = 2 for any incomplete time-interval [*t_I_*, *t*], we have, for a particular 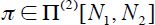: 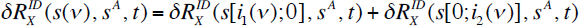, using the 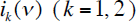 defined above. Because this equation holds for any time-interval 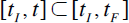 and for every map 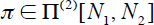, we get exactly Eq.(A5.4) for 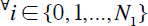, 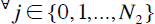, and for 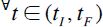. Then it is easy to show Eq.(A5.2b). Starting with the right-hand side of Eq.(A5.2b), we find

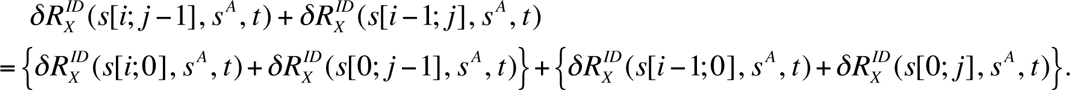

Swapping the 1st and 3rd terms on the right-hand side, we have:

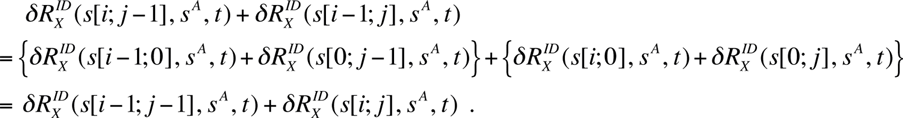

Adding 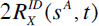 to the leftmost and rightmost sides of the above equation, we get Eq.(A5.2b). Thus, the converse was proved.

This proof of the proposition A5.1, combined with the proof above it resorting to the mathematical induction regarding *K* given the proposition A5.1, completes the proof of the key proposition 4.1.1.

**A6. Probability of LHS equivalence class under “long indel” model**

Here, we consider the “ong indel” model (Miklós et al. 2004), whose indel rate parameters are given by Eqs.(2.4.5a-e). Under this model, we will calculate the probability of a LHS equivalence class of (global) indel histories, conditioned on a given ancestral sequence, according to the prescription proposed by Miklós et al. (2004). And we will show that the probability calculated this way is indeed identical to that calculated via our theoretical formulation.

We first briefly review the method of Miklós et al. (2004). In their method, a PWA is scanned from left to right, and horizontally partitioned into “chop-zones.” In the bulk of the PWA, a chop-zone starts immediately on the right of a preserved ancestral site (PAS) and ends exactly at the next PAS. The leftmost chop-zone starts at the left-end of the PWA and ends exactly at the first PAS if at all, or otherwise ends at the right-end of the PWA. The rightmost chop-zone starts immediately on the right of the rightmost PAS, if at all, and ends at the right-end of the PWA. It should be noted that each chop-zone contains at most one PAS, and that the PAS contained in the chop-zone always resides at the right-end of the zone.

Conceptually, the conditional probability of each chop-zone is calculated by summing the contributions of all local indel histories consistent with the homology structure (Lunter et al. 2005) of the chop-zone. Then, according to the recipe of Miklós et al. (2004), (the indel component of) the probability of a given PWA, conditioned on the ancestral sequence, is given by the product of the conditional probabilities over all chop-zones that make up the PWA. Therefore, by extension, Miklós et al.’s probability of a LHS equivalence class of indel histories (consistent with the PWA) should be given by the product of the contributions from the local indel histories (including the empty histories), each confined in every chop-zone, over all chop-zones constituting the PWA. This is exactly what we will calculate in the following.

Now, as in Subsection 4.1 of Results, consider a LHS equivalence class, 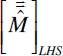 with 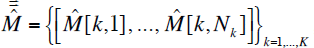, that is consistent with a given PWA, 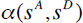, of an ancestral sequence (*s^A^*) and its descendant (*s^D^*). As near the bottom of Subsection 4.1, we can define the regions of 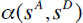 each of which potentially accommodates a local indel history, namely, 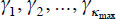, as the region on the left of the leftmost PAS, the regions between two PASs next to each other, and the region on the right of the rightmost PAS. (Because the indel model at hand is space-homogeneous and has freely mutable flanking regions, every local indel history in each such region is independent of the histories outside, both physically and regarding the multiplication factor, as shown in Subsection 5.1 of Results.) Then, by appropriately distributing the local histories into such regions, we can provide a vector-representation of the LHS: 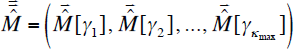. Using these regions, each chop-zone of Miklós et al. (2004) can be constructed by putting together a region 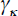 with its right-flanking PAS (for 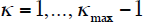), or by a region alone (for 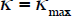). According to Appendix A of Miklós et al. (2004), the contribution from the local history, 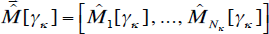, in the chop-zone, 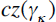, that is associated with γ_κ_ is calculated as:

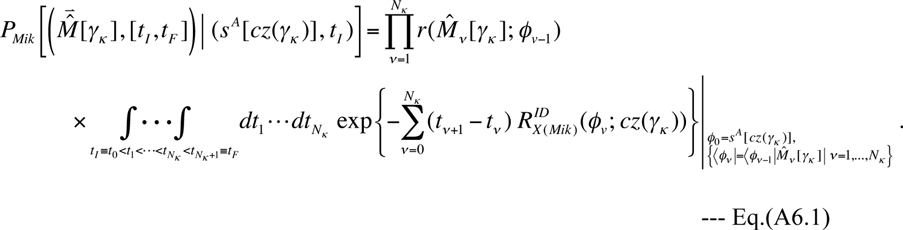

Here, 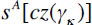 is the portion of the ancestral state confined in the chop-zone 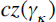, and 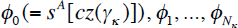 are the chop-zone-confined states that the local indel history went through. The expression is quite similar to each term in the perturbation expansion, Eq.(3.1.8b). Because each indel rate, 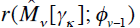, is independent of time, it was put outside of the multiple-time integration. And, because each “exit rate,” 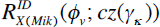 (detailed later), is also time-independent, its time integration (in the exponent) was reduced to a simple multiplication by the time-lapse. The “exit rate” 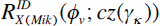 needs some explanation. Because each chop zone (except 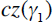) is defined conditionally on the PAS that is left-flanking the zone, and *because we now know that the probability is factorable*, we do not have to consider deletions that pierce through this PAS. Neither do we have to consider indel events completely outside of the chop zone. Therefore, taking advantage of the space-homogeneity of the indel rates, using the correspondence with Dawn’s model (Cartwright 2005), Eqs.(2.4.7a,b,c,d), and letting 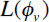 be the number of sites in the state 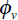 (including the PAS at the right-end of the zone, if at all), the “exit rate” 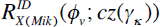 according to Miklos et al.’s definition is expressed as:

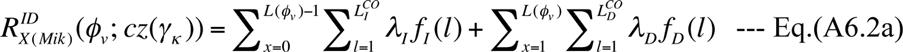

for 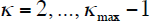. It should be noted that the summation over the insertion positions (*x*) has the upper bound 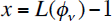, because an insertion on the immediate right of 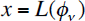 belongs to the right-neighboring chop-zone 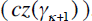. The summation over the indel lengths (*l*’s) is easily performed, and we get:

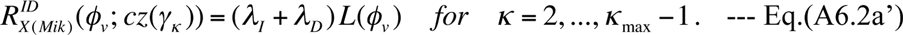

When 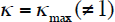, the expression of 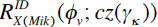 is almost the same as Eq.(A6.2a); the only difference is that it also needs to include the insertions right-flanking the PWA (i.e., with 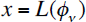), whose rates are given by Eq.(2.4.6c). Thus, we have:

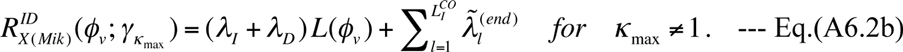

When 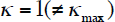, Eq.(A6.2a) is still useful, but we need two modifications, both because this chop-zone is not left-flanked by a PAS. First, insertions on the left-end (*i.e.*, with *x* = 0) must have the rates given by Eq.(2.4.6c). Second, deletions “starting” at *x* = 1 must have the rates 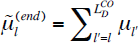. Taking account of these modifications, we have:

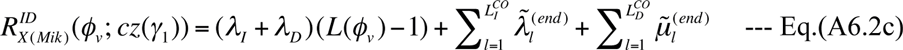

when 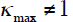. Because 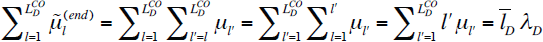, we get:

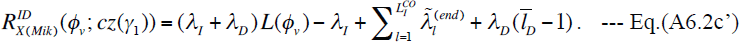

From Eqs.(A6.2a,b,c’), we find that 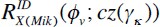‘s are always affine functions of 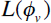 with the slope 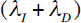, which is the same as that of the exit rate, 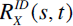 given by Eq.(2.4.7e), for the evolution of an *entire* sequence under the “long indel” model. Thus, we have:

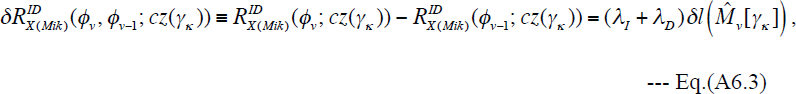

where 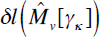 is the change in 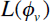 caused by the event 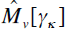. This is exactly the same as the increment of the (actually time-independent) exit-rate:

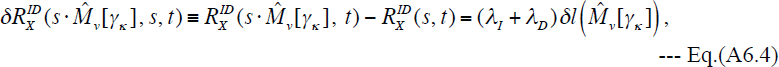

caused by the event 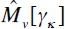 on the entire sequence. By successively applying 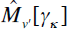 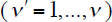, we have:

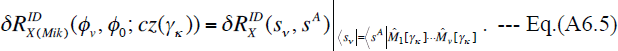

Therefore, we can rewrite the exponent in Eq.(A6.1) as:

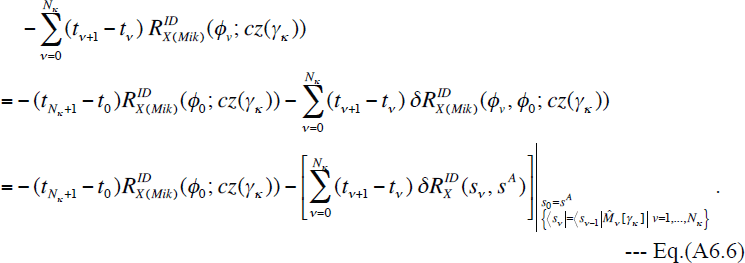

Substituting this back into the right hand side of Eq.(A6.1), and comparing the result with Eq.(4.1.1b) supplemented by Eq.(3.1.8b), we have:

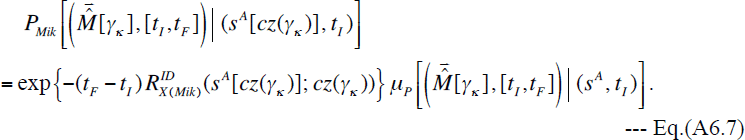

According to the method of Miklós et al. (2004), the probability of the LHS equivalence class, 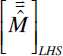 with 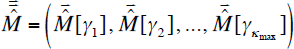, should be *defined* as:

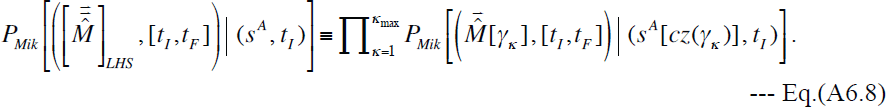

Substituting Eq.(A6.7) into Eq.(A6.8) yields:

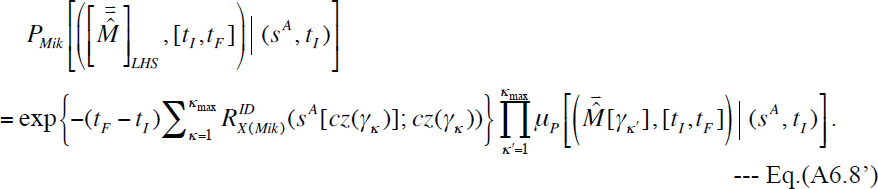

Substituting Eqs.(A6.2a’,b,c’) into the summation in the exponent on the right hand side, we get:

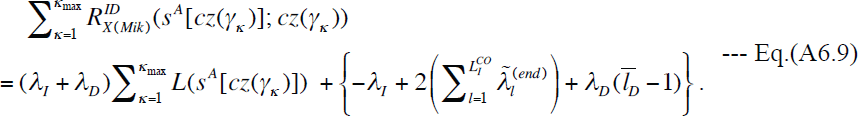

On the right hand side, the expression in the braces is exactly 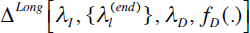 in Eq.(2.4.7e), and we also have 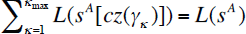. Thus, the equation is further reduced to:

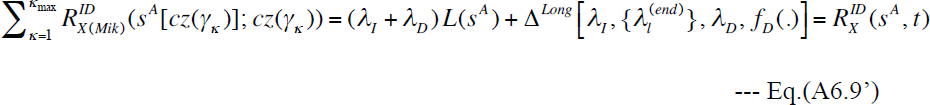

for 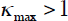. [In the case where 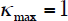, by the way, arguments similar to those leading to Eqs.(A6.2b,c’) reveals that 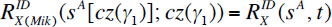 holds, and thus that Eq.(A6.9’) trivially holds.] Now, substituting Eq.(A6.9’) back into Eq.(A6.8’) while taking account of its the time-independence of 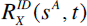 under this model, we finally get:

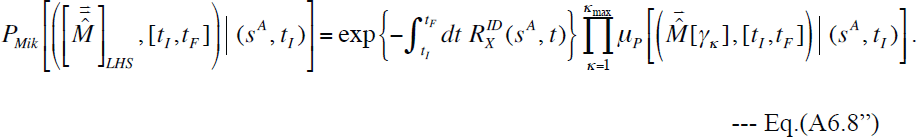

The right hand side of Eq.(A6.8”) is exactly that of Eq.(4.1.7), *i.e.*, the probability of the LHS equivalence class 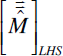 calculated via our *ab initio* theoretical formulation, under the “long indel” model, Eqs.(2.4.7a-e).

Actually, this equivalence between the probability via our *ab initio* formulation and that via Miklos et al.’s method (2004) should hold under any indel models with factorable PWA probabilities described in Section 5 of Results, as long as the “chop-zones” are re-defined appropriately. Its explicit proof will be left as an exercise for the readers. (The key is the decomposition of the entire exit rate into the contributions from (modified) chop-zones.)

**A7. Derivation of Eq.(5.2.6) for “difference between differences” of exit rate of neutral region flanked by completely conserved regions**

Here we derive Eq.(5.2.6), which explicitly expresses the “difference between differences” of the exit rate in the model where a neutrally evolving region is flanked by biologically essential regions or sites. Remember that we are considering a sequence state 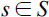 with 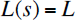, and the action of two separated deletions, 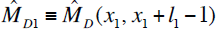 and 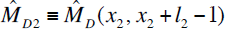 with 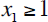 and 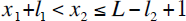, on the state. And we use the notations, 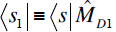, 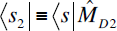, and 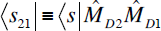. Then, substituting 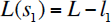, 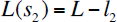, and 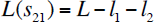 into Eq.(5.2.5), we have:

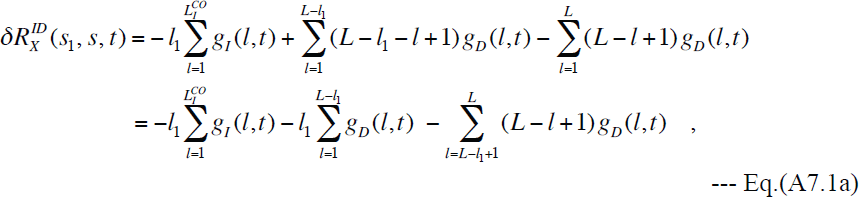

and

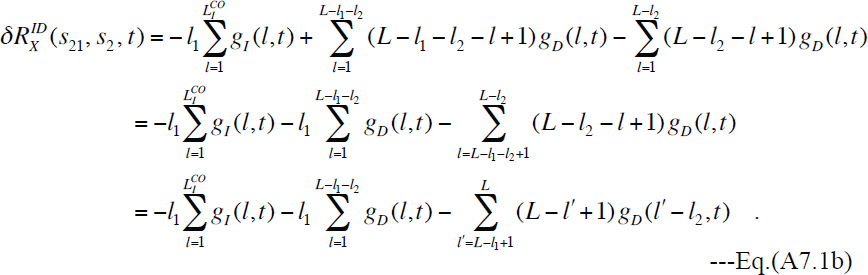

Subtracting Eq.(A7.1b) from Eq.(A7.1a), we get:

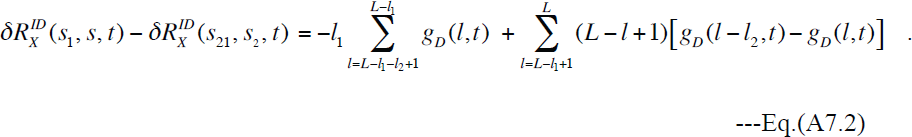

This is exactly Eq.(5.2.6).

## List of abbreviations

CK, Chapman-Kolmogorov; HMM, hidden Markov model; indel, insertion/deletion; LHS, local history set; MSA, multiple sequence alignment; PAS, preserved ancestral site; PWA, pairwise alignment.

